## Supplementary Tables 1-5 for "Enhancing autophagy by redox regulation extends lifespan in *Drosophila*"

**Supplementary Table 1 | Summary of survival data**
Lennicke *et al.*

**Fig. 1a**

| Background | Sex | Genotype | Flies per vial | # vials | Total flies (set-up) | Deaths | Censors | Total flies (actual) | Median (d) | ΔMedian (%) | |
|---|---|---|---|---|---|---|---|---|---|---|---|
| $w^{Dah}$ | female | UAS-cat/+ | 10 | 20 | 200 | 186 | 12 | 198 | 65.3 | 0 | *a* |
| | | da-GAL4/+ | 10 | 20 | 200 | 189 | 11 | 200 | 64.6 | -1.07 | *b* |
| | | da-GAL4>UAS-cat | 10 | 20 | 200 | 159 | 39 | 198 | 69.3 | +6.13 | *c* |
| $w^{Dah}$ | male | UAS-cat/+ | 10 | 20 | 200 | 122 | 75 | 197 | 52.1 | 0 | *d* |
| | | da-GAL4/+ | 10 | 20 | 200 | 145 | 55 | 200 | 51.8 | -0.58 | *e* |
| | | da-GAL4>UAS-cat | 10 | 20 | 200 | 127 | 67 | 194 | 52.8 | +1.34 | *f* |

p-value (Log-Rank test)

| | *b* | *c* |
|---|---|---|
| *a* | 0.7504 | 5.7E-07 |
| *b* | | 3.3E-06 |

| | *e* | *f* |
|---|---|---|
| *d* | 0.5654 | 0.0779 |
| *e* | | 0.1925 |

**Fig. 1b**

| Background/ Sex | Genotype | Condition | Flies per vial | # vials | Total flies (set-up) | Deaths | Censors | Total flies (actual) | Median (d) | ΔMedian (%) | |
|---|---|---|---|---|---|---|---|---|---|---|---|
| $w^{Dah}$ female | da-GS/+ | –RU (0 µM) | 15 | 15 | 225 | 215 | 2 | 217 | 77.3 | 0 | *a* |
| | | +RU (200 µM) | 15 | 15 | 225 | 224 | 1 | 225 | 76.9 | -0.52 | *b* |
| | da-GS> UAS-cat | –RU (0 µM) | 15 | 20 | 300 | 295 | 3 | 298 | 77.8 | +0.65 | *c* |
| | | +RU (200 µM) | 15 | 20 | 300 | 285 | 1 | 286 | 85.1 | +10.09 | *d* |

p-value (Log-Rank test)

| | *b* | *c* | *d* |
|---|---|---|---|
| *a* | 0.7161 | 0.0737 | 1.3E-21 |
| *b* | | 0.1751 | 1.0E-18 |
| *c* | | | 1.1E-16 |

**Fig. 1f**

| Background/ Sex | Condition | Genotype | Flies per vial | # vials | Total flies (set-up) | Deaths | Censors | Total flies (actual) | Median (d) | ΔMedian (%) | |
|---|---|---|---|---|---|---|---|---|---|---|---|
| $w^{Dah}$ female | 5% $H_2O_2$ | UAS-cat/+ | 15 | 8 | 120 | 120 | 0 | 120 | 3.150 | 0 | *a* |
| | | da-GAL4/+ | 15 | 8 | 120 | 114 | 6 | 120 | 3.325 | +5.56 | *b* |
| | | da-GAL4>UAS-cat | 15 | 8 | 120 | 120 | 0 | 120 | 12.200 | +287.30 | *c* |
| $w^{Dah}$ male | 5% $H_2O_2$ | UAS-cat/+ | 15 | 7 | 105 | 105 | 0 | 105 | 3.975 | 0 | *d* |
| | | da-GAL4/+ | 15 | 7 | 105 | 105 | 0 | 105 | 3.825 | -3.77 | *e* |
| | | da-GAL4>UAS-cat | 15 | 5 | 75 | 74 | 1 | 75 | 6.625 | +66.67 | *f* |

p-value (Log-Rank test)

| | *b* | *c* |
|---|---|---|
| *a* | 0.0164 | 5.0E-61 |
| *b* | | 4.9E-62 |

| | *e* | *f* |
|---|---|---|
| *d* | 0.8243 | 3.2E-33 |
| *e* | | 4.6E-33 |

**Fig. 1g**

| Background/ Sex | Condition | Genotype | Flies per vial | # vials | Total flies (set-up) | Deaths | Censors | Total flies (actual) | Median (d) | ΔMedian (%) | |
|---|---|---|---|---|---|---|---|---|---|---|---|
| $w^{Dah}$ female | 20 mM paraquat | UAS-cat/+ | 10 | 10 | 100 | 100 | 0 | 100 | 0.925 | 0 | *a* |
| | | da-GAL4>UAS-cat | 10 | 10 | 100 | 99 | 1 | 100 | 3.550 | +283.78 | *b* |
| $w^{Dah}$ male | 20 mM paraquat | UAS-cat/+ | 10 | 10 | 100 | 100 | 0 | 100 | 3.625 | 0 | *c* |
| | | da-GAL4>UAS-cat | 10 | 10 | 100 | 100 | 0 | 100 | 6.050 | +66.90 | *d* |

p-value (Log-Rank test)

| | *b* |
|---|---|
| *a* | 6.9E-31 |

| | *d* |
|---|---|
| *c* | 8.0E-16 |

**Fig. 1i**

| Background/ Sex | Condition | Genotype | Flies per vial | # vials | Total flies (set-up) | Deaths | Censors | Total flies (actual) | Median (d) | ΔMedian (%) | |
|---|---|---|---|---|---|---|---|---|---|---|---|
| $w^{Dah}$ female | 90% $O_2$ | UAS-cat/+ | 10 | 9 | 90 | 90 | 0 | 90 | 4.525 | 0 | *a* |
| | | da-GAL4/+ | 10 | 12 | 120 | 120 | 0 | 120 | 4.375 | -3.31 | *b* |
| | | da-GAL4>UAS-cat | 10 | 12 | 120 | 118 | 2 | 120 | 4.975 | +9.94 | *c* |
| $w^{Dah}$ male | 90% $O_2$ | UAS-cat/+ | 10 | 12 | 120 | 119 | 0 | 119 | 5.475 | 0 | *d* |
| | | da-GAL4/+ | 10 | 12 | 120 | 110 | 10 | 120 | 5.400 | -1.37 | *e* |
| | | da-GAL4>UAS-cat | 10 | 12 | 120 | 120 | 0 | 120 | 5.575 | +1.83 | *f* |

p-value (Log-Rank test)

| | *b* | *c* |
|---|---|---|
| *a* | 0.6057 | 1.7E-08 |
| *b* | | 1.4E-10 |

| | *e* | *f* |
|---|---|---|
| *d* | 0.4073 | 8.1E-03 |
| *e* | | 5.2E-04 |

**Fig. S1e / Fig. S1f**

| Fig. | Background | Sex | Genotype | Flies per vial | # vials | Total flies (set-up) | Deaths | Censors | Total flies (actual) | Median (d) | ΔMedian (%) | |
|---|---|---|---|---|---|---|---|---|---|---|---|---|
| Fig. S1e | $w^{Dah+}$ | female | da-GAL4/+ | 15 | 10 | 150 | 145 | 3 | 148 | 57.0 | 0 | *a* |
| | | | UAS-cat/+ | 15 | 10 | 150 | 149 | 2 | 151 | 57.3 | +0.53 | *b* |
| | | | da-GAL4>UAS-cat | 15 | 10 | 150 | 146 | 1 | 147 | 63.0 | +10.53 | *c* |
| | | | UAS-mito-cat/+ | 15 | 10 | 150 | 147 | 1 | 148 | 54.8 | -3.86 | *d* |
| | | | da-GAL4>UAS-mito-cat | 15 | 10 | 150 | 146 | 4 | 150 | 56.7 | -0.53 | *e* |
| Fig. S1f | $w^{Dah+}$ | male | da-GAL4/+ | 20 | 10 | 200 | 187 | 9 | 196 | 61.5 | 0 | *f* |
| | | | UAS-cat/+ | 20 | 10 | 200 | 195 | 6 | 201 | 61.4 | -0.16 | *g* |
| | | | da-GAL4>UAS-cat | 20 | 10 | 200 | 192 | 4 | 196 | 62.0 | +0.81 | *h* |
| | | | UAS-mito-cat/+ | 20 | 10 | 200 | 203 | 4 | 207 | 59.8 | -2.76 | *i* |
| | | | da-GAL4>UAS-mito-cat | 20 | 10 | 200 | 192 | 6 | 198 | 62.1 | +0.98 | *j* |

p-value (Log-Rank test)

| | *b* | *c* | *d* | *e* |
|---|---|---|---|---|
| *a* | 0.8414 | 3.5E-07 | 0.0539 | 0.3451 |
| *b* | | 4.4E-08 | 0.0541 | 0.4352 |
| *c* | | | 9.07E-12 | 1.5E-09 |
| *d* | | | | 0.2589 |

| | *g* | *h* | *i* | *j* |
|---|---|---|---|---|
| *f* | 0.6437 | 0.7669 | 0.1325 | 0.4905 |
| *g* | | 0.4550 | 0.2864 | 0.9281 |
| *h* | | | 0.0616 | 0.3259 |
| *i* | | | | 0.3096 |

**Fig. S1h**

| Background/ Sex | Genotype | Condition | Flies per vial | # vials | Total flies (set-up) | Deaths | Censors | Total flies (actual) | Median (d) | ΔMedian (%) | |
|---|---|---|---|---|---|---|---|---|---|---|---|
| $w^{Dah}$ female | da-GS> UAS-cat | 0 µM RU | 15 | 17 | 255 | 252 | 4 | 256 | 75.3 | 0 | *a* |
| | | 50 µM RU | 15 | 16 | 240 | 241 | 2 | 243 | 82.5 | +9.56 | *b* |
| | | 200 µM RU | 15 | 17 | 255 | 246 | 5 | 251 | 83.8 | +11.29 | *c* |
| | | 400 µM RU | 15 | 17 | 255 | 215 | 40 | 255 | 86.7 | +15.14 | *d* |
| $w^{Dah}$ male | da-GS> UAS-cat | 0 µM RU | 15 | 17 | 255 | 239 | 16 | 255 | 60.9 | 0 | *e* |
| | | 50 µM RU | 15 | 17 | 255 | 247 | 3 | 250 | 63.1 | +3.61 | *f* |
| | | 200 µM RU | 15 | 17 | 255 | 244 | 6 | 250 | 61.3 | +0.66 | *g* |
| | | 400 µM RU | 15 | 17 | 255 | 245 | 8 | 253 | 61.4 | +0.82 | *h* |

p-value (Log-Rank test)

| | *b* | *c* | *d* |
|---|---|---|---|
| *a* | 3.94E-12 | 1.26E-16 | 4.82E-27 |
| *b* | | 0.1145 | 3.90E-06 |
| *c* | | | 0.0016 |

| | *f* | *g* | *h* |
|---|---|---|---|
| *e* | 0.0027 | 0.2355 | 0.4520 |
| *f* | | 0.0620 | 0.0280 |
| *g* | | | 0.7127 |

**Fig. S1i**

| Background | Sex | Genotype | Flies per vial | # vials | Total flies (set-up) | Deaths | Censors | Total flies (actual) | Median (d) | ΔMedian (%) | |
|---|---|---|---|---|---|---|---|---|---|---|---|
| $w^{Dah}$ | female | +/+ | 10 | 20 | 200 | 191 | 8 | 199 | 72.1 | 0 | *a* |
| | | UAS-cat/+ | 10 | 20 | 200 | 184 | 11 | 195 | 69.9 | -3.05 | *b* |
| | | act5c-GAL4/+ | 10 | 20 | 200 | 192 | 5 | 197 | 74.3 | +3.05 | *c* |
| | | act5c-GAL4>UAS-cat | 10 | 20 | 200 | 192 | 7 | 199 | 81.3 | +12.76 | *d* |

p-value (Log-Rank test)

| | *b* | *c* | *d* |
|---|---|---|---|
| *a* | 0.7335 | 0.0663 | 3.0E-11 |
| *b* | | 0.0266 | 3.6E-12 |
| *c* | | | 1.4E-06 |

**Fig. S1j / Fig. S1k**

| Fig. | Background | Sex | Genotype | Flies per vial | # vials | Total flies (set-up) | Deaths | Censors | Total flies (actual) | Median (d) | ΔMedian (%) | |
|---|---|---|---|---|---|---|---|---|---|---|---|---|
| Fig. S1j | $w^{Dah+}$ | female | +/+ | 10 | 20 | 200 | 185 | 10 | 195 | 58.5 | 0 | *a* |
| | | | UAS-cat/+ | 10 | 20 | 200 | 191 | 13 | 204 | 61.7 | +5.47 | *b* |
| | | | da-GAL4/+ | 10 | 20 | 200 | 195 | 8 | 203 | 61.8 | +5.64 | *c* |
| | | | da-GAL4>UAS-cat | 10 | 20 | 200 | 191 | 5 | 196 | 70.4 | +20.34 | *d* |
| Fig. S1k | $w^{Dah+}$ | male | +/+ | 10 | 20 | 200 | 186 | 10 | 196 | 52.4 | 0 | *e* |
| | | | UAS-cat/+ | 10 | 20 | 200 | 184 | 16 | 200 | 50.1 | -4.39 | *f* |
| | | | da-GAL4/+ | 10 | 20 | 200 | 189 | 9 | 198 | 56.2 | +7.25 | *g* |
| | | | da-GAL4>UAS-cat | 10 | 20 | 200 | 188 | 12 | 200 | 52.1 | -0.57 | *h* |

p-value (Log-Rank test)

| | *b* | *c* | *d* |
|---|---|---|---|
| *a* | 0.0060 | 0.0014 | 3.9E-21 |
| *b* | | 0.7162 | 4.7E-12 |
| *c* | | | 1.5E-11 |

| | *f* | *g* | *h* |
|---|---|---|---|
| *e* | 0.1594 | 0.0251 | 0.4451 |
| *f* | | 0.0004 | 0.0477 |
| *g* | | | 0.1916 |

**Fig. S1l**

| Background/ Sex | Genotype | Condition | Flies per vial | # vials | Total flies (set-up) | Deaths | Censors | Total flies (actual) | Median (d) | ΔMedian (%) | |
|---|---|---|---|---|---|---|---|---|---|---|---|
| $w^{Dah}$ female | da-GS> UAS-cat | –RU (0 µM) | 15 | 18 | 270 | 255 | 3 | 258 | 78.0 | 0 | *a* |
| | | +RU (200 µM) from d2 | 15 | 18 | 270 | 260 | 3 | 263 | 85.3 | +9.36 | *b* |
| | | +RU (200 µM) from d28 | 15 | 18 | 270 | 261 | 1 | 262 | 81.7 | +4.74 | *c* |
| | | +RU (200 µM) from d42 | 15 | 18 | 270 | 250 | 2 | 252 | 82.0 | +5.13 | *d* |
| | | +RU (200 µM) from d56 | 15 | 18 | 270 | 261 | 1 | 262 | 80.9 | +3.72 | *e* |

p-value (Log-Rank test)

| | *b* | *c* | *d* | *e* |
|---|---|---|---|---|
| *a* | 2.4E-22 | 7.2E-08 | 1.0E-07 | 1.4E-03 |
| *b* | | 1.7E-05 | 4.8E-05 | 8.0E-12 |
| *c* | | | 0.8959 | 0.0144 |
| *d* | | | | 0.0153 |

p-value (Log-Rank test)

| | Background | Sex | Condition | Flies per vial | # vials | Total flies (set-up) | Deaths | Censors | Total flies (actual) | Median (d) | ΔMedian (%) | | b | c |
|---|---|---|---|---|---|---|---|---|---|---|---|---|---|---|
| Fig. S1n | $w^{Dah}$ | female | UAS-cat/+ | 15 | 19 | 285 | 279 | 5 | 284 | 68.8 | 0 | *a* | 0.0149 | 0.5810 |
| | | | Uo-GAL4/+ | 15 | 20 | 300 | 276 | 13 | 289 | 67.8 | -1.45 | *b* | | 0.0025 |
| | | | Uo-GAL4>UAS-cat | 15 | 20 | 300 | 287 | 7 | 294 | 69.9 | +1.60 | *c* | | |

| | Background | Sex | Condition | Flies per vial | # vials | Total flies (set-up) | Deaths | Censors | Total flies (actual) | Median (d) | ΔMedian (%) | | b | c |
|---|---|---|---|---|---|---|---|---|---|---|---|---|---|---|
| Fig. S1o | $w^{Dah}$ | female | UAS-cat/+ | 15 | 19 | 285 | 279 | 5 | 284 | 68.8 | 0 | *a* | 0.2270 | 0.0023 |
| | | | dilp2-GAL4/+ | 15 | 20 | 300 | 280 | 8 | 288 | 68.0 | -1.16 | *b* | | 3.7E-05 |
| | | | dilp2-GAL4>UAS-cat | 15 | 20 | 300 | 293 | 11 | 304 | 70.7 | +2.76 | *c* | | |

p-value (Log-Rank test)

| | Background/ Sex | Genotype | Condition | Flies per vial | # vials | Total flies (set-up) | Deaths | Censors | Total flies (actual) | Median (d) | ΔMedian (%) | | b |
|---|---|---|---|---|---|---|---|---|---|---|---|---|---|
| Fig. S1p | $w^{Dah}$ female | elav-GS> | –RU (0 μM) | 15 | 10 | 150 | 134 | 17 | 151 | 57.4 | 0 | *a* | 0.0317 |
| | | UAS-cat | +RU (200 μM) | 15 | 10 | 150 | 148 | 5 | 153 | 59 | +2.79 | *b* | |

p-value (Log-Rank test)

| | Background/ Sex | Genotype | Condition | Flies per vial | # vials | Total flies (set-up) | Deaths | Censors | Total flies (actual) | Median (d) | ΔMedian (%) | | b |
|---|---|---|---|---|---|---|---|---|---|---|---|---|---|
| Fig. S1q | $w^{Dah}$ female | $S_1$106-GS> | –RU (0 μM) | 15 | 15 | 225 | 221 | 5 | 226 | 65.8 | 0 | *a* | 0.0941 |
| | | UAS-cat | +RU (200 μM) | 15 | 15 | 225 | 208 | 14 | 222 | 67.6 | +2.74 | *b* | |

p-value (Log-Rank test)

| | Background/ Sex | Genotype | Condition | Flies per vial | # vials | Total flies (set-up) | Deaths | Censors | Total flies (actual) | Median (d) | ΔMedian (%) | | b |
|---|---|---|---|---|---|---|---|---|---|---|---|---|---|
| Fig. S1r | $w^{Dah}$ female | TIGS2> | –RU (0 μM) | 15 | 10 | 150 | 141 | 4 | 145 | 66.8 | 0 | *a* | 0.6981 |
| | | UAS-cat | +RU (200 μM) | 15 | 10 | 150 | 148 | 4 | 152 | 67.4 | +0.90 | *b* | |

p-value (Log-Rank test)

| | Background/ Sex | Genotype | Condition | Flies per vial | # vials | Total flies (set-up) | Deaths | Censors | Total flies (actual) | Median (d) | ΔMedian (%) | | b |
|---|---|---|---|---|---|---|---|---|---|---|---|---|---|
| Fig. S1s | $w^{Dah}$ female | MHC-GS> | –RU (0 μM) | 15 | 10 | 150 | 150 | 2 | 152 | 67.6 | 0 | *a* | 4.96E-05 |
| | | UAS-cat | +RU (200 μM) | 15 | 10 | 150 | 139 | 5 | 144 | 62.1 | -8.14 | *b* | |

p-value (Log-Rank test)

| | Background/ Sex | Condition | Genotype | Flies per vial | # vials | Total flies (set-up) | Deaths | Censors | Total flies (actual) | Median (d) | ΔMedian (%) | | b |
|---|---|---|---|---|---|---|---|---|---|---|---|---|---|
| Fig. 2b | $w^{Dah}$ female | 0.1x yeast | UAS-cat/+ | 10 | 16 | 160 | 157 | 4 | 161 | 18.25 | 0 | *a* | 3.6E-10 |
| | | | da-GAL4>UAS-cat | 10 | 16 | 160 | 158 | 2 | 160 | 21.45 | +17.53 | *b* | |

p-value (Log-Rank test)

| | Background/ Sex | Condition | Genotype | Flies per vial | # vials | Total flies (set-up) | Deaths | Censors | Total flies (actual) | Median (d) | ΔMedian (%) | | b | c |
|---|---|---|---|---|---|---|---|---|---|---|---|---|---|---|
| Fig. 2c | $w^{Dah}$ female | starvation (1.5% agar) | UAS-cat/+ | 10 | 8 | 80 | 80 | 0 | 80 | 7.050 | 0 | *a* | 0.7521 | 5.6E-05 |
| | | | da-GAL4/+ | 10 | 12 | 120 | 120 | 0 | 120 | 7.125 | +1.06 | *b* | | 7.6E-07 |
| | | | da-GAL4>UAS-cat | 10 | 12 | 120 | 120 | 0 | 120 | 6.275 | -10.99 | *c* | | |
| | | | | | | | | | | | | | *e* | *f* |
| | $w^{Dah}$ male | starvation (1.5% agar) | UAS-cat/+ | 10 | 12 | 120 | 120 | 0 | 120 | 3.650 | 0 | *d* | 0.7746 | 0.3292 |
| | | | da-GAL4/+ | 10 | 12 | 120 | 120 | 0 | 120 | 3.700 | +1.37 | *e* | | 0.4287 |
| | | | da-GAL4>UAS-cat | 10 | 12 | 120 | 120 | 0 | 120 | 3.625 | -0.68 | *f* | | |

p-value (Log-Rank test)

| | Background/ Sex | Condition | Genotype | Flies per vial | # vials | Total flies (set-up) | Deaths | Censors | Total flies (actual) | Median (d) | ΔMedian (%) | | b | | |
|---|---|---|---|---|---|---|---|---|---|---|---|---|---|---|---|
| Fig. 2h | $w^{Dah+}$ female | da-GS> | –RU (0 μM) | 15 | 16 | 240 | 231 | 10 | 241 | 81.6 | 0 | *a* | 0.4177 | | |
| | | UAS-Atg5RNAi | +RU (200 μM) | 15 | 5 | 225 | 229 | 2 | 231 | 79.5 | -2.57 | *b* | | | |
| | | | | | | | | | | | | | *d* | *e* | *f* |
| Fig. 2i | $w^{Dah+}$ female | da-GS> | –RU (0 μM) | 15 | 5 | 225 | 224 | 2 | 226 | 79.4 | 0 | *c* | 7.2E-15 | 0.1938 | 0.0026 |
| | | UAS-cat | +RU (200 μM) | 15 | 5 | 225 | 224 | 1 | 225 | 89.0 | +12.09 | *d* | | 3.4E-09 | 2.0E-06 |
| | | da-GS>UAS-cat+ | –RU (0 μM) | 15 | 5 | 225 | 223 | 4 | 227 | 80.8 | +1.76 | *e* | | | 0.1701 |
| | | UAS-Atg5RNAi | +RU (200 μM) | 15 | 5 | 225 | 214 | 12 | 226 | 83.7 | +5.42 | *f* | | | |

p-value (Log-Rank test)

| | Background/ Sex | Condition | Genotype | Flies per vial | # vials | Total flies (set-up) | Deaths | Censors | Total flies (actual) | Median (d) | ΔMedian (%) | | b | c |
|---|---|---|---|---|---|---|---|---|---|---|---|---|---|---|
| Fig. S2f | $w^{Dah+}$ female | 10 mM chloroquine | UAS-cat/+ | 20 | 8 | 160 | 147 | 0 | 147 | 10.4 | 0 | *a* | 0.0988 | 2.6E-05 |
| | | | da-GAL4/+ | 20 | 8 | 160 | 156 | 0 | 156 | 10.8 | +3.85 | *b* | | 3.5E-09 |
| | | | da-GAL4>UAS-cat | 20 | 8 | 160 | 156 | 0 | 156 | 9.45 | -9.13 | *c* | | |

p-value (Log-Rank test)

| | Background/ Sex | Condition | Genotype | Flies per vial | # vials | Total flies (set-up) | Deaths | Censors | Total flies (actual) | Median (d) | ΔMedian (%) | | b | c | d |
|---|---|---|---|---|---|---|---|---|---|---|---|---|---|---|---|
| Fig. S2h | $w^{Dah+}$ female | starvation (1.5% agar) | +/+ | 20 | 6 | 120 | 114 | 0 | 114 | 8.825 | 0 | *a* | 0.4035 | 0.1846 | 3.9E-18 |
| | | | UAS-Atg5RNAi/+ | 20 | 7 | 140 | 130 | 0 | 130 | 8.675 | -1.70 | *b* | | 0.0297 | 1.6E-17 |
| | | | da-GAL4/+ | 20 | 5 | 100 | 103 | 0 | 103 | 8.975 | +1.70 | *c* | | | 7.4E-22 |
| | | | da-GAL4>UAS-Atg5RNAi | 20 | 1 | 20 | 22 | 0 | 22 | 7.000 | -20.68 | *d* | | | |

p-value (Log-Rank test)

| | Background/ Sex | Atg4a | Genotype | Flies per vial | # vials | Total flies (set-up) | Deaths | Censors | Total flies (actual) | Median (d) | ΔMedian (%) | | b | c |
|---|---|---|---|---|---|---|---|---|---|---|---|---|---|---|
| Fig. 4e | $w^{Dah+}$ female | Atg4a-WT | +/+ | 15 | 9 | 135 | 125 | 2 | 127 | 67.1 | 0 | *a* | 0.4857 | 1.8E-08 |
| | | | da-GAL4/+ | 15 | 10 | 150 | 139 | 2 | 141 | 67.9 | +1.19 | *b* | | 6.0E-08 |
| | | | da-GAL4>UAS-cat | 15 | 10 | 150 | 144 | 2 | 146 | 74.5 | +11.03 | *c* | | |
| | | | | | | | | | | | | | *e* | *f* |
| Fig. 4f | $w^{Dah+}$ female | Atg4a-C102S | +/+ | 15 | 10 | 150 | 144 | 1 | 145 | 68.9 | 0 | *d* | 4.9E-04 | 0.2154 |
| | | | da-GAL4/+ | 15 | 10 | 150 | 145 | 3 | 148 | 65.1 | -5.52 | *e* | | 0.0165 |
| | | | da-GAL4>UAS-cat | 15 | 10 | 150 | 141 | 1 | 142 | 67.8 | -1.60 | *f* | | |

p-value (Log-Rank test)

| | Background/ Sex/Condition | Atg4a | Genotype | Flies per vial | # vials | Total flies (set-up) | Deaths | Censors | Total flies (actual) | Median (d) | ΔMedian (%) | | b | c | d |
|---|---|---|---|---|---|---|---|---|---|---|---|---|---|---|---|
| Fig. S4d | $w^{Dah+}$ female, 5% $H_2O_2$ | Atg4a-WT | +/+ | 20 | 10 | 200 | 199 | 0 | 199 | 4.15 | 0 | *a* | 0.0836 | 0.0421 | 9.98E-95 |
| | | | UAS-cat/+ | 20 | 10 | 200 | 200 | 0 | 200 | 4.60 | 10.84 | *b* | | 0.5679 | 4.80E-92 |
| | | | da-GAL4/+ | 20 | 8 | 160 | 149 | 0 | 149 | 4.25 | 2.41 | *c* | | | 6.19E-86 |
| | | | da-GAL4>UAS-cat | 20 | 10 | 200 | 189 | 0 | 189 | 11.05 | 166.27 | *d* | | | |
| | | | | | | | | | | | | | *f* | *g* | *h* |
| | $w^{Dah+}$ female, 5% $H_2O_2$ | Atg4a-C102S | +/+ | 20 | 10 | 200 | 196 | 0 | 196 | 4.40 | 0 | *e* | 0.0094 | 0.3001 | 5.0E-93 |
| | | | UAS-cat/+ | 20 | 10 | 200 | 194 | 0 | 194 | 4.60 | +4.55 | *f* | | 0.1531 | 4.8E-90 |
| | | | da-GAL4/+ | 20 | 8 | 160 | 160 | 0 | 160 | 4.35 | -1.14 | *g* | | | 3.6E-87 |
| | | | da-GAL4>UAS-cat | 20 | 10 | 200 | 199 | 0 | 199 | 10.75 | +144.32 | *h* | | | |

**Supplementary Table 2 | List of cysteine residues identified in this study by OxICAT redox proteomics**
Lennicke *et al.*

| Uniprot ID | Cys # | Name | UAS-cat/+ | | | | | | | | | da-GAL4>UAS-cat | | | | | | | | |
|---|---|---|---|---|---|---|---|---|---|---|---|---|---|---|---|---|---|---|---|---|
| | | | d7 | | | d28 | | | d56 | | | d7 | | | d28 | | | d56 | | |
| | | | % ox. | SD | n | % ox. | SD | n | % ox. | SD | n | % ox. | SD | n | % ox. | SD | n | % ox. | SD | n |
| **A1Z7H3** | 218 | Acyl-CoA synthetase long-chain (Lethal (2) 44DEa; isoform I) | | | | | | | | | | 31.8 | 7.5 | 4 | 48.6 | 8.9 | 5 | | | |
| **A1Z8U9** | 109 | Ribosomal protein S11; isoform B | | | | | | | | | | | | | 19.3 | 4.1 | 3 | | | |
| **A1Z8U9** | 127 | Ribosomal protein S11; isoform B | 11.1 | 2.7 | 4 | 14.4 | 3.1 | 3 | | | | 13.6 | 1.5 | 4 | 19.0 | 1.4 | 5 | 17.6 | 8.1 | 3 |
| **A1Z992** | 187 | 1,4-Alpha-glucan branching enzyme (CG33138) | 8.4 | 2.3 | 5 | 5.8 | 1.3 | 5 | 9.1 | 1.8 | 4 | 9.4 | 1.0 | 4 | 16.1 | 2.7 | 3 | 17.8 | 5.0 | 4 |
| **A1Z992** | 294 | 1,4-Alpha-glucan branching enzyme (CG33138) | | | | | | | | | | 11.7 | 0.1 | 3 | 18.3 | 3.5 | 4 | | | |
| **A1Z992** | 652 | 1,4-Alpha-glucan branching enzyme (CG33138) | | | | | | | | | | 10.7 | 1.0 | 4 | | | | | | |
| **A1Z9E3** | 83 | Elongation factor Tu | 14.5 | 2.3 | 5 | 12.5 | 3.8 | 4 | 12.4 | 3.2 | 5 | 14.0 | 1.6 | 5 | 19.5 | 4.2 | 5 | 17.5 | 4.9 | 5 |
| **A1Z9E3** | 152 | Elongation factor Tu | | | | | | | | | | 13.6 | 1.0 | 4 | 18.3 | 4.8 | 5 | 17.4 | 5.4 | 5 |
| **A1Z9E3** | 246 | Elongation factor Tu | | | | | | | | | | | | | 17.2 | 2.5 | 3 | 20.0 | 2.1 | 5 |
| **A1Z9G2** | 122 | fandango (FI18620p1) | 12.6 | 2.3 | 5 | 11.8 | 3.0 | 5 | 8.6 | 2.8 | 3 | 13.6 | 1.5 | 5 | 19.9 | 2.2 | 5 | 22.6 | 9.4 | 5 |
| **A1ZA47** | 131 | PDZ and LIM domain protein Zasp | 12.5 | 2.4 | 5 | | | | | | | | | | | | | | | |
| **A1ZA47** | 305 | PDZ and LIM domain protein Zasp | | | | 23.0 | 8.0 | 4 | 30.5 | 8.4 | 5 | 30.7 | 2.8 | 5 | 32.3 | 5.7 | 5 | 30.7 | 9.2 | 4 |
| **A1ZA47** | 329 | PDZ and LIM domain protein Zasp | 13.3 | 3.9 | 5 | 10.9 | 3.2 | 5 | 11.6 | 3.9 | 5 | 13.0 | 1.6 | 5 | 18.5 | 2.4 | 5 | 18.2 | 5.8 | 5 |
| **A1ZA47** | 497 | PDZ and LIM domain protein Zasp | 12.6 | 1.9 | 5 | 12.7 | 3.6 | 5 | 10.8 | 3.0 | 5 | 12.4 | 1.5 | 5 | 20.4 | 1.2 | 4 | 19.3 | 0.5 | 3 |
| **A1ZA47** | 741 | PDZ and LIM domain protein Zasp | | | | | | | | | | 6.6 | 0.3 | 3 | 9.8 | 1.9 | 3 | | | |
| **A1ZA47** | 823 | PDZ and LIM domain protein Zasp | 9.6 | 3.2 | 4 | 9.1 | 0.3 | 3 | 8.8 | 2.6 | 4 | 8.3 | 0.6 | 5 | 13.4 | 3.7 | 5 | 16.6 | 5.6 | 5 |
| **A1ZA47** | 1210 | PDZ and LIM domain protein Zasp | | | | | | | | | | | | | 14.2 | 4.3 | 5 | | | |
| **A1ZA47** | 1539 | PDZ and LIM domain protein Zasp | | | | | | | 10.2 | 3.5 | 3 | 8.8 | 0.4 | 4 | 13.5 | 3.2 | 5 | 17.8 | 5.1 | 5 |
| **A1ZA47** | 1776 | PDZ and LIM domain protein Zasp | | | | 6.6 | 1.4 | 3 | | | | 7.6 | 0.4 | 4 | 14.6 | 3.9 | 5 | 10.6 | 3.8 | 5 |
| **A1ZA47** | 2081 | PDZ and LIM domain protein Zasp | 21.5 | 1.5 | 5 | 18.5 | 4.8 | 5 | 20.1 | 5.0 | 5 | 26.4 | 3.0 | 5 | 26.7 | 2.8 | 5 | 24.1 | 4.8 | 5 |
| **A1ZA47** | 2092 | PDZ and LIM domain protein Zasp | 21.4 | 3.1 | 5 | 22.2 | 7.7 | 5 | 22.0 | 8.8 | 5 | 28.2 | 2.3 | 5 | 33.6 | 4.1 | 5 | 27.0 | 7.1 | 5 |
| **A1ZA47** | 2189 | PDZ and LIM domain protein Zasp | 17.0 | 2.4 | 5 | 15.8 | 4.3 | 4 | 14.5 | 4.0 | 5 | | | | | | | 20.3 | 0.7 | 3 |
| **A1ZA73** | 103 | Stretchin-Mlck; isoform E | 10.1 | 3.0 | 5 | 7.8 | 2.6 | 5 | 8.7 | 2.4 | 5 | 7.8 | 0.6 | 5 | 13.4 | 4.7 | 5 | 15.9 | 5.9 | 5 |
| **A1ZA73** | 477 | Stretchin-Mlck; isoform E | 9.9 | 0.4 | 3 | 8.3 | 1.6 | 3 | 14.3 | 4.6 | 4 | 7.8 | 0.9 | 5 | 14.2 | 5.8 | 5 | 12.4 | 5.7 | 5 |
| **A1ZA73** | 591 | Stretchin-Mlck; isoform E | 4.2 | 1.1 | 5 | 5.9 | 2.1 | 5 | 6.6 | 3.9 | 5 | 5.7 | 0.7 | 5 | 10.5 | 3.8 | 5 | 7.1 | 2.8 | 5 |
| **A1ZA73** | 610 | Stretchin-Mlck; isoform E | 9.5 | 2.9 | 4 | 6.9 | 2.7 | 3 | 11.5 | 2.8 | 4 | 6.6 | 1.3 | 5 | 10.0 | 4.9 | 5 | 10.1 | 4.8 | 5 |
| **A1ZA73** | 799 | Stretchin-Mlck; isoform E | | | | 17.1 | 2.2 | 3 | 21.7 | 2.9 | 4 | 16.3 | 1.6 | 4 | 18.3 | 5.5 | 4 | 20.8 | 6.8 | 4 |
| **A1ZA73** | 1124 | Stretchin-Mlck; isoform E | | | | | | | | | | 4.0 | 0.7 | 3 | 8.7 | 5.9 | 3 | | | |
| **A1ZA73** | 1267 | Stretchin-Mlck; isoform E | 7.4 | 2.6 | 5 | 5.8 | 1.7 | 5 | 7.8 | 3.0 | 5 | 6.7 | 0.8 | 5 | 10.5 | 4.8 | 5 | 11.3 | 5.1 | 5 |
| **A1ZA73** | 1300 | Stretchin-Mlck; isoform E | | | | | | | | | | 14.0 | 1.1 | 3 | 19.1 | 5.1 | 3 | | | |
| **A1ZA73** | 1330 | Stretchin-Mlck; isoform E | 20.2 | 1.7 | 3 | 18.5 | 2.5 | 3 | 22.2 | 3.8 | 4 | 19.6 | 0.5 | 4 | 24.4 | 9.9 | 3 | | | |
| **A1ZA73** | 1401 | Stretchin-Mlck; isoform E | 9.5 | 1.5 | 5 | 8.8 | 2.5 | 5 | 10.0 | 3.2 | 5 | 10.4 | 0.5 | 5 | 15.9 | 6.0 | 5 | 14.9 | 4.5 | 5 |
| **A1ZA73** | 1418 | Stretchin-Mlck; isoform E | 13.7 | 2.6 | 5 | 9.7 | 2.6 | 5 | 11.8 | 2.9 | 5 | 12.1 | 0.9 | 5 | 17.3 | 7.2 | 5 | 18.8 | 6.4 | 5 |
| **A1ZA73** | 1593 | Stretchin-Mlck; isoform E | | | | | | | | | | 11.8 | 3.7 | 3 | | | | | | |
| **A1ZA73** | 2166 | Stretchin-Mlck; isoform E | 12.5 | 3.8 | 5 | | | | 10.5 | 2.5 | 4 | | | | | | | | | |
| **A1ZA73** | 2277 | Stretchin-Mlck; isoform E | 15.2 | 4.5 | 4 | | | | | | | | | | | | | 22.8 | 6.8 | 3 |
| **A1ZBJ2** | 328 | Acyl-CoA dehydrogenase very long chain (CG7461) | | | | | | | 41.3 | 9.5 | 3 | 27.5 | 2.6 | 4 | 13.6 | 4.7 | 5 | 18.1 | 8.9 | 3 |
| **A4UZR3** | 21 | Trehalase; isoform E | | | | | | | | | | 11.2 | 0.7 | 4 | 16.3 | 3.6 | 4 | | | |
| **A4UZR3** | 261 | Trehalase; isoform E | 20.0 | 3.7 | 3 | | | | | | | 11.7 | 1.6 | 4 | | | | 20.2 | 3.7 | 3 |
| **A4V4Q6** | 78 | wings up A; isoform F | 7.5 | 2.1 | 5 | 6.9 | 1.5 | 4 | 6.8 | 4.6 | 4 | 10.6 | 3.0 | 3 | | | | | | |
| **A4V4Q6** | 96 | wings up A; isoform F | 6.4 | 1.7 | 5 | | | | 8.6 | 2.3 | 4 | 6.3 | 0.8 | 5 | 10.6 | 1.9 | 5 | 14.1 | 7.6 | 5 |
| **A4V4Q6** | 106 | wings up A; isoform F | 10.7 | 3.6 | 4 | 12.6 | 0.9 | 3 | 11.7 | 0.8 | 3 | 9.5 | 1.2 | 4 | 16.5 | 4.5 | 5 | 20.2 | 5.7 | 5 |
| **A5XCL5** | 125 | UGP; isoform D | | | | | | | | | | | | | 16.0 | 1.9 | 4 | | | |
| **A5XCL5** | 194 | UGP; isoform D | | | | | | | | | | 8.0 | 0.3 | 4 | 12.2 | 2.4 | 4 | 12.2 | 4.1 | 5 |
| **A5XCL5** | 380 | UGP; isoform D | | | | | | | | | | 5.7 | 0.5 | 4 | 9.4 | 2.3 | 5 | 11.2 | 4.1 | 3 |
| **A8JRC2** (obsolete) | 22 | Isocitrate dehydrogenase (NAD(+)) 3 non-catalytic subunit gamma (CG5028; isoform E) | 5.1 | 1.6 | 3 | | | | 6.5 | 3.0 | 4 | 5.2 | 0.2 | 5 | 8.6 | 2.1 | 5 | 7.6 | 2.9 | 5 |
| **A8JRC2** (obsolete) | 90 | Isocitrate dehydrogenase (NAD(+)) 3 non-catalytic subunit gamma (CG5028; isoform E) | 10.5 | 1.7 | 5 | 9.7 | 2.5 | 5 | 10.4 | 2.6 | 5 | 10.8 | 1.5 | 5 | 14.0 | 2.4 | 5 | 15.3 | 4.6 | 5 |
| **A8Y535** | 47 | Ubiquinol-cytochrome C reductase complex 11 kDa subunit; isoform D | 82.0 | 7.0 | 5 | 90.5 | 3.4 | 5 | 88.0 | 5.7 | 5 | 83.1 | 9.7 | 5 | 82.6 | 6.0 | 5 | 85.7 | 8.3 | 5 |
| **B7YZP9** | 35 | Muscle LIM protein at 60A; isoform B | 44.8 | 1.3 | 5 | 42.0 | 9.4 | 5 | 41.5 | 6.5 | 5 | 47.0 | 5.6 | 3 | | | | | | |
| **B7YZP9** | 50 | Muscle LIM protein at 60A; isoform B | 27.4 | 2.5 | 5 | | | | | | | | | | 38.8 | 7.9 | 5 | 34.0 | 10.7 | 5 |
| **B7YZP9** | 80 | Muscle LIM protein at 60A; isoform B | 18.5 | 2.3 | 5 | 17.6 | 4.6 | 5 | 16.8 | 5.2 | 5 | 19.9 | 3.2 | 5 | 28.2 | 4.5 | 5 | 24.6 | 7.8 | 5 |
| **B7YZP9** | 108 | Muscle LIM protein at 60A; isoform B | 38.1 | 2.4 | 4 | 35.0 | 11.2 | 4 | 28.9 | 1.8 | 3 | | | | | | | 39.2 | 6.3 | 5 |
| **B7YZP9** | 213 | Muscle LIM protein at 60A; isoform B | 30.6 | 2.7 | 5 | 26.7 | 8.2 | 5 | 30.5 | 6.2 | 3 | 36.1 | 3.9 | 5 | | | | | | |
| **B7YZP9** | 216 | Muscle LIM protein at 60A; isoform B | 34.5 | 5.8 | 4 | 24.0 | 2.9 | 3 | 31.4 | 7.8 | 3 | 39.9 | 4.2 | 5 | 36.6 | 4.5 | 5 | 33.5 | 9.1 | 4 |
| **B7YZP9** | 461 | Muscle LIM protein at 60A; isoform B | 28.1 | 3.2 | 3 | 23.7 | 6.2 | 3 | 28.4 | 4.7 | 5 | 32.8 | 1.7 | 5 | 30.4 | 5.8 | 5 | 37.1 | 7.0 | 3 |
| **B7YZQ7** | 25 | Nucleosome remodeling factor - 38kD; isoform B | | | | | | | | | | | | | 20.0 | 3.1 | 4 | | | |
| **B7Z061** | 162 | Photoreceptor dehydrogenase; isoform D | 6.8 | 2.2 | 5 | 7.3 | 1.6 | 5 | 9.1 | 4.9 | 5 | | | | 13.3 | 3.8 | 3 | | | |
| **B7Z061** | 223 | Photoreceptor dehydrogenase; isoform D | | | | 9.8 | 2.2 | 3 | | | | 8.5 | 2.6 | 4 | 14.3 | 3.7 | 5 | 13.7 | 3.8 | 4 |
| **E1JH64** | 1922 | short stop; isoform Q | | | | | | | | | | | | | 18.1 | 11.3 | 3 | | | |
| **E1JH64** | 2575 | short stop; isoform Q | | | | | | | | | | 9.8 | 1.0 | 5 | 14.8 | 6.6 | 5 | 13.2 | 4.8 | 5 |
| **E1JH64** | 2978 | short stop; isoform Q | | | | | | | | | | 4.1 | 0.9 | 4 | 6.2 | 2.2 | 4 | | | |
| **E1JHJ5** | 38 | Myosin heavy chain; isoform P | 8.5 | 0.6 | 5 | 7.3 | 2.2 | 5 | 7.3 | 2.2 | 5 | 9.2 | 1.2 | 5 | 13.8 | 3.3 | 5 | 13.0 | 4.1 | 5 |
| **E1JHJ5** | 389 | Myosin heavy chain; isoform P | 7.3 | 2.2 | 5 | 9.7 | 1.6 | 4 | 14.4 | 7.4 | 5 | 9.0 | 1.1 | 5 | 16.7 | 3.3 | 5 | 13.0 | 3.9 | 5 |
| **E1JHJ5** | 427 | Myosin heavy chain; isoform P | | | | 4.0 | 0.9 | 4 | 4.5 | 3.2 | 5 | 3.1 | 0.4 | 5 | 8.1 | 1.7 | 5 | 4.9 | 2.6 | 4 |
| **E1JHJ5** | 442 | Myosin heavy chain; isoform P | 2.4 | 1.0 | 5 | 3.0 | 0.8 | 5 | 4.6 | 3.1 | 5 | 2.3 | 0.2 | 5 | 4.7 | 1.4 | 5 | 3.9 | 2.3 | 5 |
| **E1JHJ5** | 673 | Myosin heavy chain; isoform P | 8.5 | 2.3 | 5 | 10.3 | 2.8 | 5 | 8.4 | 5.2 | 5 | 11.9 | 1.8 | 5 | 19.0 | 3.0 | 5 | 12.0 | 3.0 | 5 |
| **E1JHJ5** | 740 | Myosin heavy chain; isoform P | 9.4 | 2.4 | 5 | 7.8 | 2.8 | 3 | 9.2 | 2.7 | 4 | 10.2 | 1.3 | 5 | 14.1 | 4.1 | 5 | 13.0 | 4.1 | 5 |
| **E1JHJ5** | 1223 | Myosin heavy chain; isoform P | 16.4 | 1.4 | 5 | 14.9 | 4.9 | 5 | 19.3 | 6.6 | 5 | 17.4 | 1.0 | 5 | 20.8 | 4.8 | 5 | 24.7 | 6.7 | 5 |
| **E1JHJ5** | 1234 | Myosin heavy chain; isoform P | 16.8 | 1.2 | 5 | 15.0 | 4.5 | 5 | 18.0 | 6.0 | 5 | 18.6 | 1.3 | 5 | 21.6 | 4.1 | 5 | 22.6 | 5.6 | 5 |
| **E1JHJ5** | 1410 | Myosin heavy chain; isoform P | 9.0 | 3.3 | 5 | 5.8 | 2.4 | 5 | 7.7 | 4.4 | 5 | 8.8 | 1.2 | 5 | 13.7 | 5.5 | 5 | 15.2 | 7.7 | 5 |
| **E1JHJ5** | 1473 | Myosin heavy chain; isoform P | 8.6 | 2.3 | 5 | 6.5 | 2.0 | 5 | 9.5 | 3.9 | 5 | 6.8 | 0.7 | 5 | 11.7 | 2.8 | 5 | 13.6 | 5.6 | 5 |
| **E1JHR5** | 400 | Enolase; isoform F | 14.4 | 2.2 | 5 | 16.7 | 4.3 | 5 | 12.8 | 4.1 | 5 | | | | | | | 21.2 | 0.1 | 3 |
| **E1JIR4** | 240 | Na pump alpha subunit; isoform I | 4.7 | 1.6 | 5 | 3.9 | 1.3 | 5 | 4.8 | 2.6 | 5 | 4.5 | 0.4 | 5 | 7.0 | 3.4 | 5 | 7.4 | 3.6 | 5 |
| **E1JIR4** | 353 | Na pump alpha subunit; isoform I | 8.4 | 1.8 | 5 | 6.6 | 2.6 | 5 | 7.1 | 2.7 | 5 | 8.1 | 0.2 | 5 | 14.5 | 5.3 | 5 | 12.3 | 4.1 | 5 |
| **E1JIR4** | 407 | Na pump alpha subunit; isoform I | 2.6 | 0.9 | 5 | 3.3 | 1.0 | 5 | 4.3 | 3.3 | 4 | 3.1 | 0.4 | 5 | 8.3 | 3.0 | 5 | 4.2 | 1.4 | 5 |
| **E1JIR4** | 497 | Na pump alpha subunit; isoform I | | | | 4.0 | 2.1 | 5 | | | | 3.2 | 0.4 | 5 | 9.3 | 5.6 | 5 | 6.7 | 3.3 | 5 |
| **E1JIR4** | 535 | Na pump alpha subunit; isoform I | | | | | | | | | | 7.8 | 0.3 | 4 | 14.3 | 5.4 | 5 | | | |
| **E1JIR4** | 684 | Na pump alpha subunit; isoform I | 6.6 | 1.8 | 5 | 5.5 | 1.3 | 4 | 6.2 | 3.3 | 5 | 6.0 | 0.4 | 5 | 8.5 | 1.9 | 4 | 10.8 | 4.7 | 5 |
| **E2QCF1** | 29 | ATP citrate lyase; isoform F | 18.0 | 2.6 | 5 | 13.0 | 3.2 | 5 | 13.9 | 3.6 | 5 | 14.2 | 1.6 | 5 | 22.6 | 4.9 | 5 | 21.8 | 6.0 | 4 |
| **O01666** | 105 | ATP synthase subunit gamma; mitochondrial | 9.4 | 2.7 | 5 | 9.6 | 2.2 | 5 | 8.8 | 3.7 | 5 | 10.2 | 0.6 | 5 | 17.2 | 3.2 | 5 | 14.2 | 4.7 | 5 |
| **O01666** | 131 | ATP synthase subunit gamma; mitochondrial | 8.1 | 2.0 | 5 | 7.1 | 1.7 | 5 | 7.4 | 3.6 | 5 | 8.0 | 0.7 | 5 | 15.6 | 2.3 | 5 | 13.3 | 4.3 | 5 |
| **O01666** | 197 | ATP synthase subunit gamma; mitochondrial | | | | | | | | | | | | | 12.4 | 4.0 | 5 | 14.0 | 4.9 | 4 |
| **O18404** | 85 | 3-hydroxyacyl-CoA dehydrogenase type-2 | | | | | | | | | | | | | 12.9 | 1.9 | 3 | | | |
| **O46037** | 623 | Vinculin | | | | | | | | | | 7.2 | 1.6 | 3 | | | | | | |
| **O46231** | 67 | Tropomodulin; isoform A | | | | | | | | | | | | | 12.4 | 1.1 | 3 | 12.3 | 3.3 | 4 |
| **O61231** | 105 | 60S ribosomal protein L10 | | | | | | | | | | | | | 23.1 | 6.0 | 3 | | | |
| **O62619** | 34 | Pyruvate kinase | 12.6 | 1.6 | 5 | 9.6 | 1.6 | 5 | 10.2 | 3.7 | 5 | 11.6 | 0.5 | 5 | 15.7 | 3.3 | 5 | 16.0 | 5.2 | 5 |
| **O62619** | 198 | Pyruvate kinase | 6.3 | 1.8 | 5 | 5.4 | 1.6 | 5 | 6.8 | 3.8 | 5 | 6.7 | 0.4 | 5 | 12.7 | 3.6 | 5 | 12.6 | 4.7 | 5 |
| **O62619** | 329 | Pyruvate kinase | | | | | | | 15.5 | 7.2 | 3 | 17.5 | 4.6 | 5 | 18.7 | 3.3 | 4 | 15.2 | 3.9 | 3 |
| **O62619** | 361 | Pyruvate kinase | | | | | | | | | | 12.4 | 0.5 | 4 | 18.2 | 4.8 | 4 | | | |
| **O62619** | 377 | Pyruvate kinase | | | | | | | | | | 18.1 | 1.6 | 4 | 25.3 | 4.7 | 5 | 20.5 | 7.1 | 3 |
| **O62619** | 385 | Pyruvate kinase | 13.8 | 3.3 | 4 | 12.3 | 2.0 | 3 | 11.8 | 5.3 | 4 | 49.0 | 31.6 | 3 | 41.6 | 29.4 | 3 | | | |
| **O62619** | 450 | Pyruvate kinase | 27.2 | 4.9 | 5 | 24.2 | 6.9 | 5 | 22.0 | 6.1 | 5 | 22.3 | 1.9 | 5 | 38.4 | 5.4 | 5 | 28.9 | 7.3 | 5 |
| **O76927** | 56 | 40S ribosomal protein S21 | | | | | | | | | | 26.9 | 2.8 | 3 | 41.3 | 11.7 | 5 | 35.0 | 8.7 | 5 |
| **O97394** | 1447 | Protein sidekick | | | | 13.5 | 4.6 | 3 | 17.9 | 5.2 | 5 | 15.8 | 1.6 | 5 | 21.7 | 3.9 | 5 | 16.2 | 5.5 | 3 |
| **O97418** | 58 | NADH dehydrogenase (ubiquinone) B14.5 A subunit (SD16673p) | 14.3 | 2.3 | 5 | 14.1 | 2.7 | 5 | 14.8 | 3.2 | 5 | 14.6 | 2.4 | 5 | 21.6 | 4.9 | 5 | 24.2 | 6.0 | 5 |
| **P00334** | 136 | Alcohol dehydrogenase | 7.8 | 2.1 | 5 | 7.6 | 1.0 | 4 | 11.6 | 4.7 | 4 | 9.2 | 0.9 | 5 | 12.4 | 3.4 | 5 | 16.2 | 5.6 | 4 |
| **P00334** | 219 | Alcohol dehydrogenase | 9.9 | 2.9 | 5 | 9.5 | 1.8 | 5 | 9.6 | 4.8 | 5 | 10.8 | 0.8 | 5 | 18.4 | 2.8 | 5 | 18.2 | 5.7 | 5 |
| **P02574** | 53 | Actin 79B (Actin; larval muscle) | 9.8 | 1.8 | 5 | 7.5 | 2.4 | 5 | 9.0 | 4.4 | 5 | 9.7 | 0.7 | 5 | 14.9 | 2.9 | 5 | 15.9 | 6.0 | 5 |
| **P02574** | 218 | Actin 79B (Actin; larval muscle) | 7.3 | 1.8 | 5 | 6.0 | 1.6 | 5 | 8.0 | 3.6 | 5 | 7.5 | 1.0 | 5 | 12.4 | 2.8 | 5 | 13.3 | 5.3 | 5 |
| **P02574** | 273 | Actin 79B (Actin; larval muscle) | 12.8 | 1.7 | 5 | 10.4 | 2.2 | 5 | 11.7 | 3.3 | 5 | 11.0 | 0.4 | 5 | 17.3 | 3.3 | 5 | 18.5 | 6.0 | 5 |
| **P02574** | 286 | Actin 79B (Actin; larval muscle) | 11.8 | 2.3 | 5 | 9.7 | 2.9 | 5 | 11.5 | 3.7 | 5 | 11.9 | 1.1 | 5 | 17.3 | 3.6 | 5 | 19.4 | 7.0 | 5 |
| **P02828** | 460 | Heat shock protein 83 | | | | | | | | | | 7.6 | 0.2 | 4 | 12.7 | 2.5 | 5 | 11.9 | 3.1 | 4 |
| **P04359** | 95 | Ribosomal protein L32 (60S ribosomal protein L32) | | | | | | | | | | | | | 13.5 | 2.1 | 5 | | | |
| **P06605** | 295 | alpha-Tubulin at 84D (Tubulin alpha-3 chain) | 10.1 | 2.0 | 5 | 9.0 | 2.2 | 5 | 9.6 | 2.0 | 5 | 10.9 | 1.0 | 5 | 15.2 | 1.9 | 5 | 14.5 | 4.1 | 5 |
| **P06605** | 347 | alpha-Tubulin at 84D (Tubulin alpha-3 chain) | 8.7 | 3.2 | 4 | 8.3 | 2.6 | 4 | 10.3 | 3.6 | 4 | 8.5 | 0.4 | 4 | 14.8 | 3.0 | 5 | 14.0 | 3.8 | 4 |
| **P06605** | 376 | alpha-Tubulin at 84D (Tubulin alpha-3 chain) | | | | | | | | | | | | | 14.5 | 1.0 | 3 | | | |
| **P06607** | 103 | Yolk protein 3 (Vitellogenin-3) | 15.3 | 4.3 | 5 | 11.3 | 2.2 | 5 | | | | | | | | | | 24.0 | 9.4 | 3 |
| **P06754** | 25 | Tropomyosin-1, isoforms 9A/A/B | 21.9 | 13.5 | 5 | 9.7 | 6.5 | 5 | 11.8 | 6.9 | 5 | 10.0 | 0.8 | 5 | 15.6 | 3.7 | 5 | 20.5 | 7.0 | 5 |
| **P06754** | 118 | Tropomyosin-1, isoforms 9A/A/B | | | | | | | | | | 11.9 | 9.1 | 3 | 15.9 | 4.7 | 4 | | | |
| **P07486** | 130 | Glyceraldehyde-3-phosphate dehydrogenase 1 | 13.8 | 2.5 | 5 | 11.1 | 2.4 | 5 | 12.5 | 4.5 | 5 | 16.1 | 1.6 | 5 | 17.6 | 3.2 | 5 | 19.3 | 6.4 | 5 |
| **P07487** | 130 | Glyceraldehyde-3-phosphate dehydrogenase 2 | 13.2 | 2.0 | 5 | 11.7 | 2.4 | 5 | 12.5 | 4.0 | 5 | 14.9 | 1.2 | 5 | 17.6 | 3.0 | 5 | 18.5 | 4.8 | 5 |

| | | | | | | | | | | | | | | | | | | | | |
|---|---|---|---|---|---|---|---|---|---|---|---|---|---|---|---|---|---|---|---|---|
| P07764 | 143 | Aldolase 1 (Fructose-bisphosphate aldolase) | 15.5 | 1.8 | 5 | 14.2 | 2.5 | 5 | 15.9 | 6.3 | 5 | 18.6 | 1.1 | 5 | 21.7 | 2.8 | 5 | 22.3 | 6.0 | 5 |
| P07764 | 150 | Aldolase 1 (Fructose-bisphosphate aldolase) | 16.6 | 2.8 | 5 | 12.5 | 3.9 | 5 | 15.4 | 8.3 | 5 | 17.8 | 0.3 | 3 | 20.8 | 1.8 | 3 | 23.3 | 3.6 | 4 |
| P07764 | 178 | Aldolase 1 (Fructose-bisphosphate aldolase) | 7.8 | 2.0 | 5 | 7.9 | 1.9 | 5 | 8.0 | 4.7 | 5 | 9.0 | 1.2 | 5 | 14.3 | 2.2 | 5 | 13.2 | 4.1 | 5 |
| P08736 | 411 | Elongation factor 1-alpha 1 | 10.3 | 1.5 | 5 | | | | | | | 10.1 | 0.8 | 5 | 15.7 | 2.0 | 5 | 13.0 | 2.2 | 5 |
| P08879 | 110 | abnormal wing discs (Nucleoside diphosphate kinase) | | | | 8.8 | 1.6 | 3 | 9.1 | 5.1 | 5 | 8.5 | 2.0 | 3 | 15.4 | 4.3 | 4 | 19.7 | 4.2 | 4 |
| P08928 | 150 | Lamin Dm0 | 6.5 | 1.7 | 5 | 5.6 | 2.0 | 5 | 6.5 | 3.7 | 5 | 7.1 | 2.0 | 5 | 12.1 | 2.4 | 5 | 12.2 | 5.0 | 4 |
| P09180 | 26 | 60S ribosomal protein L4 | | | | | | | | | | 7.4 | 0.3 | 3 | | | | | | |
| P09180 | 99 | 60S ribosomal protein L4 | 15.2 | 4.1 | 3 | | | | | | | 15.1 | 3.7 | 5 | 26.4 | 4.1 | 5 | | | |
| P09491 | 25 | Tropomyosin-2 | 9.5 | 1.9 | 5 | 8.3 | 1.9 | 5 | 10.0 | 3.2 | 5 | 10.4 | 1.0 | 5 | 18.4 | 3.7 | 5 | 21.8 | 6.8 | 5 |
| P09491 | 127 | Tropomyosin-2 | 11.2 | 2.3 | 5 | 9.6 | 2.2 | 5 | 11.4 | 3.7 | 5 | | | | | | | 26.4 | 6.2 | 3 |
| P09491 | 271 | Tropomyosin-2 | 6.0 | 1.8 | 5 | 6.3 | 0.4 | 3 | 8.2 | 0.7 | 3 | | | | 14.6 | 2.1 | 4 | 16.3 | 4.4 | 5 |
| P11046 | 756 | Laminin subunit beta-1 | | | | | | | | | | 85.5 | 5.0 | 4 | | | | | | |
| P11147 | 267 | Heat shock 70 kDa protein cognate 4 | | | | | | | | | | | | | | | | 8.7 | 1.7 | 3 |
| P11147 | 546 | Heat shock 70 kDa protein cognate 4 | | | | | | | | | | | | | 13.7 | 3.3 | 3 | | | |
| P11147 | 603 | Heat shock 70 kDa protein cognate 4 | | | | | | | | | | | | | 14.5 | 3.8 | 3 | | | |
| P12024 | 64 | Chaoptin | 81.8 | 3.2 | 5 | 77.3 | 7.0 | 4 | | | | 76.8 | 0.9 | 5 | | | | | | |
| P12024 | 1260 | Chaoptin | | | | | | | 73.1 | 11.9 | 4 | 71.4 | 11.9 | 5 | 89.1 | 1.5 | 5 | | | |
| P12982 | 125 | Serine/threonine-protein phosphatase alpha-2 isoform | | | | | | | | | | 16.2 | 1.6 | 3 | 19.0 | 5.3 | 5 | 17.2 | 5.8 | 4 |
| P13060 | 553 | Elongation factor 2 | | | | | | | | | | | | | 17.9 | 6.9 | 5 | | | |
| P13060 | 714 | Elongation factor 2 | 10.8 | 1.2 | 5 | 8.9 | 2.4 | 5 | 9.8 | 4.5 | 5 | 11.2 | 0.3 | 5 | 15.6 | 2.5 | 5 | 14.9 | 4.4 | 5 |
| P13217 | 148 | no receptor potential A (1-phosphatidylinositol 4,5-bisphosphate phosphodiesterase) | | | | | | | | | | 17.0 | 2.5 | 4 | | | | | | |
| P13395 | 325 | Spectrin alpha chain | | | | | | | | | | 12.9 | 3.4 | 3 | 15.5 | 1.9 | 4 | 17.9 | 11.3 | 3 |
| P13395 | 681 | Spectrin alpha chain | | | | | | | | | | | | | 12.9 | 6.1 | 3 | 15.2 | 1.4 | 3 |
| P13395 | 1570 | Spectrin alpha chain | | | | 5.5 | 1.9 | 4 | 6.8 | 3.0 | 5 | 7.0 | 0.3 | 5 | 11.1 | 2.6 | 5 | 11.2 | 3.3 | 5 |
| P13395 | 2068 | Spectrin alpha chain | | | | | | | | | | 5.9 | 0.2 | 3 | | | | | | |
| P15372 | 154 | Arrestin 1 (Phosrestin-2) | 9.6 | 2.1 | 5 | 7.8 | 2.1 | 3 | | | | 13.8 | 2.1 | 3 | 17.1 | 2.8 | 5 | 18.2 | 0.7 | 3 |
| P17336 | 93 | Catalase | | | | | | | | | | 8.8 | 0.7 | 5 | 15.9 | 3.4 | 5 | 14.7 | 4.9 | 5 |
| P17336 | 208 | Catalase | | | | | | | | | | 18.3 | 0.8 | 5 | 23.6 | 4.1 | 5 | 22.5 | 7.0 | 5 |
| P17336 | 375 | Catalase | | | | | | | | | | 29.1 | 1.1 | 5 | 44.5 | 9.2 | 5 | 40.2 | 7.4 | 5 |
| P17336 | 413 | Catalase | | | | | | | | | | 43.2 | 8.1 | 4 | 40.9 | 4.9 | 4 | | | |
| P17336 | 455 | Catalase | | | | | | | | | | 22.3 | 2.7 | 5 | 27.1 | 5.6 | 5 | 23.7 | 7.9 | 4 |
| P17704 | 35 | 40S ribosomal protein S17 | | | | | | | | | | | | | 13.5 | 4.3 | 5 | 18.4 | 4.4 | 4 |
| P18930 | 41 | Mitochondrial NADH-ubiquinone oxidoreductase chain 3 | 10.1 | 3.2 | 4 | 11.6 | 3.5 | 5 | | | | 10.0 | 1.1 | 5 | 20.8 | 1.5 | 5 | 14.5 | 4.1 | 4 |
| P19967 | 409 | Cytochrome b5-related protein | | | | | | | | | | | | | 8.8 | 2.1 | 4 | | | |
| P20228 | 89 | Glutamate decarboxylase | | | | | | | | | | | | | 7.4 | 1.7 | 3 | | | |
| P20432 | 13 | Glutathione S-transferase D1 | | | | | | | 12.9 | 3.1 | 5 | 17.1 | 1.6 | 4 | | | | 22.1 | 5.7 | 5 |
| P20478 | 48 | Glutamine synthetase 2 cytoplasmic | 12.7 | 1.6 | 5 | | | | | | | | | | | | | | | |
| P20478 | 105 | Glutamine synthetase 2 cytoplasmic | 15.0 | 3.4 | 5 | 12.5 | 4.2 | 5 | 14.0 | 5.6 | 5 | 17.0 | 0.7 | 5 | 20.7 | 3.1 | 5 | 22.1 | 6.6 | 5 |
| P20478 | 123 | Glutamine synthetase 2 cytoplasmic | 20.4 | 2.2 | 5 | 17.4 | 3.9 | 5 | 18.9 | 5.0 | 5 | 21.9 | 0.9 | 5 | 24.5 | 3.4 | 5 | 25.9 | 6.1 | 5 |
| P20478 | 169 | Glutamine synthetase 2 cytoplasmic | 19.7 | 2.5 | 3 | 17.5 | 3.1 | 4 | | | | | | | | | | | | |
| P20478 | 189 | Glutamine synthetase 2 cytoplasmic | | | | | | | | | | 11.2 | 0.5 | 4 | 12.0 | 0.7 | 3 | | | |
| P20478 | 327 | Glutamine synthetase 2 cytoplasmic | 13.5 | 2.5 | 5 | 13.4 | 3.3 | 5 | 13.1 | 5.9 | 5 | 14.5 | 0.6 | 5 | 21.0 | 4.5 | 4 | 18.7 | 6.5 | 4 |
| P20478 | 352 | Glutamine synthetase 2 cytoplasmic | | | | | | | | | | 13.9 | 0.3 | 4 | 17.3 | 3.9 | 4 | 15.6 | 6.4 | 4 |
| P21914 | 75 | Succinate dehydrogenase [ubiquinone]iron-sulfur subunit; mitochondrial | | | | | | | | | | 5.4 | 0.6 | 4 | 9.1 | 1.0 | 3 | | | |
| P22464 | 11 | Annexin B9 | | | | | | | 11.9 | 3.0 | 3 | | | | 19.3 | 2.2 | 5 | 18.5 | 1.0 | 4 |
| P22700 | 471 | Calcium-transporting ATPase sarcoplasmic/endoplasmic reticulum type | 7.5 | 1.8 | 5 | 5.7 | 1.8 | 5 | 7.3 | 2.6 | 5 | 6.7 | 0.8 | 5 | 12.0 | 6.7 | 5 | 10.7 | 4.9 | 5 |
| P22700 | 498 | Calcium-transporting ATPase sarcoplasmic/endoplasmic reticulum type | 7.6 | 1.7 | 5 | 6.9 | 2.1 | 5 | 6.9 | 3.3 | 5 | 8.4 | 0.4 | 5 | 14.0 | 6.5 | 5 | 9.9 | 3.2 | 5 |
| P22700 | 525 | Calcium-transporting ATPase sarcoplasmic/endoplasmic reticulum type | 7.1 | 1.5 | 5 | 6.3 | 2.1 | 5 | 5.4 | 3.4 | 5 | | | | | | | 9.3 | 1.0 | 3 |
| P22700 | 561 | Calcium-transporting ATPase sarcoplasmic/endoplasmic reticulum type | | | | | | | | | | 7.4 | 0.8 | 4 | 11.6 | 6.8 | 3 | | | |
| P22700 | 636 | Calcium-transporting ATPase sarcoplasmic/endoplasmic reticulum type | 4.7 | 1.0 | 5 | 4.7 | 1.1 | 5 | 6.1 | 2.3 | 5 | 4.7 | 0.3 | 5 | 8.7 | 4.9 | 5 | 6.3 | 3.3 | 5 |
| P22700 | 876 | Calcium-transporting ATPase sarcoplasmic/endoplasmic reticulum type | | | | | | | | | | 44.2 | 9.2 | 4 | 69.0 | 4.4 | 5 | | | |
| P25007 | 103 | Cyclophilin 1 (Peptidyl-prolyl cis-trans isomerase) | | | | | | | 9.6 | 8.3 | 4 | 8.8 | 0.1 | 5 | 17.6 | 2.6 | 5 | 15.1 | 4.8 | 5 |
| P25007 | 125 | Cyclophilin 1 (Peptidyl-prolyl cis-trans isomerase) | 12.3 | 3.8 | 4 | 14.8 | 3.1 | 5 | 12.0 | 5.9 | 5 | 15.4 | 2.8 | 5 | 23.1 | 2.5 | 5 | 19.9 | 4.5 | 5 |
| P25007 | 178 | Cyclophilin 1 (Peptidyl-prolyl cis-trans isomerase) | | | | | | | | | | 11.0 | 2.9 | 4 | 10.4 | 2.1 | 3 | | | |
| P29310 | 97 | 14-3-3 protein zeta | | | | | | | 3.9 | 0.3 | 3 | | | | 4.6 | 1.4 | 4 | 5.1 | 2.5 | 4 |
| P29413 | 105 | Calreticulin | | | | | | | | | | 66.6 | 3.7 | 4 | 73.4 | 6.2 | 3 | | | |
| P29413 | 312 | Calreticulin | | | | | | | | | | 71.1 | 3.3 | 3 | | | | | | |
| P29613 | 6 | Triose phosphate isomerase | 18.3 | 3.2 | 5 | 20.7 | 2.6 | 5 | 21.4 | 11.5 | 5 | 37.0 | 15.0 | 5 | 29.9 | 6.6 | 5 | | | |
| P29613 | 42 | Triose phosphate isomerase | 24.2 | 4.7 | 5 | 27.0 | 5.0 | 5 | 22.8 | 8.1 | 5 | 24.4 | 2.0 | 5 | 39.0 | 4.0 | 5 | 30.4 | 6.8 | 5 |
| P29613 | 56 | Triose phosphate isomerase | 16.0 | 4.3 | 5 | 20.8 | 4.2 | 5 | 16.0 | 8.0 | 5 | 19.4 | 2.1 | 5 | 28.5 | 3.8 | 5 | 23.9 | 5.5 | 5 |
| P29613 | 125 | Triose phosphate isomerase | 13.1 | 1.6 | 5 | 10.0 | 2.4 | 5 | 11.7 | 5.8 | 4 | 12.1 | 2.0 | 5 | 18.6 | 4.7 | 5 | 18.1 | 6.1 | 5 |
| P29613 | 147 | Triose phosphate isomerase | 20.0 | 2.9 | 5 | 16.3 | 4.1 | 5 | 17.3 | 7.1 | 5 | 18.5 | 1.9 | 5 | 25.4 | 5.8 | 5 | 25.8 | 7.0 | 5 |
| P29742 | 871 | Clathrin heavy chain | | | | | | | | | | 9.7 | 2.0 | 5 | 16.1 | 6.0 | 5 | | | |
| P29829 | 32 | Guanine nucleotide-binding protein subunit beta-2 | | | | | | | | | | 24.3 | 0.5 | 4 | 25.4 | 4.1 | 5 | 23.9 | 6.2 | 4 |
| P29829 | 74 | Guanine nucleotide-binding protein subunit beta-2 | | | | | | | | | | | | | 11.6 | 2.1 | 4 | | | |
| P29829 | 155 | Guanine nucleotide-binding protein subunit beta-2 | | | | | | | | | | | | | 20.5 | 4.2 | 4 | 19.0 | 7.0 | 3 |
| P29829 | 173 | Guanine nucleotide-binding protein subunit beta-2 | | | | | | | | | | | | | 20.3 | 2.2 | 3 | | | |
| P29829 | 282 | Guanine nucleotide-binding protein subunit beta-2 | | | | | | | | | | 21.2 | 3.0 | 5 | 23.7 | 4.4 | 5 | 26.2 | 7.7 | 4 |
| P31009 | 165 | 40S ribosomal protein S2 | | | | 16.2 | 4.6 | 5 | | | | 22.2 | 2.2 | 5 | 28.1 | 5.8 | 5 | | | |
| P31009 | 205 | 40S ribosomal protein S2 | | | | 28.8 | 0.8 | 3 | | | | | | | | | | | | |
| P31409 | 91 | V-type proton ATPase subunit B | | | | 10.4 | 3.2 | 3 | 10.6 | 3.7 | 4 | 9.2 | 1.5 | 5 | 15.9 | 3.7 | 5 | 16.6 | 6.5 | 5 |
| P31409 | 186 | V-type proton ATPase subunit B | | | | | | | 8.8 | 4.2 | 3 | 5.0 | 0.3 | 4 | 9.4 | 2.2 | 5 | 9.1 | 4.6 | 5 |
| P31409 | 405 | V-type proton ATPase subunit B | | | | | | | | | | 12.0 | 2.7 | 4 | 15.3 | 2.0 | 4 | 21.1 | 5.5 | 4 |
| P35381 | 243 | ATP synthase subunit alpha; mitochondrial | 8.8 | 2.4 | 4 | 5.3 | 1.1 | 4 | 8.2 | 2.9 | 5 | 7.0 | 0.6 | 5 | 9.2 | 2.7 | 5 | 11.1 | 5.3 | 5 |
| P35381 | 489 | ATP synthase subunit alpha; mitochondrial | 8.4 | 4.0 | 3 | | | | 10.6 | 4.4 | 3 | | | | 10.7 | 2.1 | 5 | 13.5 | 5.8 | 5 |
| P35415 | 368 | Paramyosin; long form | 10.9 | 2.2 | 3 | 7.9 | 3.2 | 3 | 8.9 | 3.1 | 5 | | | | | | | 16.1 | 7.2 | 3 |
| P35415 | 784 | Paramyosin; long form | | | | | | | 11.9 | 1.8 | 4 | | | | 9.9 | 1.3 | 4 | | | |
| P38979 | 206 | 40S ribosomal protein SA | | | | | | | | | | | | | 17.2 | 2.6 | 4 | | | |
| P39018 | 93 | 40S ribosomal protein S19a | 13.0 | 3.8 | 5 | 14.9 | 3.9 | 5 | 13.0 | 5.4 | 5 | 17.1 | 4.9 | 4 | 19.4 | 4.1 | 4 | 16.3 | 5.3 | 4 |
| P46461 | 10 | Vesicle-fusing ATPase 1 | 13.4 | 3.1 | 5 | | | | | | | | | | | | | 21.4 | 6.4 | 3 |
| P48148 | 107 | Ras-like GTP-binding protein Rho1 | 15.0 | 3.9 | 5 | 15.1 | 3.0 | 5 | 14.5 | 4.6 | 5 | 15.7 | 1.7 | 5 | 21.6 | 3.0 | 5 | 17.9 | 3.6 | 5 |
| P48375 | 77 | FK506-binding protein 12 kDa | 9.2 | 1.6 | 5 | 8.3 | 2.7 | 5 | 9.6 | 4.9 | 5 | 9.4 | 1.4 | 5 | 15.8 | 3.7 | 5 | 16.7 | 4.5 | 5 |
| P48554 | 178 | Ras-related protein Rac2 | | | | | | | | | | | | | 37.4 | 6.2 | 4 | | | |
| P48602 | 391 | V-type proton ATPase catalytic subunit A isoform 1 | 8.2 | 2.1 | 3 | 7.3 | 0.3 | 3 | 7.5 | 4.1 | 3 | | | | 12.6 | 2.4 | 4 | 13.5 | 2.4 | 3 |
| P48610 | 139 | Arginine kinase | 18.5 | 2.1 | 5 | 16.4 | 4.9 | 5 | 18.6 | 4.3 | 5 | 20.0 | 2.8 | 5 | 25.6 | 4.8 | 5 | 23.4 | 5.1 | 5 |
| P48610 | 201 | Arginine kinase | 10.5 | 2.1 | 5 | 11.8 | 2.2 | 5 | 11.3 | 6.0 | 5 | 12.9 | 1.2 | 5 | 20.3 | 3.2 | 5 | 17.8 | 4.2 | 5 |
| P48610 | 222 | Arginine kinase | 6.5 | 2.2 | 5 | 5.9 | 1.2 | 5 | 8.0 | 5.0 | 5 | 6.8 | 1.0 | 5 | 10.8 | 2.6 | 5 | 13.5 | 5.4 | 5 |
| P48610 | 271 | Arginine kinase | 9.2 | 2.4 | 5 | 9.7 | 2.3 | 5 | 9.4 | 4.5 | 5 | 10.9 | 1.1 | 5 | 18.1 | 3.2 | 5 | 16.7 | 5.1 | 5 |
| P49455 | 268 | Tropomyosin-1, isoforms 33/34 | 7.6 | 1.3 | 5 | 6.6 | 1.5 | 5 | 7.7 | 2.1 | 5 | 8.6 | 0.9 | 5 | 12.8 | 2.8 | 5 | 15.9 | 4.7 | 5 |
| P50882 | 9 | 60S ribosomal protein L9 | 9.2 | 2.2 | 4 | 11.1 | 4.2 | 4 | | | | 10.7 | 0.2 | 4 | 15.7 | 1.7 | 5 | 14.2 | 1.8 | 4 |
| P52029 | 408 | Glucose-6-phosphate isomerase | 7.9 | 1.6 | 5 | 8.2 | 2.2 | 5 | 7.2 | 4.1 | 5 | 9.5 | 1.2 | 5 | 15.2 | 3.1 | 5 | 11.2 | 3.0 | 5 |
| P52029 | 459 | Glucose-6-phosphate isomerase | | | | 8.8 | 3.3 | 3 | | | | 10.0 | 0.7 | 5 | 17.5 | 4.0 | 5 | 12.8 | 3.7 | 5 |
| P52029 | 536 | Glucose-6-phosphate isomerase | | | | | | | | | | 7.3 | 1.1 | 5 | 13.1 | 2.1 | 5 | 9.2 | 2.9 | 4 |
| P52034 | 91 | 6-phosphofructokinase | 6.6 | 1.3 | 5 | 5.6 | 2.2 | 4 | 6.0 | 3.2 | 5 | | | | 11.2 | 1.2 | 3 | 10.6 | 4.5 | 4 |
| P52034 | 246 | 6-phosphofructokinase | | | | | | | | | | 10.1 | 1.8 | 3 | | | | | | |
| P52034 | 316 | 6-phosphofructokinase | 7.8 | 1.8 | 4 | 7.3 | 1.6 | 3 | 8.2 | 2.5 | 4 | | | | | | | | | |
| P52034 | 353 | 6-phosphofructokinase | | | | | | | | | | 11.1 | 3.2 | 5 | 19.3 | 3.7 | 4 | 14.9 | 5.1 | 4 |
| P52034 | 422 | 6-phosphofructokinase | 12.2 | 2.1 | 5 | 12.1 | 3.5 | 5 | 11.0 | 4.6 | 5 | 12.1 | 0.7 | 5 | 20.3 | 4.2 | 5 | 15.5 | 3.9 | 5 |
| P52034 | 663 | 6-phosphofructokinase | 9.1 | 2.3 | 5 | 6.8 | 2.7 | 4 | 8.8 | 3.0 | 5 | | | | | | | 13.8 | 6.6 | 4 |
| P52034 | 692 | 6-phosphofructokinase | | | | | | | | | | 3.4 | 0.3 | 3 | 9.0 | 1.6 | 5 | | | |
| P52034 | 712 | 6-phosphofructokinase | 7.1 | 1.4 | 5 | 6.4 | 1.8 | 5 | 7.1 | 2.6 | 4 | 7.7 | 0.4 | 5 | 14.1 | 4.6 | 5 | 9.8 | 3.0 | 5 |
| P53777 | 80 | Muscle LIM protein 1 | 25.2 | 1.8 | 5 | 24.8 | 3.8 | 5 | 22.2 | 4.2 | 5 | 25.8 | 2.9 | 5 | 35.4 | 4.0 | 5 | 32.7 | 6.4 | 5 |
| P54385 | 69 | Glutamate dehydrogenase, mitochondrial | | | | | | | | | | 8.8 | 0.5 | 3 | | | | | | |
| P54385 | 245 | Glutamate dehydrogenase, mitochondrial | 9.2 | 1.8 | 5 | 7.7 | 1.8 | 5 | 8.7 | 2.6 | 5 | 9.5 | 1.4 | 5 | 15.2 | 2.6 | 5 | 14.4 | 4.5 | 5 |
| P54385 | 368 | Glutamate dehydrogenase, mitochondrial | | | | | | | | | | 6.2 | 0.6 | 4 | 10.2 | 2.9 | 3 | 11.4 | 4.7 | 4 |
| P54611 | 136 | V-type proton ATPase subunit E | 7.9 | 2.0 | 5 | 7.9 | 2.3 | 5 | 8.0 | 3.9 | 5 | 8.6 | 2.0 | 5 | 15.4 | 2.4 | 5 | 13.3 | 4.1 | 4 |
| P55828 | 38 | 40S ribosomal protein S20 | | | | | | | | | | | | | 9.2 | 1.9 | 4 | | | |
| P55830 | 157 | 40S ribosomal protein S3a | 12.1 | 3.7 | 5 | 10.1 | 2.6 | 4 | 10.5 | 2.9 | 5 | 11.3 | 1.1 | 5 | 15.2 | 4.4 | 4 | 17.7 | 4.0 | 5 |
| P61851 | 7 | Superoxide dismutase 1 [Cu-Zn] | 32.3 | 3.0 | 5 | 25.5 | 4.0 | 5 | 31.2 | 8.4 | 5 | 34.9 | 3.8 | 5 | 39.7 | 8.4 | 5 | 45.7 | 10.8 | 5 |
| P61851 | 35 | Superoxide dismutase 1 [Cu-Zn] | 45.3 | 1.8 | 5 | 41.0 | 6.4 | 5 | 39.3 | 7.4 | 5 | 43.5 | 3.7 | 5 | 49.8 | 5.4 | 5 | 49.2 | 8.3 | 5 |
| P61851 | 56 | Superoxide dismutase 1 [Cu-Zn] | 62.8 | 3.6 | 4 | 57.2 | 4.6 | 5 | 60.0 | 9.5 | 4 | 59.0 | 2.2 | 5 | 68.1 | 5.1 | 5 | 71.8 | 6.3 | 5 |
| P61851 | 92 | Superoxide dismutase 1 [Cu-Zn] | 28.0 | 4.8 | 4 | 24.8 | 8.2 | 5 | 25.3 | 6.4 | 5 | 27.6 | 2.8 | 5 | 37.7 | 4.0 | 5 | 37.4 | 9.2 | 5 |
| P61851 | 145 | Superoxide dismutase 1 [Cu-Zn] | 54.4 | 6.2 | 5 | 48.3 | 7.5 | 5 | 50.9 | 10.3 | 5 | 52.1 | 3.5 | 5 | 63.4 | 8.2 | 5 | 67.6 | 8.8 | 5 |
| P80455 | 46 | 40S ribosomal protein S12 | 9.6 | 2.8 | 4 | 9.8 | 1.7 | 5 | 12.3 | 9.2 | 4 | 9.1 | 0.4 | 5 | 18.0 | 4.8 | 5 | 15.8 | 5.8 | 5 |
| P80455 | 57 | 40S ribosomal protein S12 | | | | | | | | | | 15.4 | 3.2 | 3 | | | | | | |
| P80455 | 76 | 40S ribosomal protein S12 | | | | | | | | | | 11.1 | 2.5 | 3 | 17.1 | 5.0 | 5 | | | |
| P83967 | 286 | Actin; indirect flight muscle | 15.4 | 3.3 | 5 | 11.2 | 2.3 | 5 | 14.0 | 6.3 | 5 | 13.4 | 1.3 | 5 | 20.2 | 6.7 | 4 | 19.7 | 7.8 | 4 |

| | | | | | | | | | | | | | | | | | | | | |
|---|---|---|---|---|---|---|---|---|---|---|---|---|---|---|---|---|---|---|---|---|
| **P91929** | 86 | NADH dehydrogenase (ubiquinone) 42 kDa subunit | 7.0 | 2.1 | 5 | 6.3 | 1.4 | 5 | 7.7 | 4.0 | 5 | 8.3 | 1.1 | 5 | 12.8 | 1.9 | 5 | 14.9 | 6.1 | 5 |
| **P91929** | 139 | NADH dehydrogenase (ubiquinone) 42 kDa subunit | 12.6 | 1.7 | 5 | 11.6 | 2.3 | 5 | 11.2 | 3.7 | 5 | 14.3 | 0.9 | 5 | 18.4 | 2.2 | 5 | 17.3 | 4.3 | 5 |
| **P91929** | 148 | NADH dehydrogenase (ubiquinone) 42 kDa subunit | | | | | | | | | | 16.6 | 2.2 | 4 | 17.7 | 1.1 | 4 | 16.9 | 4.2 | 5 |
| **P91929** | 333 | NADH dehydrogenase (ubiquinone) 42 kDa subunit | | | | | | | | | | 13.9 | 2.1 | 3 | 16.1 | 0.8 | 4 | | | |
| **P91929** | 340 | NADH dehydrogenase (ubiquinone) 42 kDa subunit | | | | 15.4 | 4.0 | 3 | | | | 17.1 | 1.8 | 4 | 18.5 | 1.4 | 4 | 23.5 | 7.2 | 4 |
| **P91938** | 142 | Thioredoxin reductase 1, mitochondrial | 14.4 | 3.5 | 3 | | | | | | | 17.0 | 2.9 | 3 | 19.7 | 3.2 | 5 | | | |
| **P92177** | 97 | 14-3-3 protein epsilon | | | | | | | | | | 4.0 | 0.6 | 4 | 7.2 | 1.9 | 5 | 7.2 | 3.5 | 4 |
| **P92177** | 111 | 14-3-3 protein epsilon | | | | 7.7 | 2.5 | 4 | 10.3 | 3.3 | 4 | 10.4 | 0.5 | 5 | 17.6 | 3.2 | 5 | 14.1 | 4.5 | 5 |
| **Q00637** | 36 | Superoxide dismutase 2 [Mn]; mitochondrial | 9.8 | 2.0 | 5 | 8.8 | 1.4 | 5 | 9.2 | 3.0 | 4 | 10.6 | 0.9 | 5 | 16.0 | 4.1 | 5 | 14.7 | 4.7 | 5 |
| **Q02645** | 97 | Protein hu-li tai shao | | | | | | | | | | 12.7 | 3.8 | 4 | | | | | | |
| **Q02748** | 149 | Eukaryotic translation initiation factor 4A | | | | | | | | | | 10.2 | 2.1 | 4 | 17.2 | 3.8 | 5 | | | |
| **Q07327** | 375 | Ras opposite (Protein ROP) | | | | | | | | | | | | | 16.2 | 4.9 | 4 | | | |
| **Q0E8V7** | 39 | uncharacterized protein CG34132 (IP17740p) | | | | 84.3 | 0.7 | 3 | | | | | | | | | | | | |
| **Q0KIA8** | 256 | Carnitine O-Acetyl-Transferase (CG1041; isoform B) | | | | | | | | | | 14.2 | 1.2 | 4 | 16.0 | 2.0 | 4 | 16.3 | 6.0 | 3 |
| **Q23997** | 435 | Imaginal disc growth factor 6 (Chitinase-like protein CG5210) | | | | | | | | | | 64.5 | 11.5 | 4 | 85.1 | 6.9 | 4 | | | |
| **Q24048** | 153 | Nervana 2 (Sodium/potassium-transporting ATPase subunit beta-2) | | | | | | | | | | 64.5 | 8.6 | 4 | 80.3 | 5.4 | 4 | | | |
| **Q24048** | 165 | Nervana 2 (Sodium/potassium-transporting ATPase subunit beta-2) | 70.9 | 5.8 | 4 | | | | 74.3 | 9.2 | 3 | 65.4 | 5.0 | 5 | 82.2 | 6.3 | 3 | | | |
| **Q24048** | 175 | Nervana 2 (Sodium/potassium-transporting ATPase subunit beta-2) | 74.1 | 6.0 | 5 | 73.1 | 6.2 | 4 | 68.4 | 10.4 | 4 | 66.9 | 2.8 | 5 | 81.4 | 9.2 | 5 | | | |
| **Q24048** | 189 | Nervana 2 (Sodium/potassium-transporting ATPase subunit beta-2) | | | | | | | | | | 57.2 | 0.8 | 4 | | | | | | |
| **Q24186** | 90 | 40S ribosomal protein S5a | | | | | | | | | | 23.1 | 3.6 | 5 | 35.6 | 9.3 | 5 | | | |
| **Q24186** | 196 | 40S ribosomal protein S5a | | | | | | | | | | | | | 7.7 | 3.5 | 5 | | | |
| **Q24251** | 48 | ATP synthase subunit d, mitochondrial | 7.4 | 3.1 | 5 | 8.4 | 2.4 | 5 | 6.9 | 3.2 | 5 | 8.0 | 0.7 | 5 | 15.3 | 3.3 | 5 | 13.6 | 3.6 | 5 |
| **Q24253** | 381 | AP complex subunit beta (LP17054p) | | | | | | | | | | 5.3 | 0.9 | 4 | 7.7 | 3.7 | 4 | | | |
| **Q24400** | 12 | Muscle LIM protein Mlp84B | 36.0 | 1.1 | 5 | 33.6 | 6.2 | 5 | 28.0 | 5.1 | 5 | | | | | | | 35.2 | 5.2 | 3 |
| **Q24400** | 15 | Muscle LIM protein Mlp84B | | | | 31.8 | 5.0 | 5 | | | | | | | | | | | | |
| **Q24400** | 36 | Muscle LIM protein Mlp84B | 39.5 | 3.0 | 5 | | | | | | | | | | | | | | | |
| **Q24400** | 51 | Muscle LIM protein Mlp84B | 24.2 | 2.7 | 5 | 20.0 | 6.0 | 5 | 22.2 | 6.0 | 5 | 27.5 | 2.5 | 5 | 31.8 | 5.3 | 5 | 28.5 | 8.9 | 5 |
| **Q24400** | 120 | Muscle LIM protein Mlp84B | 35.9 | 2.7 | 5 | 30.7 | 7.6 | 5 | 28.7 | 7.0 | 5 | 39.1 | 3.6 | 5 | 37.4 | 3.4 | 5 | 33.9 | 7.5 | 5 |
| **Q24400** | 123 | Muscle LIM protein Mlp84B | 42.2 | 1.6 | 3 | | | | | | | 46.8 | 6.6 | 5 | 42.0 | 4.4 | 5 | 46.2 | 5.9 | 3 |
| **Q24400** | 195 | Muscle LIM protein Mlp84B | 13.9 | 1.8 | 5 | 13.8 | 4.8 | 5 | 12.5 | 4.7 | 5 | 16.2 | 2.0 | 5 | 21.9 | 2.9 | 5 | 19.6 | 6.1 | 5 |
| **Q24400** | 222 | Muscle LIM protein Mlp84B | 33.7 | 1.3 | 5 | 29.2 | 7.0 | 5 | 27.2 | 5.7 | 5 | 38.2 | 3.5 | 5 | 35.4 | 2.9 | 5 | 33.4 | 6.5 | 5 |
| **Q24400** | 225 | Muscle LIM protein Mlp84B | 35.3 | 2.7 | 5 | 30.3 | 8.1 | 5 | 31.2 | 6.5 | 4 | 38.5 | 2.2 | 5 | 34.7 | 4.9 | 5 | 34.2 | 9.5 | 5 |
| **Q24400** | 271 | Muscle LIM protein Mlp84B | 40.1 | 1.5 | 5 | 33.6 | 8.4 | 5 | 35.7 | 6.6 | 5 | 46.9 | 3.3 | 5 | 41.3 | 6.6 | 5 | 41.5 | 8.8 | 5 |
| **Q24400** | 328 | Muscle LIM protein Mlp84B | 33.8 | 2.4 | 5 | 30.0 | 6.4 | 4 | 31.3 | 4.8 | 4 | 36.1 | 5.8 | 5 | 37.7 | 8.5 | 5 | 37.3 | 9.6 | 5 |
| **Q24400** | 377 | Muscle LIM protein Mlp84B | 30.5 | 2.2 | 5 | 27.9 | 7.6 | 4 | 26.4 | 5.6 | 5 | | | | | | | | | |
| **Q24400** | 470 | Muscle LIM protein Mlp84B | 36.7 | 1.8 | 4 | 32.4 | 5.8 | 4 | 40.2 | 6.4 | 3 | | | | | | | | | |
| **Q24498** | 34 | Ryanodine receptor 44F | | | | | | | | | | 21.9 | 0.8 | 4 | | | | | | |
| **Q24498** | 1633 | Ryanodine receptor 44F | | | | | | | | | | 9.4 | 1.0 | 4 | | | | | | |
| **Q24560** | 12 | Tubulin beta-1 chain | 12.1 | 2.0 | 5 | 9.8 | 2.3 | 5 | 10.2 | 3.0 | 5 | 12.0 | 1.5 | 5 | 16.4 | 3.3 | 5 | 17.3 | 5.0 | 5 |
| **Q24560** | 239 | Tubulin beta-1 chain | | | | | | | | | | 9.4 | 2.2 | 4 | 11.6 | 2.7 | 4 | | | |
| **Q24560** | 303 | Tubulin beta-1 chain | 6.1 | 1.8 | 3 | 5.0 | 1.0 | 5 | 6.6 | 4.6 | 5 | 6.4 | 0.5 | 5 | 10.9 | 2.4 | 5 | 9.7 | 3.7 | 5 |
| **Q24560** | 354 | Tubulin beta-1 chain | 6.6 | 1.8 | 5 | 8.0 | 1.6 | 5 | 6.6 | 2.6 | 5 | 10.1 | 3.7 | 5 | 13.9 | 1.7 | 5 | 11.5 | 2.8 | 5 |
| **Q26365** | 72 | stress-sensitive B (ADP,ATP carrier protein) | 3.8 | 1.3 | 5 | 4.4 | 1.3 | 5 | 4.9 | 3.2 | 5 | 4.5 | 0.2 | 5 | 8.8 | 2.3 | 5 | 6.4 | 2.8 | 5 |
| **Q26365** | 144 | stress-sensitive B (ADP,ATP carrier protein) | 5.0 | 1.5 | 5 | 5.3 | 1.2 | 5 | 6.3 | 3.4 | 5 | 4.9 | 0.4 | 5 | 7.8 | 1.9 | 5 | 7.0 | 3.1 | 5 |
| **Q26365** | 174 | stress-sensitive B (ADP,ATP carrier protein) | 6.1 | 2.8 | 5 | 8.5 | 2.9 | 5 | 6.6 | 5.3 | 5 | 7.0 | 0.8 | 5 | 15.8 | 3.5 | 5 | 9.9 | 3.0 | 5 |
| **Q26365** | 271 | stress-sensitive B (ADP,ATP carrier protein) | 6.6 | 1.4 | 4 | 6.8 | 1.4 | 4 | 7.2 | 3.1 | 5 | 4.7 | 0.4 | 5 | 7.8 | 1.8 | 5 | 4.7 | 1.9 | 3 |
| **Q27268** | 236 | ATP-dependent RNA helicase WM6 | | | | | | | | | | | | | 14.5 | 2.8 | 3 | | | |
| **Q59DP8** | 43 | Plasma membrane calcium ATPase; isoform N | 5.3 | 2.3 | 3 | 4.8 | 2.0 | 4 | 4.6 | 3.8 | 4 | 3.7 | 0.5 | 4 | 11.8 | 6.6 | 5 | 7.5 | 2.3 | 4 |
| **Q59DP8** | 626 | Plasma membrane calcium ATPase; isoform N | | | | | | | | | | 18.0 | 9.6 | 3 | | | | | | |
| **Q59E65 (A8DYP0)** | 1139 | Obscurin (Unc-89; isoform B) | | | | | | | | | | 19.2 | 1.0 | 4 | | | | 24.7 | 7.3 | 3 |
| **Q59E65 (A8DYP0)** | 1537 | Obscurin (Unc-89; isoform B) | | | | | | | | | | 5.0 | 0.9 | 3 | | | | | | |
| **Q59E65 (A8DYP0)** | 2247 | Obscurin (Unc-89; isoform B) | 12.1 | 2.0 | 5 | 8.9 | 2.9 | 5 | 9.8 | 4.0 | 5 | 11.8 | 0.6 | 5 | 16.5 | 8.3 | 5 | 17.2 | 5.7 | 5 |
| **Q59E65 (A8DYP0)** | 2331 | Obscurin (Unc-89; isoform B) | 11.1 | 3.7 | 5 | 7.1 | 2.5 | 5 | 8.7 | 3.7 | 5 | | | | 13.4 | 1.8 | 3 | 16.3 | 7.0 | 4 |
| **Q59E65 (A8DYP0)** | 2509 | Obscurin (Unc-89; isoform B) | | | | | | | | | | 11.5 | 0.1 | 3 | 15.7 | 1.9 | 3 | | | |
| **Q59E65 (A8DYP0)** | 2617 | Obscurin (Unc-89; isoform B) | | | | | | | | | | 26.5 | 2.3 | 4 | | | | | | |
| **Q59E65 (A8DYP0)** | 2730 | Obscurin (Unc-89; isoform B) | 21.7 | 4.3 | 3 | | | | | | | | | | | | | | | |
| **Q59E65 (A8DYP0)** | 3427 | Obscurin (Unc-89; isoform B) | 9.8 | 1.6 | 5 | 9.0 | 2.6 | 5 | 9.1 | 3.8 | 4 | | | | | | | | | |
| **Q59E65 (A8DYP0)** | 3721 | Obscurin (Unc-89; isoform B) | | | | | | | | | | 11.2 | 0.2 | 4 | | | | | | |
| **Q59E65 (A8DYP0)** | 3820 | Obscurin (Unc-89; isoform B) | | | | | | | | | | 5.6 | 0.7 | 3 | 10.9 | 7.6 | 4 | | | |
| **Q59E65 (A8DYP0)** | 4085 | Obscurin (Unc-89; isoform B) | | | | | | | | | | 4.6 | 0.5 | 4 | 10.3 | 4.4 | 5 | 7.7 | 3.2 | 4 |
| **Q59E65 (A8DYP0)** | 4115 | Obscurin (Unc-89; isoform B) | | | | 8.3 | 3.3 | 3 | | | | | | | 13.5 | 3.7 | 4 | | | |
| **Q7JS69** | 34 | Nervana 3; isoform D | | | | | | | | | | 14.5 | 0.4 | 3 | 21.4 | 3.8 | 4 | | | |
| **Q7JUS9** | 101 | Mitochondrial phosphate carrier protein 1 (RH64567p) | 7.0 | 2.2 | 5 | 8.7 | 2.9 | 5 | 6.7 | 4.2 | 5 | | | | 19.9 | 7.5 | 4 | 15.1 | 12.3 | 5 |
| **Q7JVH6** | 185 | uncharacterized protein CG9436 (LD24696p) | | | | 16.3 | 0.9 | 3 | | | | 19.8 | 2.3 | 4 | 25.1 | 3.7 | 5 | 20.8 | 4.0 | 4 |
| **Q7JXB5** | 211 | Phosphoenolpyruvate carboxykinase 2 (RE12569p) | | | | | | | | | | | | | | | | 14.9 | 6.5 | 3 |
| **Q7K1C0** | 17 | NADH dehydrogenase (ubiquinone) 15 kDa subunit (MIP04232p) | | | | 12.9 | 1.4 | 3 | | | | 10.9 | 2.0 | 4 | 22.7 | 5.6 | 4 | | | |
| **Q7K1Q6** | 172 | Jabba, isoform B (LD47410p) | | | | | | | | | | | | | 11.3 | 3.3 | 5 | | | |
| **Q7K4T8** | 295 | uncharacterized protein CG8520 (LD23856p) | | | | 5.7 | 1.5 | 3 | | | | | | | 10.1 | 2.0 | 5 | | | |
| **Q7K569** | 78 | Glycerophosphate oxidase-1; isoform D | | | | | | | | | | | | | 22.2 | 4.7 | 4 | 20.1 | 2.0 | 3 |
| **Q7K569** | 207 | Glycerophosphate oxidase-1; isoform D | | | | | | | | | | 13.6 | 0.9 | 4 | 19.7 | 4.6 | 5 | 19.1 | 7.4 | 4 |
| **Q7K569** | 223 | Glycerophosphate oxidase-1; isoform D | 12.8 | 1.7 | 5 | 9.3 | 2.2 | 5 | 12.9 | 4.9 | 5 | 13.1 | 0.5 | 5 | 18.6 | 3.9 | 5 | 17.6 | 6.8 | 5 |
| **Q7K569** | 238 | Glycerophosphate oxidase-1; isoform D | 18.3 | 13.4 | 5 | 10.2 | 3.4 | 5 | 12.7 | 6.4 | 5 | 12.9 | 1.0 | 5 | 20.4 | 3.6 | 5 | 19.4 | 6.5 | 5 |
| **Q7K569** | 259 | Glycerophosphate oxidase-1; isoform D | 14.0 | 2.2 | 5 | 10.0 | 2.6 | 5 | 12.8 | 4.7 | 5 | 13.4 | 0.3 | 5 | 18.0 | 4.7 | 5 | 18.8 | 6.3 | 5 |
| **Q7K569** | 278 | Glycerophosphate oxidase-1; isoform D | 15.5 | 2.5 | 5 | 14.2 | 0.7 | 3 | 18.5 | 3.4 | 3 | 16.5 | 0.4 | 5 | 20.3 | 3.0 | 5 | 22.1 | 7.8 | 5 |
| **Q7K569** | 352 | Glycerophosphate oxidase-1; isoform D | | | | | | | | | | 11.8 | 0.4 | 4 | 18.6 | 1.9 | 4 | 14.9 | 4.6 | 4 |
| **Q7K569** | 448 | Glycerophosphate oxidase-1; isoform D | 6.4 | 1.5 | 5 | 6.4 | 1.5 | 5 | 7.4 | 4.2 | 5 | 6.9 | 0.2 | 5 | 13.4 | 2.8 | 5 | 11.3 | 3.6 | 5 |
| **Q7K569** | 486 | Glycerophosphate oxidase-1; isoform D | 9.4 | 2.7 | 5 | 9.1 | 1.8 | 5 | 9.7 | 4.6 | 5 | 9.0 | 0.1 | 5 | 16.6 | 3.1 | 5 | 15.6 | 5.6 | 5 |
| **Q7K569** | 541 | Glycerophosphate oxidase-1; isoform D | | | | | | | | | | 5.4 | 0.3 | 3 | 10.8 | 3.1 | 4 | | | |
| **Q7K5K3** | 302 | Pyruvate dehydrogenase E1 subunit beta, mitochondrial (GH08474p) | 17.5 | 5.2 | 5 | 14.4 | 9.3 | 4 | 18.4 | 3.6 | 5 | 13.4 | 2.6 | 5 | 18.3 | 8.5 | 5 | | | |
| **Q7K5K3** | 321 | Pyruvate dehydrogenase E1 subunit beta, mitochondrial (GH08474p) | 14.1 | 1.7 | 5 | 11.4 | 4.4 | 5 | 10.8 | 3.7 | 5 | 15.2 | 1.4 | 5 | 20.3 | 2.4 | 5 | 18.3 | 5.2 | 5 |
| **Q7K860** | 71 | Troponin C isoform 4; isoform B | 5.7 | 2.7 | 5 | 6.6 | 2.3 | 5 | 8.2 | 7.7 | 5 | 6.3 | 1.3 | 5 | 12.3 | 4.4 | 5 | 10.1 | 3.7 | 5 |
| **Q7KN62** | 569 | Transitional endoplasmic reticulum ATPase TER94 | | | | | | | 12.7 | 6.2 | 3 | 12.7 | 1.3 | 5 | 18.7 | 2.8 | 5 | 15.9 | 4.1 | 5 |
| **Q7KN75** | 1042 | Dodeca-satellite-binding protein 1; isoform F | | | | | | | | | | | | | 26.6 | 19.3 | 4 | | | |
| **Q7KN97** | 378 | Pyruvate carboxylase | 8.2 | 1.6 | 4 | 5.7 | 1.7 | 5 | 7.7 | 1.6 | 4 | 8.3 | 0.4 | 4 | 13.1 | 2.6 | 3 | 14.5 | 1.7 | 3 |
| **Q7KN97** | 550 | Pyruvate carboxylase | | | | | | | | | | | | | 12.3 | 2.9 | 4 | 11.7 | 3.4 | 4 |
| **Q7KN97** | 853 | Pyruvate carboxylase | 13.2 | 2.4 | 3 | 8.7 | 0.3 | 3 | 13.0 | 4.3 | 3 | 14.5 | 2.4 | 5 | 18.3 | 2.4 | 5 | 17.5 | 5.1 | 5 |
| **Q7KN97** | 1003 | Pyruvate carboxylase | | | | | | | | | | 9.3 | 0.6 | 4 | 13.9 | 4.0 | 4 | 14.5 | 2.5 | 3 |
| **Q7KR04** | 13 | 40S ribosomal protein S15Ab | | | | | | | | | | | | | | | | 15.0 | 3.0 | 3 |
| **Q7KR04** | 72 | 40S ribosomal protein S15Ab | | | | | | | | | | | | | 19.6 | 5.9 | 5 | | | |
| **Q7KSQ0** | 78 | Citrate transport protein (RH27308p) | | | | | | | 9.5 | 4.9 | 3 | | | | | | | | | |
| **Q7KSU6** | 183 | Transketolase (CG8036; isoform D) | | | | 7.4 | 1.0 | 3 | | | | | | | | | | | | |
| **Q7KTK9** | 224 | Dihydrolipoamide S-acetyltransferase (CG5261; isoform A) | | | | | | | 10.5 | 2.3 | 3 | 6.7 | 2.2 | 4 | 9.8 | 2.7 | 5 | 13.4 | 4.7 | 3 |
| **Q7KTW5** | 37 | Inositol-1-monophosphatase (RE10407p) | | | | | | | 12.6 | 2.9 | 4 | | | | | | | 22.4 | 5.0 | 3 |
| **Q7KUB0** | 60 | Isocitrate dehydrogenase [NADP] | 10.4 | 2.2 | 4 | 9.0 | 2.6 | 5 | 10.3 | 4.1 | 5 | 11.9 | 0.8 | 5 | 15.5 | 2.8 | 5 | 19.0 | 7.1 | 5 |
| **Q7KUB0** | 73 | Isocitrate dehydrogenase [NADP] | | | | 12.8 | 2.8 | 3 | | | | 18.2 | 3.5 | 4 | | | | | | |
| **Q7KUB0** | 377 | Isocitrate dehydrogenase [NADP] | | | | | | | | | | | | | 11.3 | 4.5 | 4 | | | |
| **Q7KUQ5** | 33 | Limpet; isoform F | 36.0 | 3.6 | 5 | 34.8 | 4.0 | 5 | 36.0 | 4.9 | 5 | 41.6 | 0.7 | 5 | 39.0 | 5.8 | 4 | 37.1 | 6.2 | 5 |
| **Q7KV27** | 144 | Alanine transaminase (CG1640; isoform F) | | | | | | | | | | 12.4 | 2.0 | 4 | 19.0 | 4.6 | 5 | 14.3 | 6.2 | 3 |
| **Q7KV27** | 250 | Alanine transaminase (CG1640; isoform F) | | | | | | | | | | | | | 18.0 | 2.6 | 3 | | | |
| **Q7KV27** | 392 | Alanine transaminase (CG1640; isoform F) | | | | | | | | | | 7.1 | 0.6 | 3 | 9.1 | 0.2 | 3 | 9.3 | 3.7 | 3 |
| **Q7KVP4** | 152 | Glycogen debranching enzyme (CG9485; isoform E) | | | | | | | | | | 6.7 | 0.5 | 4 | 13.2 | 3.0 | 4 | | | |
| **Q7KVP4** | 354 | Glycogen debranching enzyme (CG9485; isoform E) | 8.9 | 1.8 | 5 | 8.0 | 2.1 | 5 | 7.5 | 2.7 | 5 | 7.9 | 1.3 | 5 | 12.1 | 2.1 | 4 | 11.4 | 3.5 | 4 |
| **Q7KVP4** | 928 | Glycogen debranching enzyme (CG9485; isoform E) | | | | | | | | | | | | | 10.3 | 4.4 | 3 | | | |
| **Q7KVP4** | 976 | Glycogen debranching enzyme (CG9485; isoform E) | | | | | | | | | | 9.2 | 1.2 | 4 | 17.3 | 6.6 | 5 | 9.4 | 3.3 | 4 |
| **Q7YU05** | 114 | Pyruvate dehydrogenase E1 alpha subunit (Lethal (1) G0334; isoform B) | | | | | | | | | | 25.0 | 0.7 | 4 | 28.2 | 3.8 | 4 | 29.5 | 11.5 | 3 |
| **Q7YU05** | 155 | Pyruvate dehydrogenase E1 alpha subunit (Lethal (1) G0334; isoform B) | | | | | | | | | | 16.5 | 2.9 | 4 | | | | | | |
| **Q7YU05** | 171 | Pyruvate dehydrogenase E1 alpha subunit (Lethal (1) G0334; isoform B) | 11.0 | 1.3 | 5 | 9.7 | 2.4 | 5 | 10.9 | 3.8 | 5 | 11.8 | 1.7 | 5 | 17.6 | 2.4 | 5 | 17.3 | 5.7 | 5 |
| **Q868Z9** | 2824 | Papilin | | | | | | | 75.9 | 8.2 | 3 | | | | | | | | | |
| **Q86BQ4** | 38 | Histidine triad nucleotide binding protein 1 (CG2862; isoform B) | 29.9 | 5.1 | 4 | | | | 23.1 | 7.4 | 5 | | | | 33.7 | 4.4 | 4 | 28.3 | 9.3 | 4 |
| **Q8I0E9** | 460 | Heat shock protein 70 cognate 1; isoform D | | | | | | | | | | | | | | | | | | |
| **Q8I0E9** | 517 | Heat shock protein 70 cognate 1; isoform D | | | | | | | | | | | | | 13.1 | 2.3 | 4 | | | |
| **Q8INE6** | 54 | Oligomycin sensitivity-conferring protein; isoform B | 7.6 | 2.0 | 5 | 6.0 | 1.2 | 5 | 8.8 | 4.2 | 5 | 8.5 | 1.3 | 5 | 12.4 | 3.7 | 5 | 16.0 | 7.7 | 5 |
| **Q8IPE8** | 322 | Mitochondrial trifunctional protein alpha subunit (CG4389; isoform C) | | | | 8.2 | 2.1 | 4 | 8.1 | 3.3 | 5 | | | | | | | | | |
| **Q8IPG9** | 51 | Basigin (LD19437p) | | | | | | | 67.3 | 15.1 | 5 | 65.4 | 5.2 | 5 | 87.8 | 2.6 | 4 | | | |
| **Q8IPP8** | 181 | uncharacterized protein CG31548 | 9.8 | 4.2 | 3 | | | | | | | 9.9 | 0.4 | 4 | 15.9 | 2.5 | 4 | 14.4 | 4.6 | 4 |
| **Q8IQ53** | 29 | Ubiquitin-conjugating enzyme variant 1A; isoform B | | | | | | | | | | 12.4 | 2.2 | 3 | 15.2 | 0.6 | 3 | | | |

| | | | | | | | | | | | | | | | | | | | | |
|---|---|---|---|---|---|---|---|---|---|---|---|---|---|---|---|---|---|---|---|---|
| **Q8IQQ0** | 160 | Oxoglutarate dehydrogenase, mitochondrial (RE42354p) | 9.0 | 1.8 | 3 | | | | | | | | | | | | | | | |
| **Q8IQQ0** | 219 | Oxoglutarate dehydrogenase, mitochondrial (RE42354p) | | | | | | | | | | | | | 8.0 | 3.0 | 3 | | | |
| **Q8IQQ0** | 287 | Oxoglutarate dehydrogenase, mitochondrial (RE42354p) | | | | | | | | | | | | | 12.3 | 3.9 | 3 | | | |
| **Q8IQQ0** | 326 | Oxoglutarate dehydrogenase, mitochondrial (RE42354p) | 5.0 | 1.4 | 5 | 6.0 | 1.7 | 5 | 5.5 | 3.5 | 5 | 6.4 | 0.8 | 5 | 13.3 | 3.4 | 5 | 7.8 | 2.2 | 5 |
| **Q8IQQ0** | 464 | Oxoglutarate dehydrogenase, mitochondrial (RE42354p) | 11.2 | 2.0 | 5 | 10.5 | 2.1 | 5 | 10.0 | 3.0 | 4 | 11.4 | 0.6 | 5 | 18.9 | 2.8 | 5 | 15.1 | 3.2 | 5 |
| **Q8IQQ0** | 490 | Oxoglutarate dehydrogenase, mitochondrial (RE42354p) | | | | | | | | | | 10.6 | 0.7 | 4 | 14.1 | 4.4 | 5 | 14.6 | 6.2 | 4 |
| **Q8IQQ0** | 504 | Oxoglutarate dehydrogenase, mitochondrial (RE42354p) | 10.9 | 1.5 | 3 | | | | | | | 7.6 | 0.3 | 4 | 13.3 | 3.6 | 5 | 13.0 | 3.9 | 5 |
| **Q8IQQ0** | 539 | Oxoglutarate dehydrogenase, mitochondrial (RE42354p) | 14.5 | 2.6 | 5 | | | | 12.8 | 2.1 | 4 | 11.1 | 1.0 | 5 | 17.1 | 4.2 | 5 | 15.8 | 5.1 | 5 |
| **Q8IQQ0** | 569 | Oxoglutarate dehydrogenase, mitochondrial (RE42354p) | | | | 7.5 | 3.1 | 4 | 9.7 | 4.6 | 3 | 7.0 | 1.0 | 5 | 11.6 | 3.4 | 5 | 11.9 | 5.5 | 5 |
| **Q8IQQ0** | 795 | Oxoglutarate dehydrogenase, mitochondrial (RE42354p) | | | | | | | | | | 15.6 | 1.2 | 3 | | | | | | |
| **Q8IQW2** | 24 | Cytochrome c oxidase subunit 6B (RE29690p) | 89.8 | 3.5 | 5 | 92.2 | 2.5 | 5 | 92.7 | 3.9 | 5 | 89.7 | 4.0 | 5 | 96.9 | 1.4 | 5 | 96.3 | 2.0 | 5 |
| **Q8IQW2** | 45 | Cytochrome c oxidase subunit 6B (RE29690p) | 89.2 | 4.1 | 5 | 91.5 | 2.5 | 5 | 91.6 | 3.7 | 5 | 88.0 | 7.7 | 5 | 96.0 | 1.1 | 5 | 93.8 | 3.4 | 5 |
| **Q8IR13** | 87 | Eukaryotic translation initiation factor 4H1 (RNA-binding protein 2; isoform B) | | | | | | | | | | | | | 18.8 | 6.8 | 3 | | | |
| **Q8IRQ5** | 222 | Fumarate hydratase 1 (Lethal (1) G0255; isoform B) | 5.1 | 1.1 | 5 | 4.7 | 1.2 | 5 | 5.6 | 3.1 | 5 | 5.9 | 0.6 | 5 | 10.6 | 1.9 | 5 | 9.2 | 3.6 | 5 |
| **Q8IRQ5** | 391 | Fumarate hydratase 1 (Lethal (1) G0255; isoform B) | 7.4 | 1.9 | 5 | 7.4 | 1.8 | 5 | 7.4 | 3.5 | 5 | 8.7 | 0.9 | 5 | 14.4 | 2.6 | 5 | 12.6 | 3.7 | 5 |
| **Q8MRK8** | 205 | uncharacterized protein CG12136 (GH26280p) | 9.0 | 1.5 | 5 | 8.3 | 1.7 | 4 | | | | | | | | | | | | |
| **Q8MSI2** | 32 | Sarcoplasmic calcium-binding protein 1 | 14.2 | 2.6 | 5 | 13.2 | 2.4 | 5 | 16.0 | 6.9 | 5 | 15.1 | 1.1 | 5 | 21.7 | 5.1 | 5 | 24.6 | 8.2 | 5 |
| **Q8MSI2** | 38 | Sarcoplasmic calcium-binding protein 1 | 14.6 | 2.3 | 5 | 13.2 | 3.1 | 5 | 18.4 | 8.9 | 5 | 18.6 | 2.4 | 5 | 27.0 | 5.2 | 4 | 25.5 | 8.4 | 5 |
| **Q8MSI2** | 46 | Sarcoplasmic calcium-binding protein 1 | 14.9 | 2.5 | 5 | 15.5 | 4.0 | 5 | 15.9 | 7.8 | 5 | | | | | | | 23.7 | 7.9 | 4 |
| **Q8MSI2** | 90 | Sarcoplasmic calcium-binding protein 1 | 11.6 | 2.2 | 5 | 11.4 | 3.4 | 5 | 11.4 | 4.8 | 5 | | | | 21.2 | 3.4 | 3 | 20.4 | 0.2 | 3 |
| **Q8MSI2** | 130 | Sarcoplasmic calcium-binding protein 1 | 11.5 | 2.9 | 5 | 9.4 | 3.0 | 5 | 12.4 | 6.8 | 5 | | | | | | | 25.8 | 7.3 | 3 |
| **Q8MST5** | 12 | beta-Tubulin at 97EF (LP10436p) | | | | | | | | | | | | | 18.6 | 3.2 | 4 | 18.3 | 8.0 | 4 |
| **Q8MST5** | 303 | beta-Tubulin at 97EF (LP10436p) | | | | | | | | | | 9.9 | 1.0 | 4 | 14.0 | 3.0 | 3 | | | |
| **Q8SY61** | 35 | General odorant-binding protein 56d | | | | 75.9 | 7.1 | 3 | 80.4 | 10.6 | 3 | 67.0 | 17.4 | 4 | 90.3 | 1.1 | 4 | 77.2 | 8.6 | 3 |
| **Q8SY69** | 64 | MICOS subunit 10b (RH61753p) | 37.9 | 5.5 | 5 | 33.2 | 5.7 | 4 | 35.4 | 8.5 | 5 | 41.7 | 5.2 | 5 | 51.7 | 13.5 | 5 | | | |
| **Q8T8R1** | 10 | CCHC-type zinc finger nucleic acid binding protein | | | | 47.4 | 6.5 | 4 | | | | | | | | | | 48.8 | 8.4 | 4 |
| **Q8T8R1** | 20 | CCHC-type zinc finger nucleic acid binding protein | 32.4 | 4.1 | 4 | | | | | | | | | | | | | | | |
| **Q8T8R1** | 60 | CCHC-type zinc finger nucleic acid binding protein | 58.6 | 4.7 | 4 | | | | | | | 51.0 | 0.7 | 3 | 56.8 | 6.3 | 3 | | | |
| **Q8T8R1** | 70 | CCHC-type zinc finger nucleic acid binding protein | 45.7 | 3.6 | 5 | 45.2 | 6.5 | 5 | 42.8 | 7.1 | 5 | 47.7 | 2.3 | 5 | 52.2 | 4.7 | 4 | 41.8 | 10.5 | 3 |
| **Q8T8R1** | 80 | CCHC-type zinc finger nucleic acid binding protein | | | | | | | 48.2 | 6.4 | 4 | 51.0 | 3.3 | 5 | 53.8 | 5.2 | 5 | 47.4 | 6.6 | 3 |
| **Q8T8R1** | 111 | CCHC-type zinc finger nucleic acid binding protein | | | | | | | | | | 35.3 | 0.7 | 3 | | | | | | |
| **Q8T8R1** | 125 | CCHC-type zinc finger nucleic acid binding protein | | | | 46.0 | 3.1 | 5 | 48.3 | 8.2 | 5 | 51.3 | 3.3 | 5 | 53.5 | 4.7 | 5 | 51.9 | 6.1 | 5 |
| **Q94511** | 85 | NADH dehydrogenase (ubiquinone) 75 kDa subunit | 29.9 | 2.3 | 5 | 23.8 | 7.0 | 5 | 26.6 | 7.9 | 5 | 32.3 | 2.1 | 5 | 32.9 | 5.7 | 5 | 33.4 | 8.6 | 5 |
| **Q94511** | 88 | NADH dehydrogenase (ubiquinone) 75 kDa subunit | 25.8 | 2.4 | 5 | | | | | | | | | | | | | | | |
| **Q94511** | 102 | NADH dehydrogenase (ubiquinone) 75 kDa subunit | | | | 14.2 | 4.8 | 4 | | | | 16.3 | 1.7 | 5 | 22.8 | 4.0 | 5 | 20.0 | 5.1 | 4 |
| **Q94511** | 308 | NADH dehydrogenase (ubiquinone) 75 kDa subunit | 10.7 | 2.3 | 5 | 10.0 | 2.0 | 5 | 9.6 | 4.0 | 5 | 11.3 | 0.9 | 5 | 16.7 | 2.0 | 5 | 15.3 | 3.9 | 5 |
| **Q94511** | 467 | NADH dehydrogenase (ubiquinone) 75 kDa subunit | 9.5 | 1.6 | 5 | 8.2 | 2.3 | 5 | 8.1 | 3.8 | 5 | 9.9 | 0.7 | 5 | 14.1 | 2.2 | 5 | 12.6 | 3.6 | 5 |
| **Q94511** | 508 | NADH dehydrogenase (ubiquinone) 75 kDa subunit | 4.0 | 1.7 | 3 | 3.9 | 0.6 | 5 | 6.4 | 5.5 | 4 | 3.6 | 0.7 | 5 | 9.1 | 2.5 | 5 | 7.4 | 2.6 | 5 |
| **Q94511** | 570 | NADH dehydrogenase (ubiquinone) 75 kDa subunit | | | | | | | | | | 8.7 | 1.0 | 4 | 12.7 | 2.6 | 5 | 16.9 | 3.9 | 4 |
| **Q94511** | 712 | NADH dehydrogenase (ubiquinone) 75 kDa subunit | 8.4 | 2.0 | 5 | 7.2 | 2.8 | 5 | 8.6 | 4.4 | 5 | 9.7 | 0.5 | 5 | 15.3 | 3.4 | 5 | 15.0 | 5.2 | 5 |
| **Q94514** | 100 | Cytochrome c oxidase subunit 5A; mitochondrial | 24.1 | 5.2 | 5 | 23.4 | 6.0 | 5 | 21.7 | 5.2 | 5 | 21.4 | 1.6 | 5 | 30.5 | 6.4 | 5 | 26.7 | 6.2 | 5 |
| **Q94516** | 193 | ATP synthase subunit b, mitochondrial | 6.3 | 1.5 | 5 | 5.5 | 1.6 | 5 | 6.4 | 2.8 | 5 | 6.6 | 0.6 | 5 | 11.6 | 2.7 | 5 | 12.0 | 4.6 | 5 |
| **Q94516** | 229 | ATP synthase subunit b, mitochondrial | 5.4 | 1.8 | 5 | 5.6 | 1.7 | 5 | 6.6 | 4.4 | 5 | 6.3 | 0.6 | 5 | 12.2 | 2.8 | 5 | 11.6 | 4.2 | 5 |
| **Q94522** | 42 | Succinyl-CoA ligase [ADP/GDP-forming] subunit alpha; mitochondrial | 19.7 | 2.0 | 5 | 16.6 | 4.0 | 5 | 18.9 | 6.6 | 5 | 19.7 | 1.6 | 5 | 24.8 | 4.7 | 5 | 27.2 | 6.1 | 5 |
| **Q960M4** | 79 | Peroxiredoxin-5 (RE23139p) | 50.9 | 2.5 | 5 | 49.0 | 6.3 | 5 | 43.9 | 5.0 | 5 | 45.6 | 3.4 | 3 | 53.5 | 7.3 | 5 | 49.5 | 8.4 | 5 |
| **Q960M4** | 181 | Peroxiredoxin-5 (RE23139p) | 33.8 | 2.1 | 5 | 32.5 | 5.1 | 5 | 28.8 | 4.6 | 5 | 32.2 | 2.8 | 5 | 38.3 | 6.9 | 5 | 38.3 | 6.3 | 5 |
| **Q9GNH8** | 439 | Hexokinase A | 7.7 | 2.4 | 3 | | | | | | | | | | | | | | | |
| **Q9GU68** | 74 | Eukaryotic translation initiation factor 5A | 22.5 | 4.8 | 3 | 16.3 | 2.4 | 3 | 21.6 | 6.1 | 3 | 18.1 | 0.9 | 4 | 27.5 | 3.5 | 5 | 26.7 | 6.5 | 5 |
| **Q9TVP3** | 70 | J domain-containing protein | | | | | | | | | | | | | 15.0 | 2.8 | 5 | 14.4 | 2.8 | 4 |
| **Q9TVP3** | 146 | J domain-containing protein | | | | | | | | | | | | | 25.5 | 5.7 | 3 | | | |
| **Q9V3P0** | 168 | Peroxiredoxin 2 | | | | | | | 16.3 | 5.4 | 5 | | | | 26.5 | 3.0 | 4 | | | |
| **Q9V3W0** | 109 | RutC family protein UK114 | | | | | | | | | | | | | | | | 18.2 | 6.6 | 3 |
| **Q9V3W2** | 78 | NADH dehydrogenase (ubiquinone) B17 subunit (Lethal (2) 35Di; isoform A) | | | | | | | | | | 9.8 | 0.5 | 4 | 16.9 | 3.3 | 5 | | | |
| **Q9V3W2** | 136 | NADH dehydrogenase (ubiquinone) B17 subunit (Lethal (2) 35Di; isoform A) | 7.8 | 1.7 | 5 | 7.7 | 2.0 | 5 | 7.7 | 3.6 | 5 | 9.4 | 0.9 | 5 | 15.3 | 3.7 | 5 | 14.3 | 4.5 | 5 |
| **Q9V427** | 322 | Innexin 2 | | | | | | | | | | | | | 14.9 | 7.4 | 3 | | | |
| **Q9V496** | 88 | Apolipophorins | | | | 66.7 | 3.9 | 4 | | | | 72.6 | 0.5 | 4 | 85.4 | 7.5 | 3 | 81.9 | 9.9 | 3 |
| **Q9V496** | 168 | Apolipophorins | 68.4 | 5.5 | 5 | 65.4 | 7.0 | 5 | 63.3 | 10.4 | 5 | 60.0 | 0.6 | 5 | 84.1 | 9.9 | 5 | 84.0 | 8.3 | 5 |
| **Q9V496** | 192 | Apolipophorins | 73.5 | 5.0 | 5 | 68.2 | 8.9 | 5 | 68.7 | 10.1 | 5 | 62.3 | 2.5 | 5 | 90.7 | 3.3 | 4 | 86.1 | 6.7 | 4 |
| **Q9V496** | 435 | Apolipophorins | | | | | | | | | | 51.9 | 13.7 | 4 | 80.5 | 4.5 | 3 | | | |
| **Q9V496** | 454 | Apolipophorins | 74.5 | 7.4 | 5 | 71.1 | 9.3 | 5 | 68.1 | 12.8 | 5 | 56.2 | 8.9 | 4 | | | | | | |
| **Q9V496** | 489 | Apolipophorins | 77.3 | 5.3 | 5 | 73.0 | 7.6 | 5 | 74.0 | 9.7 | 5 | 67.8 | 0.9 | 5 | 88.7 | 6.8 | 5 | 87.1 | 6.4 | 3 |
| **Q9V496** | 511 | Apolipophorins | 55.4 | 9.2 | 5 | 65.6 | 11.0 | 4 | 69.3 | 5.5 | 3 | | | | 77.2 | 5.6 | 4 | | | |
| **Q9V496** | 2048 | Apolipophorins | | | | | | | | | | 62.6 | 4.0 | 5 | 78.9 | 4.1 | 4 | 67.6 | 11.8 | 3 |
| **Q9V496** | 2810 | Apolipophorins | 66.8 | 4.4 | 5 | 57.7 | 6.9 | 5 | 62.4 | 7.6 | 4 | 54.5 | 1.1 | 5 | 79.9 | 5.4 | 5 | 75.3 | 8.9 | 5 |
| **Q9V496** | 2949 | Apolipophorins | 69.2 | 7.2 | 5 | 62.4 | 8.3 | 5 | 61.1 | 9.8 | 5 | 59.5 | 2.1 | 5 | 84.2 | 4.2 | 5 | 80.6 | 7.8 | 5 |
| **Q9V496** | 2994 | Apolipophorins | 65.7 | 9.1 | 5 | 67.3 | 10.2 | 5 | 60.3 | 16.4 | 5 | 58.9 | 6.8 | 5 | 89.2 | 2.9 | 5 | 83.6 | 9.7 | 5 |
| **Q9V496** | 3037 | Apolipophorins | | | | 79.0 | 3.0 | 3 | | | | 82.0 | 8.0 | 4 | | | | 82.0 | 10.6 | 3 |
| **Q9V496** | 3321 | Apolipophorins | 67.9 | 7.7 | 4 | 62.1 | 8.8 | 4 | 63.7 | 10.7 | 5 | 66.0 | 5.8 | 5 | 85.4 | 1.3 | 4 | 79.8 | 9.7 | 4 |
| **Q9V496** | 3323 | Apolipophorins | 78.3 | 5.7 | 5 | 72.8 | 9.7 | 5 | 72.6 | 10.9 | 5 | 70.3 | 5.9 | 5 | 91.9 | 2.9 | 5 | 88.3 | 7.3 | 5 |
| **Q9V4E0** | 352 | NADH dehydrogenase (ubiquinone) 49 kDa subunit (LD47962p) | 6.5 | 1.2 | 5 | 5.6 | 1.5 | 5 | 7.8 | 2.9 | 4 | 7.9 | 0.9 | 5 | 12.6 | 2.1 | 5 | 12.2 | 4.3 | 5 |
| **Q9V6U9** | 242 | Mitochondrial trans-2-enoyl-CoA reductase | | | | | | | | | | 9.4 | 1.8 | 4 | 13.5 | 2.2 | 5 | 14.8 | 5.5 | 4 |
| **Q9V8M5** | 64 | Probable 3-hydroxyisobutyrate dehydrogenase, mitochondrial | | | | | | | | | | 11.4 | 1.0 | 4 | 18.8 | 5.3 | 5 | 16.7 | 5.0 | 5 |
| **Q9VAC1** | 57 | uncharacterized protein CG7920 (GM14349p) | 11.7 | 2.2 | 5 | 9.7 | 2.3 | 5 | 11.7 | 4.5 | 5 | 13.0 | 0.7 | 5 | 18.1 | 3.1 | 5 | 20.3 | 6.6 | 5 |
| **Q9VAJ4** | 125 | General odorant-binding protein 99a | | | | 94.6 | 2.6 | 3 | | | | | | | | | | | | |
| **Q9VAM6** | 100 | CDGSH iron-sulfur domain | 29.5 | 3.5 | 5 | 29.0 | 4.3 | 5 | 29.6 | 6.7 | 5 | | | | | | | | | |
| **Q9VAM6** | 111 | CDGSH iron-sulfur domain | | | | | | | | | | | | | 40.9 | 7.6 | 4 | | | |
| **Q9VAN7** | 24 | Phosphoglyceromutase 78; isoform C | | | | | | | | | | | | | 17.5 | 6.9 | 4 | | | |
| **Q9VB69** | 96 | Malic enzyme b | 9.7 | 2.1 | 5 | 7.2 | 2.8 | 5 | 9.5 | 4.7 | 5 | 9.5 | 0.7 | 5 | 14.3 | 2.9 | 5 | 16.6 | 6.0 | 5 |
| **Q9VB69** | 203 | Malic enzyme b | | | | | | | | | | 16.2 | 1.1 | 3 | 18.1 | 2.1 | 4 | | | |
| **Q9VB69** | 455 | Malic enzyme b | 8.4 | 2.7 | 5 | 7.4 | 2.1 | 3 | | | | | | | 14.1 | 1.7 | 4 | 14.8 | 5.2 | 4 |
| **Q9VBP6** | 243 | Succinic semialdehyde dehydrogenase; isoform E | | | | | | | | | | | | | 16.9 | 5.6 | 3 | | | |
| **Q9VBU9** | 77 | 40S ribosomal protein S27 | 12.9 | 2.2 | 5 | 14.2 | 3.8 | 5 | | | | 15.5 | 2.9 | 4 | 23.3 | 1.8 | 5 | 22.7 | 2.4 | 3 |
| **Q9VCD9** | 109 | uncharacterized protein CG6000; isoform A | 34.4 | 6.3 | 5 | 26.3 | 6.9 | 5 | 25.2 | 10.6 | 4 | 30.3 | 1.3 | 5 | 34.5 | 5.0 | 5 | 38.7 | 8.6 | 5 |
| **Q9VCI3** | 82 | Lipid storage droplets surface-binding protein 1 | | | | 24.5 | 11.0 | 3 | 17.8 | 7.3 | 3 | 9.5 | 0.0 | 4 | 17.8 | 6.9 | 5 | 14.5 | 4.4 | 4 |
| **Q9VD58** | 40 | Isocitrate dehydrogenase 3b (GH26270p) | 15.6 | 2.1 | 5 | 12.2 | 2.4 | 5 | 14.4 | 5.1 | 5 | 15.1 | 0.7 | 5 | 19.8 | 3.3 | 5 | 21.8 | 5.7 | 5 |
| **Q9VD58** | 71 | Isocitrate dehydrogenase 3b (GH26270p) | | | | | | | 9.6 | 3.7 | 4 | 9.4 | 0.7 | 4 | 14.4 | 2.6 | 5 | 12.2 | 4.8 | 4 |
| **Q9VD58** | 222 | Isocitrate dehydrogenase 3b (GH26270p) | 5.8 | 1.7 | 5 | 6.3 | 1.5 | 5 | 6.1 | 3.8 | 5 | 7.6 | 0.8 | 5 | 20.1 | 11.9 | 5 | 11.4 | 3.5 | 5 |
| **Q9VD58** | 317 | Isocitrate dehydrogenase 3b (GH26270p) | | | | 5.9 | 1.6 | 3 | | | | 4.4 | 0.3 | 4 | 7.6 | 2.3 | 5 | 6.1 | 3.4 | 3 |
| **Q9VEB1** | 30 | Malate dehydrogenase 2, mitochondrial (RE60471p) | 16.3 | 3.1 | 5 | 11.9 | 3.3 | 5 | 13.4 | 4.6 | 5 | 16.3 | 1.3 | 5 | 19.6 | 4.5 | 5 | 19.6 | 5.7 | 5 |
| **Q9VEB1** | 132 | Malate dehydrogenase 2, mitochondrial (RE60471p) | 8.5 | 2.8 | 5 | 10.4 | 2.9 | 5 | 8.8 | 5.5 | 5 | | | | | | | 15.7 | 1.2 | 3 |
| **Q9VEB1** | 147 | Malate dehydrogenase 2, mitochondrial (RE60471p) | 8.0 | 3.2 | 5 | 10.0 | 2.5 | 5 | 8.2 | 5.1 | 5 | 10.2 | 1.7 | 5 | 17.7 | 3.6 | 5 | 12.0 | 3.6 | 5 |
| **Q9VEB1** | 275 | Malate dehydrogenase 2, mitochondrial (RE60471p) | | | | | | | | | | | | | 14.3 | 3.7 | 4 | | | |
| **Q9VEJ0** | 206 | Peroxiredoxin 3 (Thioredoxin peroxidase 3) | 25.9 | 5.4 | 5 | 28.9 | 6.6 | 5 | 24.9 | 9.5 | 4 | 22.8 | 3.6 | 5 | 33.7 | 5.8 | 5 | | | |
| **Q9VEN1** | 1604 | Filamin A | 16.4 | 2.9 | 3 | | | | | | | | | | | | | | | |
| **Q9VEN1** | 1726 | Filamin A | 15.7 | 1.8 | 5 | 12.1 | 1.1 | 3 | 12.9 | 4.5 | 4 | 15.8 | 0.6 | 5 | 18.2 | 1.5 | 5 | 17.8 | 3.9 | 4 |
| **Q9VEX6** | 564 | Belphegor | | | | | | | | | | | | | 10.2 | 1.3 | 4 | 16.7 | 4.3 | 4 |
| **Q9VF24** | 184 | crossveinless d (GH05619p) | | | | 61.5 | 9.9 | 3 | | | | | | | | | | | | |
| **Q9VF27** | 197 | NADH dehydrogenase (ubiquinone) 23 kDa subunit | | | | | | | | | | | | | 18.7 | 3.4 | 5 | | | |
| **Q9VFC8** | 462 | Glycogen synthase | 16.7 | 2.2 | 5 | 13.9 | 2.0 | 3 | 15.6 | 4.4 | 4 | 15.5 | 1.2 | 5 | 24.9 | 4.1 | 5 | 20.8 | 5.0 | 5 |
| **Q9VFC8** | 494 | Glycogen synthase | | | | | | | | | | 16.5 | 0.7 | 4 | 22.6 | 6.3 | 5 | 20.6 | 8.6 | 3 |
| **Q9VFF0** | 128 | Ubiquinol-cytochrome c reductase core protein 1 (GH01077p) | 8.8 | 1.8 | 5 | 6.9 | 2.0 | 5 | 8.7 | 4.0 | 5 | | | | 12.8 | 4.0 | 4 | 13.2 | 5.0 | 5 |
| **Q9VFF0** | 370 | Ubiquinol-cytochrome c reductase core protein 1 (GH01077p) | 5.7 | 1.8 | 5 | 5.7 | 1.4 | 5 | 7.6 | 4.3 | 4 | 6.3 | 0.8 | 5 | 10.9 | 2.0 | 5 | 9.9 | 4.1 | 5 |
| **Q9VFF0** | 400 | Ubiquinol-cytochrome c reductase core protein 1 (GH01077p) | 12.9 | 2.9 | 5 | 14.8 | 3.2 | 5 | 14.3 | 5.9 | 4 | 16.5 | 1.4 | 5 | 22.5 | 2.0 | 5 | 17.5 | 4.4 | 5 |
| **Q9VFF0** | 409 | Ubiquinol-cytochrome c reductase core protein 1 (GH01077p) | 12.2 | 2.7 | 5 | 11.9 | 2.3 | 5 | 11.8 | 6.0 | 5 | 14.1 | 1.3 | 5 | 17.6 | 2.8 | 5 | 15.0 | 4.1 | 5 |
| **Q9VFF0** | 443 | Ubiquinol-cytochrome c reductase core protein 1 (GH01077p) | 12.4 | 4.0 | 5 | 15.2 | 3.5 | 5 | 12.8 | 7.6 | 5 | 14.1 | 2.1 | 5 | 24.8 | 3.6 | 5 | 18.3 | 5.0 | 5 |
| **Q9VFR2** | 804 | uncharacterized protein CG9297 (FI18807p1) | | | | | | | | | | 47.9 | 8.9 | 4 | 40.4 | 11.6 | 5 | | | |
| **Q9VG32** | 257 | Malic enzyme | | | | | | | 11.9 | 3.6 | 5 | 8.8 | 1.0 | 4 | 13.1 | 3.2 | 5 | 11.8 | 4.2 | 5 |
| **Q9VG32** | 616 | Malic enzyme | | | | | | | 15.6 | 0.8 | 3 | | | | 21.1 | 5.0 | 5 | 24.4 | 5.5 | 3 |
| **Q9VG32** | 629 | Malic enzyme | 21.3 | 5.7 | 4 | | | | 20.6 | 7.6 | 4 | | | | | | | 30.5 | 5.0 | 3 |
| **Q9VG82** | 490 | Probable cytochrome P450 9f2 | | | | | | | | | | | | | 20.3 | 7.7 | 3 | | | |
| **Q9VGS2** | 172 | Translationally-controlled tumor protein homolog | | | | | | | | | | 4.5 | 0.8 | 3 | 9.7 | 1.5 | 3 | | | |
| **Q9VHJ8** | 369 | Succinyl-coenzyme A synthetase β subunit, ADP-forming (SkpA associated protein; isoform B) | | | | 6.9 | 0.4 | 3 | 10.1 | 4.4 | 5 | 6.8 | 0.4 | 4 | 10.7 | 3.3 | 5 | 12.1 | 5.9 | 5 |
| **Q9VHX4** | 307 | uncharacterized protein CG2767 (LD24679p) | | | | | | | | | | | | | 14.5 | 1.7 | 3 | | | |
| **Q9VIE8** | 133 | Mitochondrial aconitase 1 | 8.3 | 1.8 | 5 | 8.2 | 1.9 | 5 | 7.8 | 3.2 | 5 | 9.9 | 0.8 | 5 | 14.4 | 1.7 | 5 | 12.9 | 3.8 | 5 |
| **Q9VIE8** | 165 | Mitochondrial aconitase 1 | | | | 5.4 | 1.7 | 5 | 5.9 | 5.7 | 4 | 4.9 | 1.0 | 5 | 11.3 | 2.8 | 5 | 8.1 | 3.4 | 5 |
| **Q9VIE8** | 340 | Mitochondrial aconitase 1 | 6.5 | 1.5 | 5 | 6.3 | 1.4 | 5 | 7.2 | 2.8 | 5 | 7.2 | 0.5 | 5 | 14.0 | 2.8 | 5 | 12.8 | 3.9 | 5 |
| **Q9VIE8** | 393 | Mitochondrial aconitase 1 | 19.0 | 2.1 | 5 | 15.7 | 4.4 | 5 | 17.6 | 5.7 | 5 | 21.8 | 1.3 | 5 | 23.8 | 3.1 | 5 | 24.5 | 5.9 | 5 |
| **Q9VIE8** | 404 | Mitochondrial aconitase 1 | 17.5 | 2.5 | 5 | 16.4 | 4.9 | 5 | 16.4 | 7.0 | 5 | 19.4 | 1.7 | 5 | 24.4 | 3.7 | 5 | 22.3 | 6.4 | 5 |
| **Q9VIE8** | 496 | Mitochondrial aconitase 1 | 11.9 | 1.3 | 4 | 7.0 | 1.4 | 3 | 10.0 | 3.6 | 5 | 13.6 | 0.9 | 5 | 16.5 | 2.3 | 5 | 16.1 | 5.0 | 5 |
| **Q9VIE8** | 600 | Mitochondrial aconitase 1 | 7.8 | 1.6 | 5 | 7.2 | 2.3 | 5 | 9.0 | 4.1 | 5 | 9.2 | 1.2 | 5 | 13.0 | 2.1 | 5 | 20.2 | 10.7 | 3 |

| | | | | | | | | | | | | | | | | | | | | |
|---|---|---|---|---|---|---|---|---|---|---|---|---|---|---|---|---|---|---|---|---|
| Q9VIQ8 | 51 | Cytochrome c oxidase subunit 4 (MIP33903p1) | 8.8 | 1.8 | 5 | 7.6 | 1.5 | 5 | 9.1 | 3.7 | 5 | 9.6 | 0.4 | 5 | 14.1 | 2.4 | 5 | | | |
| Q9VJZ4 | 64 | NADH dehydrogenase (ubiquinone) B22 subunit (CG9306) | 12.1 | 2.6 | 5 | 11.1 | 3.2 | 5 | 12.7 | 6.5 | 5 | 13.5 | 0.7 | 5 | 21.6 | 4.2 | 5 | 21.0 | 6.9 | 5 |
| Q9VJZ4 | 87 | NADH dehydrogenase (ubiquinone) B22 subunit (CG9306) | 20.5 | 2.4 | 5 | 20.8 | 3.3 | 5 | 19.9 | 6.0 | 5 | 21.0 | 0.5 | 5 | 31.2 | 4.2 | 5 | 29.3 | 5.7 | 5 |
| Q9VK60 | 84 | uncharacterized protein CG6180 (GH25425p) | | | | | | | | | | | | | 21.9 | 5.6 | 4 | | | |
| Q9VK99 | 51 | atilla (CG6579) | 83.0 | 3.1 | 3 | | | | | | | | | | | | | | | |
| Q9VK99 | 69 | atilla (CG6579) | | | | | | | | | | | | | | | | 95.5 | 2.0 | 4 |
| Q9VL70 | 79 | Yippee interacting protein 2 | | | | 15.0 | 3.8 | 4 | 11.3 | 2.6 | 4 | 15.5 | 3.9 | 5 | 16.9 | 3.0 | 5 | 14.2 | 4.1 | 5 |
| Q9VL70 | 94 | Yippee interacting protein 2 | | | | | | | | | | 16.8 | 1.4 | 3 | 21.0 | 5.3 | 5 | | | |
| Q9VL70 | 256 | Yippee interacting protein 2 | | | | | | | | | | | | | 16.7 | 2.0 | 4 | | | |
| Q9VLB7 | 282 | GDP dissociation inhibitor (LP03430p) | 20.9 | 2.5 | 4 | 21.9 | 5.9 | 3 | 23.8 | 5.6 | 5 | 25.6 | 0.6 | 5 | 28.9 | 8.2 | 4 | 24.2 | 8.0 | 5 |
| Q9VLP2 | 63 | uncharacterized protein CG7781 (RE71014p) | | | | | | | | | | 88.4 | 1.5 | 5 | | | | | | |
| Q9VLZ3 | 1792 | Myosin 28B1; isoform A | 8.9 | 2.2 | 3 | | | | | | | 10.3 | 2.2 | 4 | | | | | | |
| Q9VMI3 | 140 | NADH dehydrogenase (ubiquinone) 51 kDa subunit (GM14163p) | 10.6 | 1.9 | 5 | 10.2 | 2.1 | 5 | 9.1 | 2.9 | 5 | 10.9 | 1.2 | 5 | 17.9 | 2.1 | 5 | 16.0 | 4.2 | 5 |
| Q9VMI3 | 157 | NADH dehydrogenase (ubiquinone) 51 kDa subunit (GM14163p) | | | | | | | | | | 11.4 | 1.4 | 4 | 13.4 | 3.3 | 3 | | | |
| Q9VMI3 | 202 | NADH dehydrogenase (ubiquinone) 51 kDa subunit (GM14163p) | 14.5 | 2.2 | 3 | 12.7 | 3.1 | 4 | | | | 14.1 | 1.4 | 4 | 20.8 | 3.5 | 5 | 19.9 | 6.3 | 4 |
| Q9VMI3 | 221 | NADH dehydrogenase (ubiquinone) 51 kDa subunit (GM14163p) | 11.0 | 1.9 | 5 | 9.9 | 2.0 | 5 | 11.0 | 4.6 | 5 | 11.8 | 0.8 | 5 | 17.2 | 2.6 | 5 | 18.2 | 6.4 | 5 |
| Q9VMI3 | 301 | NADH dehydrogenase (ubiquinone) 51 kDa subunit (GM14163p) | | | | | | | | | | 12.2 | 1.5 | 4 | 23.5 | 4.8 | 4 | | | |
| Q9VMI3 | 319 | NADH dehydrogenase (ubiquinone) 51 kDa subunit (GM14163p) | 9.5 | 1.1 | 5 | 11.3 | 0.3 | 3 | 10.4 | 4.0 | 3 | 12.0 | 4.1 | 5 | 19.7 | 3.2 | 5 | | | |
| Q9VMQ9 | 82 | clot (RE06767p) | | | | 53.0 | 4.3 | 4 | 54.1 | 4.9 | 3 | | | | 63.9 | 10.5 | 4 | 56.1 | 9.4 | 3 |
| Q9VMS1 | 77 | Cytochrome c oxidase subunit VIc | 7.0 | 2.9 | 5 | 7.8 | 2.7 | 5 | 7.0 | 4.6 | 5 | 35.1 | 24.4 | 3 | | | | | | |
| Q9VMT2 | 67 | Troponin C at 25D | | | | | | | | | | 11.3 | 1.4 | 4 | 22.9 | 11.6 | 4 | 26.2 | 5.9 | 3 |
| Q9VMU0 | 80 | NADH dehydrogenase (ubiquinone) 13 kDa A subunit (LP20380p) | 23.5 | 2.0 | 5 | 21.1 | 4.2 | 5 | 20.0 | 5.5 | 5 | 23.0 | 1.6 | 5 | 31.7 | 5.3 | 5 | 29.4 | 6.7 | 5 |
| Q9VMU0 | 87 | NADH dehydrogenase (ubiquinone) 13 kDa A subunit (LP20380p) | 23.9 | 2.0 | 5 | 23.5 | 4.2 | 5 | 23.2 | 5.0 | 5 | 26.1 | 2.0 | 5 | 33.3 | 3.4 | 5 | 30.9 | 6.1 | 5 |
| Q9VN13 | 61 | Sideroflexin 1/3 (RH48017p) | | | | 7.5 | 1.6 | 4 | 9.4 | 4.3 | 4 | | | | | | | | | |
| Q9VNB9 | 103 | Ribosomal protein L35A | | | | | | | | | | 9.6 | 1.9 | 4 | 14.8 | 3.9 | 5 | 12.4 | 5.8 | 4 |
| Q9VNE9 | 40 | 60S ribosomal protein L13a | | | | | | | | | | 6.2 | 0.2 | 3 | 10.0 | 2.2 | 3 | | | |
| Q9VNE9 | 138 | 60S ribosomal protein L13a | | | | | | | 10.6 | 4.2 | 4 | 10.0 | 1.2 | 5 | 23.0 | 5.9 | 5 | 17.4 | 2.7 | 5 |
| Q9VNW6 | 91 | Delta-1-pyrroline-5-carboxylate synthase (GH12632p) | | | | | | | | | | | | | 6.1 | 2.0 | 4 | | | |
| Q9VNW6 | 570 | Delta-1-pyrroline-5-carboxylate synthase (GH12632p) | | | | | | | | | | | | | 16.3 | 3.3 | 3 | 18.8 | 2.3 | 4 |
| Q9VNW6 | 704 | Delta-1-pyrroline-5-carboxylate synthase (GH12632p) | | | | 11.2 | 4.3 | 3 | 10.8 | 4.2 | 5 | | | | 14.5 | 2.1 | 5 | 16.8 | 4.5 | 5 |
| Q9VNX4 | 70 | Delta-1-Pyrroline-5-carboxylate dehydrogenase 1 (FI08816p) | | | | | | | | | | | | | 13.3 | 2.1 | 3 | | | |
| Q9VNX4 | 355 | Delta-1-Pyrroline-5-carboxylate dehydrogenase 1 (FI08816p) | 23.5 | 1.4 | 4 | 20.5 | 6.2 | 4 | | | | | | | 29.5 | 6.3 | 4 | 31.2 | 4.2 | 4 |
| Q9VNX4 | 381 | Delta-1-Pyrroline-5-carboxylate dehydrogenase 1 (FI08816p) | | | | | | | | | | | | | 16.8 | 3.3 | 4 | | | |
| Q9VNX4 | 502 | Delta-1-Pyrroline-5-carboxylate dehydrogenase 1 (FI08816p) | 6.4 | 2.4 | 4 | 5.3 | 1.5 | 5 | 8.2 | 4.2 | 4 | 5.9 | 0.4 | 5 | 10.6 | 2.1 | 5 | 11.2 | 5.3 | 5 |
| Q9VP61 | 227 | Acetyl-coenzyme A synthetase | | | | | | | | | | 12.6 | 0.7 | 4 | 18.2 | 3.7 | 3 | | | |
| Q9VPE2 | 334 | NADH dehydrogenase (ubiquinone) 39 kDa subunit (CG6020) | 14.8 | 3.6 | 5 | 15.2 | 2.9 | 5 | 12.6 | 4.5 | 5 | 15.0 | 1.4 | 5 | 21.8 | 2.6 | 5 | 19.0 | 4.6 | 5 |
| Q9VPE2 | 405 | NADH dehydrogenase (ubiquinone) 39 kDa subunit (CG6020) | 12.1 | 2.1 | 5 | | | | | | | | | | | | | 20.7 | 3.0 | 3 |
| Q9VQ61 | 181 | Aspartate aminotransferase | | | | | | | | | | | | | 18.5 | 4.4 | 4 | 14.7 | 5.2 | 3 |
| Q9VQ61 | 266 | Aspartate aminotransferase | 12.3 | 3.7 | 5 | 10.1 | 1.8 | 5 | 13.1 | 6.1 | 4 | | | | 17.2 | 3.2 | 5 | 20.2 | 7.0 | 5 |
| Q9VQG4 | 63 | Congested-like trachea protein | | | | | | | | | | | | | 16.3 | 3.5 | 3 | | | |
| Q9VQL7 | 351 | Fatty acid synthase 1 (CG3523; isoform A) | | | | | | | | | | | | | 17.0 | 3.8 | 3 | | | |
| Q9VQL7 | 377 | Fatty acid synthase 1 (CG3523; isoform A) | 16.7 | 1.4 | 3 | 11.4 | 2.2 | 5 | 10.8 | 4.8 | 5 | 11.9 | 0.8 | 5 | 19.3 | 5.8 | 5 | 16.5 | 4.7 | 5 |
| Q9VQL7 | 561 | Fatty acid synthase 1 (CG3523; isoform A) | | | | | | | | | | | | | | | | 4.6 | 0.8 | 3 |
| Q9VQL7 | 1491 | Fatty acid synthase 1 (CG3523; isoform A) | | | | | | | | | | 7.9 | 2.1 | 4 | 11.4 | 8.0 | 3 | | | |
| Q9VQL7 | 1553 | Fatty acid synthase 1 (CG3523; isoform A) | 8.3 | 2.0 | 5 | 7.5 | 1.9 | 5 | 7.4 | 3.6 | 5 | 8.1 | 0.6 | 4 | 12.3 | 1.9 | 3 | 13.0 | 1.7 | 3 |
| Q9VQL7 | 2403 | Fatty acid synthase 1 (CG3523; isoform A) | 8.2 | 1.9 | 4 | 7.8 | 2.9 | 3 | 7.9 | 1.3 | 3 | 8.3 | 1.2 | 5 | 13.0 | 5.3 | 5 | 11.9 | 4.7 | 5 |
| Q9VQR2 | 90 | NADH dehydrogenase (ubiquinone) PDSW subunit (RE65754p) | 69.9 | 9.0 | 5 | 75.5 | 3.9 | 5 | 75.1 | 12.8 | 5 | 64.1 | 16.7 | 3 | 89.0 | 3.3 | 4 | 81.0 | 9.6 | 4 |
| Q9VQR2 | 102 | NADH dehydrogenase (ubiquinone) PDSW subunit (RE65754p) | | | | | | | | | | 70.4 | 15.1 | 4 | 87.5 | 2.2 | 4 | 77.7 | 6.0 | 4 |
| Q9VQT8 | 63 | Immune induced molecule 33 (RH38008p) | 94.7 | 1.7 | 5 | 93.7 | 1.1 | 4 | 95.4 | 1.2 | 5 | 94.4 | 2.2 | 5 | 97.5 | 0.9 | 4 | 95.4 | 1.7 | 4 |
| Q9VQT8 | 76 | Immune induced molecule 33 (RH38008p) | | | | 91.6 | 0.7 | 3 | | | | | | | | | | | | |
| Q9VS34 | 101 | 60S ribosomal protein L18 | 5.6 | 1.9 | 4 | 7.7 | 2.7 | 3 | 6.8 | 4.9 | 4 | 5.7 | 0.6 | 4 | 13.0 | 4.4 | 5 | 11.5 | 3.1 | 4 |
| Q9VSA3 | 355 | Medium-chain specific acyl-CoA dehydrogenase; mitochondrial | | | | | | | | | | 8.9 | 1.6 | 4 | 12.5 | 2.4 | 3 | | | |
| Q9VSA9 | 130 | uncharacterized protein CG7409 (RH03891p) | 9.1 | 2.3 | 5 | 7.5 | 4.7 | 3 | 8.9 | 4.8 | 5 | | | | 18.2 | 4.4 | 3 | 20.0 | 3.2 | 4 |
| Q9VSU6 | 19 | Dihydropteridine reductase (RE58329p) | | | | | | | | | | | | | 15.4 | 1.9 | 4 | | | |
| Q9VTB4 | 74 | NADH dehydrogenase (ubiquinone) 13 kDa B subunit (RH11203p) | 11.9 | 2.2 | 5 | 10.8 | 2.6 | 5 | 11.8 | 2.9 | 4 | 12.8 | 1.9 | 5 | 20.8 | 4.5 | 5 | 21.8 | 5.7 | 5 |
| Q9VTK9 | 46 | Aldo-keto reductase 1B (LD06393p) | | | | | | | | | | | | | 13.3 | 4.5 | 4 | 17.1 | 8.6 | 5 |
| Q9VTK9 | 186 | Aldo-keto reductase 1B (LD06393p) | | | | | | | | | | 8.1 | 0.8 | 3 | 13.7 | 4.0 | 4 | | | |
| Q9VTY2 | 202 | uncharacterized protein CG10638; isoform A | | | | | | | 11.4 | 5.6 | 4 | | | | | | | 17.0 | 1.3 | 3 |
| Q9VU39 | 135 | Adenosine kinase (GH14845p) | | | | | | | | | | | | | 19.1 | 3.4 | 3 | | | |
| Q9VU39 | 318 | Adenosine kinase (GH14845p) | | | | | | | 29.2 | 8.2 | 5 | 33.3 | 5.5 | 5 | 33.5 | 7.7 | 5 | 35.6 | 10.7 | 5 |
| Q9VU68 | 507 | flare (Actin-interacting protein 1) | | | | | | | | | | | | | | | | 21.3 | 14.1 | 3 |
| Q9VUY9 | 132 | Phosphoglucomutase | | | | | | | | | | 9.1 | 0.8 | 3 | 13.9 | 2.6 | 4 | | | |
| Q9VV75 | 304 | Ubiquinol-cytochrome c reductase core protein 2 (CG4169) | 9.8 | 1.9 | 5 | 7.6 | 2.2 | 5 | 8.6 | 2.9 | 5 | 10.0 | 1.1 | 5 | 15.0 | 2.5 | 5 | 15.8 | 5.6 | 5 |
| Q9VVH3 | 90 | MICOS complex subunit MIC13 homolog QIL1 (RE63168p) | | | | | | | | | | 15.9 | 1.3 | 5 | 23.3 | 5.2 | 5 | | | |
| Q9VVL7 | 109 | Dihydrolipoyl dehydrogenase | 11.9 | 2.5 | 5 | 11.6 | 3.9 | 4 | 12.1 | 4.5 | 5 | 14.4 | 0.6 | 5 | 19.6 | 4.0 | 5 | 19.6 | 5.9 | 5 |
| Q9VVL7 | 301 | Dihydrolipoyl dehydrogenase | 9.4 | 3.4 | 5 | 7.0 | 1.1 | 5 | 10.0 | 3.5 | 5 | 8.1 | 0.4 | 5 | 11.8 | 2.2 | 5 | 16.2 | 6.4 | 5 |
| Q9VVL7 | 351 | Dihydrolipoyl dehydrogenase | | | | | | | | | | 11.9 | 0.2 | 3 | 14.8 | 1.6 | 3 | 17.5 | 6.8 | 3 |
| Q9VVU1 | 241 | CG3902 (FI09602p) | | | | | | | | | | | | | 15.3 | 1.2 | 3 | | | |
| Q9VVU1 | 407 | CG3902 (FI09602p) | | | | 7.5 | 3.4 | 5 | 7.4 | 4.7 | 4 | | | | | | | | | |
| Q9VW68 | 303 | Gamma-aminobutyric acid transaminase (CG7433; isoform C) | 6.7 | 1.9 | 5 | 5.7 | 2.4 | 4 | 8.1 | 5.2 | 5 | 7.7 | 1.4 | 4 | 13.3 | 2.6 | 4 | 14.4 | 6.3 | 5 |
| Q9VW68 | 454 | Gamma-aminobutyric acid transaminase (CG7433; isoform C) | | | | | | | | | | 13.6 | 1.8 | 4 | 16.2 | 3.0 | 3 | | | |
| Q9VWH4 | 143 | Probable isocitrate dehydrogenase [NAD] subunit alpha, mitochondria | 10.9 | 1.8 | 5 | 9.6 | 2.5 | 5 | 9.2 | 3.0 | 5 | 11.5 | 1.3 | 5 | 15.4 | 3.0 | 5 | 15.8 | 4.0 | 5 |
| Q9VWH4 | 252 | Probable isocitrate dehydrogenase [NAD] subunit alpha, mitochondria | | | | | | | | | | 9.7 | 0.4 | 3 | | | | | | |
| Q9VWV6 | 62 | Transferrin 1 | | | | | | | 91.7 | 3.2 | 3 | | | | | | | 94.6 | 3.6 | 3 |
| Q9VWV6 | 71 | Transferrin 1 | | | | | | | 86.9 | 5.1 | 4 | | | | | | | | | |
| Q9VWV6 | 143 | Transferrin 1 | 79.5 | 5.5 | 4 | 78.3 | 6.2 | 5 | 72.7 | 13.4 | 5 | 70.2 | 3.2 | 5 | 88.1 | 12.3 | 4 | 91.6 | 5.5 | 5 |
| Q9VWV6 | 239 | Transferrin 1 | 83.1 | 1.0 | 3 | 78.8 | 6.5 | 5 | 81.8 | 7.5 | 5 | 78.2 | 3.5 | 4 | 90.8 | 6.8 | 5 | 91.5 | 5.2 | 5 |
| Q9VWV6 | 397 | Transferrin 1 | | | | | | | 86.7 | 6.8 | 5 | | | | | | | 91.3 | 6.7 | 3 |
| Q9VWV6 | 486 | Transferrin 1 | 35.8 | 4.5 | 4 | 30.1 | 7.7 | 3 | 24.9 | 7.1 | 5 | 31.9 | 5.9 | 5 | 35.5 | 3.4 | 5 | 30.4 | 6.3 | 5 |
| Q9VWV6 | 533 | Transferrin 1 | | | | | | | | | | | | | | | | 88.3 | 6.0 | 3 |
| Q9VWV6 | 547 | Transferrin 1 | | | | | | | 69.5 | 9.6 | 5 | | | | | | | 83.6 | 9.3 | 4 |
| Q9VWV6 | 633 | Transferrin 1 | | | | | | | 69.4 | 11.3 | 5 | | | | | | | | | |
| Q9VX36 | 145 | NADH dehydrogenase (ubiquinone) 24 kDa subunit (CG5703) | 14.4 | 2.1 | 5 | 13.0 | 4.0 | 5 | 17.8 | 7.7 | 4 | 15.9 | 2.1 | 5 | 22.9 | 5.3 | 5 | 24.4 | 8.0 | 5 |
| Q9VXI6 | 36 | Cytochrome b-c1 complex subunit 7 (RH44664p) | 9.5 | 3.3 | 5 | 10.8 | 3.2 | 5 | 8.6 | 3.8 | 5 | 10.5 | 0.9 | 5 | 23.2 | 3.3 | 5 | 18.7 | 6.2 | 5 |
| Q9VY93 | 22 | Probable methylthioribulose-1-phosphate dehydratase | | | | | | | | | | 10.2 | 0.7 | 4 | 13.6 | 1.3 | 4 | 11.8 | 2.7 | 5 |
| Q9VYR1 | 76 | Regucalcin (SD03837p) | | | | | | | | | | 12.7 | 0.2 | 3 | 12.0 | 1.0 | 3 | | | |
| Q9VYU9 | 230 | uncharacterized protein CG9360 (RH17287p) | | | | | | | | | | | | | 16.5 | 3.0 | 4 | 24.0 | 2.0 | 4 |
| Q9VYU9 | 22 23 | uncharacterized protein CG9360 (RH17287p) | | | | | | | | | | | | | | | | | | |
| Q9VZ01 | 102 | NADH:ubiquinone oxidoreductase subunit V3 (RH44935p) | 23.5 | 3.8 | 5 | 21.5 | 5.5 | 5 | 22.3 | 4.2 | 5 | 22.7 | 3.0 | 5 | 34.3 | 5.1 | 5 | 34.8 | 5.9 | 5 |
| Q9VZF6 | 183 | Sulfide quinone oxidoreductase (GH04863p) | | | | | | | 27.8 | 5.0 | 3 | | | | | | | | | |
| Q9VZR2 | 58 | Drosomycin-like 5 | | | | 95.7 | 0.2 | 4 | 97.1 | 1.2 | 5 | 96.0 | 4.2 | 5 | 98.3 | 0.7 | 5 | 96.9 | 1.9 | 4 |
| Q9VZU4 | 70 | NADH dehydrogenase (ubiquinone) 30 kDa subunit (RH59487p) | | | | | | | | | | 2.5 | 0.1 | 3 | 7.4 | 2.9 | 4 | 6.3 | 4.4 | 3 |
| Q9VZU4 | 159 | NADH dehydrogenase (ubiquinone) 30 kDa subunit (RH59487p) | 10.4 | 2.1 | 5 | 11.0 | 2.1 | 5 | 11.0 | 4.9 | 5 | 11.0 | 0.5 | 5 | 17.9 | 3.0 | 5 | 19.9 | 5.9 | 5 |
| Q9VZU4 | 223 | NADH dehydrogenase (ubiquinone) 30 kDa subunit (RH59487p) | 8.2 | 2.0 | 5 | 7.0 | 2.0 | 5 | 8.7 | 4.3 | 5 | 9.3 | 1.7 | 5 | 14.9 | 3.7 | 5 | 15.8 | 5.6 | 5 |
| Q9W0P5 | 154 | UDP-glucose 4-epimerase | | | | | | | | | | 15.5 | 2.3 | 5 | 19.8 | 3.6 | 5 | 22.1 | 3.0 | 3 |
| Q9W0P5 | 310 | UDP-glucose 4-epimerase | | | | | | | | | | | | | 18.8 | 2.7 | 5 | | | |
| Q9W125 | 68 | NADH dehydrogenase (ubiquinone) 19 kDa subunit (RE35078p) | | | | | | | | | | 78.1 | 17.7 | 4 | 90.9 | 1.3 | 5 | 81.2 | 7.7 | 3 |
| Q9W125 | 103 | NADH dehydrogenase (ubiquinone) 19 kDa subunit (RE35078p) | | | | | | | | | | 82.2 | 6.8 | 5 | | | | | | |
| Q9W1C9 | 46 | Ejaculatory bulb-specific protein 3 | 89.4 | 4.5 | 4 | 89.3 | 1.6 | 3 | | | | 92.0 | 0.8 | 3 | 94.9 | 1.6 | 3 | | | |
| Q9W1C9 | 72 | Ejaculatory bulb-specific protein 3 | 82.4 | 3.0 | 5 | 82.8 | 5.0 | 5 | 82.9 | 2.2 | 5 | 81.9 | 1.4 | 5 | 90.0 | 2.6 | 5 | 84.9 | 6.0 | 5 |
| Q9W303 | 426 | Imaginal disc growth factor 4 (Chitinase-like protein Idgf4) | | | | | | | | | | | | | 81.2 | 11.7 | 3 | 77.5 | 7.5 | 4 |
| Q9W334 | 23 | Ribosomal protein S28b (40S ribosomal protein S28) | | | | | | | | | | | | | 23.7 | 6.9 | 4 | 26.0 | 9.3 | 5 |
| Q9W369 | 380 | tan | | | | | | | | | | 15.3 | 1.7 | 3 | 25.8 | 7.8 | 5 | | | |
| Q9W3B3 | 141 | Uroporphyrinogen-III synthase 1 (GH17465p) | | | | | | | | | | 24.0 | 3.2 | 4 | 31.9 | 6.2 | 4 | | | |
| Q9W3B3 | 239 | Uroporphyrinogen-III synthase 1 (GH17465p) | | | | 10.7 | 3.0 | 4 | | | | 12.3 | 1.7 | 5 | 17.2 | 2.1 | 5 | 20.6 | 2.1 | 3 |
| Q9W3L4 | 25 | uncharacterized protein CG2233 (GH20802p) | | | | | | | 61.1 | 12.0 | 4 | | | | | | | | | |
| Q9W3R8 | 123 | uncharacterized protein CG2059 | | | | | | | 13.0 | 4.7 | 5 | | | | 21.2 | 3.6 | 4 | | | |
| Q9W401 | 102 | Citrate synthase 1 (Probable citrate synthase, mitochondrial) | 8.5 | 1.7 | 5 | 9.1 | 2.1 | 5 | 8.0 | 3.2 | 5 | 11.3 | 1.7 | 5 | 16.9 | 1.6 | 5 | 14.3 | 3.0 | 5 |
| Q9W401 | 219 | Citrate synthase 1 (Probable citrate synthase, mitochondrial) | | | | 5.5 | 0.3 | 3 | 9.0 | 4.3 | 4 | 5.0 | 0.8 | 4 | 7.8 | 2.2 | 4 | 9.6 | 5.6 | 4 |
| Q9W401 | 360 | Citrate synthase 1 (Probable citrate synthase, mitochondrial) | 10.4 | 1.8 | 5 | 10.0 | 2.6 | 5 | 8.8 | 3.7 | 5 | | | | | | | 14.8 | 4.7 | 3 |
| Q9W4P5 | 243 | V-type proton ATPase subunit d 1 | 10.4 | 1.7 | 5 | 8.6 | 2.7 | 3 | 8.9 | 3.9 | 4 | | | | 15.1 | 2.8 | 3 | | | |
| Q9W4Y3 | 3404 | Terribly reduced optic lobes; isoform F | | | | | | | | | | 92.7 | 2.2 | 4 | | | | | | |
| Q9W4Y3 | 3858 | Terribly reduced optic lobes; isoform F | 78.6 | 2.5 | 3 | 72.5 | 8.9 | 3 | | | | 71.4 | 7.3 | 3 | | | | | | |
| Q9W4Y3 | 4330 | Terribly reduced optic lobes; isoform F | | | | | | | | | | 77.0 | 10.0 | 4 | | | | | | |
| Q9XTL9 | 301 | Glycogen phosphorylase | | | | | | | | | | 8.1 | 4.2 | 4 | 11.0 | 3.9 | 4 | 10.0 | 4.1 | 4 |
| Q9XTL9 | 373 | Glycogen phosphorylase | | | | 7.4 | 0.0 | 3 | | | | 7.4 | 0.2 | 4 | 10.5 | 2.1 | 3 | 12.4 | 1.5 | 3 |
| Q9XTL9 | 496 | Glycogen phosphorylase | | | | | | | | | | 3.7 | 0.3 | 4 | 6.6 | 1.5 | 5 | | | |
| Q9XYN7 | 19 | Proteasome subunit beta type-3 | | | | | | | | | | 16.3 | 3.2 | 4 | 18.2 | 8.0 | 4 | 23.3 | 10.2 | 4 |
| Q9Y112 | 96 | uncharacterized protein CG10863 | 12.2 | 3.5 | 3 | 11.3 | 4.0 | 5 | 12.0 | 6.3 | 5 | 12.1 | 0.8 | 5 | 18.9 | 2.7 | 3 | 18.0 | 5.9 | 5 |
| Q9Y112 | 214 | uncharacterized protein CG10863 | 25.9 | 3.2 | 5 | | | | 23.6 | 4.2 | 5 | 23.6 | 0.8 | 5 | 36.6 | 4.4 | 5 | 33.9 | 5.7 | 5 |

**Supplementary Table 3 | Molecular function and pathway analysis of proteins identified in this study by OxICAT redox proteomics**
Lennicke *et al.*

| Uniprot ID | FlyBase Gene ID | Annotation Symbol | Gene Symbol | Name | GOTERM_MF_DIRECT (Molecular Function, Direct) | KEGG_PATHWAY |
|---|---|---|---|---|---|---|
| **A1Z7H3** | FBgn0263120 | CG8732 | Acsl | Acyl-CoA synthetase long-chain<br>(Lethal (2) 44DEa; isoform I) | GO:0004467~long-chain fatty acid-CoA ligase activity,<br>GO:0090433~palmitoyl-CoA ligase activity, | dme00061:Fatty acid biosynthesis,<br>dme00071:Fatty acid degradation,<br>dme01100:Metabolic pathways,<br>dme01212:Fatty acid metabolism,<br>dme04146:Peroxisome, |
| **A1Z8U9** | FBgn0033699 | CG8857 | RpS11 | Ribosomal protein S11; isoform B | GO:0003735~structural constituent of ribosome,<br>GO:0019843~rRNA binding, | dme03010:Ribosome, |
| **A1Z992** | FBgn0053138 | CG33138 | AGBE | 1,4-Alpha-glucan branching enzyme | GO:0003824~catalytic activity,<br>GO:0003844~1,4-alpha-glucan branching enzyme activity,<br>GO:0004553~hydrolase activity, hydrolyzing O-glycosyl compounds,<br>GO:0016798~hydrolase activity, acting on glycosyl bonds,<br>GO:0043169~cation binding, | dme00500:Starch and sucrose metabolism,<br>dme01100:Metabolic pathways, |
| **A1Z9E3** | FBgn0024556 | CG6050 | mEFTu1 | Elongation factor Tu<br>(Mitochondrial translation elongation factor Tu 1) | GO:0003746~translation elongation factor activity,<br>GO:0003924~GTPase activity,<br>GO:0005525~GTP binding, | |
| **A1Z9G2** | FBgn0033859 | CG6197 | fand | fandango<br>(Pre-mRNA-splicing factor syf1 homolog)<br>(Fl18620p1) | | dme03040:Spliceosome, |
| **A1ZA47** | FBgn0265991 | CG30084 | Zasp52 | PDZ and LIM domain protein Zasp<br>(Z band alternatively spliced PDZ-motif protein 52) | GO:0003779~actin binding,<br>GO:0042805~actinin binding,<br>GO:0046872~metal ion binding,<br>GO:0051371~muscle alpha-actinin binding, | |
| **A1ZA73** | FBgn0265045 | CG44162 | Strn-Mlck | Stretchin-Mlck; isoform E | GO:0004672~protein kinase activity<br>GO:0004674~protein serine/threonine kinase activity,<br>GO:0004683~calmodulin-dependent protein kinase activity,<br>GO:0004687~myosin light chain kinase activity,<br>GO:0005200~structural constituent of cytoskeleton,<br>GO:0005524~ATP binding, | |
| **A1ZBJ2** | FBgn0034432 | CG7461 | Acadvl | Acyl-CoA dehydrogenase very long chain<br>(CG7461) | GO:0000062~fatty-acyl-CoA binding,<br>GO:0003995~acyl-CoA dehydrogenase activity,<br>GO:0004466~long-chain-acyl-CoA dehydrogenase activity,<br>GO:0016627~oxidoreductase activity, acting on the CH-CH group of donors,<br>GO:0017099~very-long-chain-acyl-CoA dehydrogenase activity,<br>GO:0050660~flavin adenine dinucleotide binding | dme00071:Fatty acid degradation,<br>dme01100:Metabolic pathways,<br>dme01212:Fatty acid metabolism, |
| **A4UZR3** | FBgn0003748 | CG9364 | Treh | Trehalase; isoform E | GO:0004555~alpha,alpha-trehalase activity, | dme00500:Starch and sucrose metabolism,<br>dme01100:Metabolic pathways, |
| **A4V4Q6** | FBgn0283471 | CG7178 | wupA | wings up A; isoform F | GO:0003779~actin binding, | |
| **A5XCL5** | FBgn0035978 | CG4347 | UGP | UDP-glucose pyrophosphorylase; isoform D | GO:0003983~UTP:glucose-1-phosphate uridylyltransferase activity,<br>GO:0070569~uridylyltransferase activity, | dme00040:Pentose and glucuronate interconversions,<br>dme00052:Galactose metabolism,<br>dme00500:Starch and sucrose metabolism,<br>dme00520:Amino sugar and nucleotide sugar metabolism,<br>dme01100:Metabolic pathways,<br>dme01240:Biosynthesis of cofactors,<br>dme01250:Biosynthesis of nucleotide sugars, |
| **A8JRC2 (obsolete)** | FBgn0039358 | CG5028 | Idh3g | Isocitrate dehydrogenase (NAD(+)) 3 non-catalytic subunit gamma<br>(CG5028; isoform E)<br>(https://www.uniprot.org/uniparc?query=A8JRC2) | GO:0004449~isocitrate dehydrogenase (NAD+) activity, | dme00020:Citrate cycle (TCA cycle),<br>dme01100:Metabolic pathways,<br>dme01200:Carbon metabolism,<br>dme01210:2-Oxocarboxylic acid metabolism,<br>dme01230:Biosynthesis of amino acids, |
| **A8Y535** | FBgn0260008 | CG41623 | UQCR-11 | Ubiquinol-cytochrome c reductase complex 11 kDa subunit; isoform D<br>(Cytochrome b-c1 complex subunit 6) | | dme00190:Oxidative phosphorylation,<br>dme01100:Metabolic pathways, |
| **B7YZP9** | FBgn0259209 | CG42309 | Mlp60A | Muscle LIM protein at 60A; isoform B | GO:0046872~metal ion binding, | |
| **B7YZQ7** | FBgn0016687 | CG4634 | Nurf-38 | Nucleosome remodeling factor - 38kD; isoform B | GO:0000287~magnesium ion binding,<br>GO:0004427~inorganic diphosphatase activity, | dme00190:Oxidative phosphorylation,<br>dme03082:ATP-dependent chromatin remodeling, |
| **B7Z061** | FBgn0011693 | CG4899 | Pdh | Photoreceptor dehydrogenase; isoform D | GO:0000166~nucleotide binding,<br>GO:0004745~retinol dehydrogenase activity,<br>GO:0016491~oxidoreductase activity,<br>GO:0016616~oxidoreductase activity, acting on the CH-OH group of donors, NAD or NADP as acceptor, | |
| **E1JH64** | FBgn0013733 | CG18076 | shot | short stop; isoform Q | GO:0003779~actin binding,<br>GO:0005198~structural molecule activity,<br>GO:0005509~calcium ion binding,<br>GO:0005515~protein binding,<br>GO:0008017~microtubule binding,<br>GO:0008093~cytoskeletal adaptor activity, | |
| **E1JHJ5** | FBgn0264695 | CG17927 | Mhc | Myosin heavy chain; isoform P | GO:0000146~microfilament motor activity,<br>GO:0003774~motor activity,<br>GO:0005516~calmodulin binding,<br>GO:0005524~ATP binding,<br>GO:0008307~structural constituent of muscle,<br>GO:0016853~isomerase activity<br>GO:0042803~protein homodimerization activity,<br>GO:0051015~actin filament binding, | dme04814:Motor proteins, |
| **E1JHR5** | FBgn0000579 | CG17654 | Eno | Enolase; isoform F | GO:0000287~magnesium ion binding,<br>GO:0004634~phosphopyruvate hydratase activity, | dme00010:Glycolysis / Gluconeogenesis,<br>dme01100:Metabolic pathways,<br>dme01200:Carbon metabolism,<br>dme01230:Biosynthesis of amino acids,<br>dme03018:RNA degradation, |
| **E1JIR4** | FBgn0002921 | CG5670 | Atpalpha | Na pump alpha subunit; isoform I<br>(Sodium/potassium-transporting ATPase subunit alpha) | GO:0000166~nucleotide binding,<br>GO:0005391~sodium:potassium-exchanging ATPase activity,<br>GO:0005524~ATP binding,<br>GO:0008556~potassium-transporting ATPase activity,<br>GO:0016887~ATPase activity,<br>GO:0046872~metal ion binding, | |
| **E2QCF1** | FBgn0020236 | CG8322 | ATPCL<br>(Acly) | ATP citrate lyase; isoform F | GO:0003824~catalytic activity,<br>GO:0003878~ATP citrate synthase activity,<br>GO:0005524~ATP binding,<br>GO:0016829~lyase activity,<br>GO:0046872~metal ion binding,<br>GO:0046912~acyltransferase activity, acyl groups converted into alkyl on transfer | dme00020:Citrate cycle (TCA cycle),<br>dme01100:Metabolic pathways, |
| **O01666** | FBgn0020235 | CG7610 | ATPsyn-gamma | ATP synthase subunit gamma; mitochondrial | GO:0046933~proton-transporting ATP synthase activity, rotational mechanism, | dme00190:Oxidative phosphorylation,<br>dme01100:Metabolic pathways, |
| **O18404** | FBgn0021765 | CG7113 | scu | 3-hydroxyacyl-CoA dehydrogenase type-2)<br>(scully) | GO:0003857~3-hydroxyacyl-CoA dehydrogenase activity,<br>GO:0004303~estradiol 17-beta-dehydrogenase activity,<br>GO:0016229~steroid dehydrogenase activity,<br>GO:0016491~oxidoreductase activity,<br>GO:0018454~acetoacetyl-CoA reductase activity,<br>GO:0035410~dihydrotestosterone 17-beta-dehydrogenase activity,<br>GO:0047015~3-hydroxy-2-methylbutyryl-CoA dehydrogenase activity,<br>GO:0047022~7-beta-hydroxysteroid dehydrogenase (NADP+) activity,<br>GO:0047035~testosterone dehydrogenase (NAD+) activity,<br>GO:0047045~testosterone 17-beta-dehydrogenase (NADP+) activity, | dme00280:Valine, leucine and isoleucine degradation,<br>dme01100:Metabolic pathways, |
| **O46037** | FBgn0004397 | CG3299 | Vinc | Vinculin | GO:0003779~actin binding,<br>GO:0005198~structural molecule activity,<br>GO:0008013~beta-catenin binding,<br>GO:0045294~alpha-catenin binding,<br>GO:0051015~actin filament binding, | |
| **O46231** | FBgn0082582 | CG1539 | tmod | Tropomodulin; isoform A | GO:0005523~tropomyosin binding, | |
| **O61231** | FBgn0024733 | CG17521 | RpL10 | 60S ribosomal protein L10<br>(Ribosomal protein L10) | GO:0003735~structural constituent of ribosome, | dme03010:Ribosome, |
| **O62619** | FBgn0267385 | CG7070 | Pyk | Pyruvate kinase | GO:0000287~magnesium ion binding,<br>GO:0003824~catalytic activity,<br>GO:0004743~pyruvate kinase activity,<br>GO:0005524~ATP binding,<br>GO:0016301~kinase activity,<br>GO:0030955~potassium ion binding, | dme00010:Glycolysis / Gluconeogenesis,<br>dme00620:Pyruvate metabolism,<br>dme01100:Metabolic pathways,<br>dme01200:Carbon metabolism,<br>dme01230:Biosynthesis of amino acids, |
| **O76927** | FBgn0015521 | CG2986 | RpS21 | 40S ribosomal protein S21<br>(Ribosomal protein S21) | GO:0003735~structural constituent of ribosome,<br>GO:0043022~ribosome binding, | dme03010:Ribosome, |
| **O97394** | FBgn0021764 | CG5227 | sdk | Protein sidekick | | |
| **O97418** | FBgn0025839 | CG3621 | ND-B14.5A | NADH dehydrogenase (ubiquinone) B14.5 A subunit<br>(SD16673p) | GO:0016491~oxidoreductase activity, | dme00190:Oxidative phosphorylation,<br>dme01100:Metabolic pathways, |

| | | | | | | |
|---|---|---|---|---|---|---|
| **P00334** | FBgn0000055 | CG3481 | Adh | Alcohol dehydrogenase | GO:0004022~alcohol dehydrogenase (NAD) activity,<br>GO:0008774~acetaldehyde dehydrogenase (acetylating) activity,<br>GO:0016491~oxidoreductase activity,<br>GO:0042803~protein homodimerization activity, | dme00010:Glycolysis / Gluconeogenesis,<br>dme00071:Fatty acid degradation,<br>dme00350:Tyrosine metabolism,<br>dme00620:Pyruvate metabolism,<br>dme00830:Retinol metabolism,<br>dme00980:Metabolism of xenobiotics by cytochrome P450,<br>dme00982:Drug metabolism - cytochrome P450,<br>dme01100:Metabolic pathways, |
| **P02574** | FBgn0000045 | CG7478 | Act79B | Actin 79B<br>(Actin; larval muscle) | GO:0005524~ATP binding,<br>GO:0016787~hydrolase activity, | |
| **P02828** | FBgn0001233 | CG1242 | Hsp83 | Heat shock protein 83 | GO:0005158~insulin receptor binding,<br>GO:0005515~protein binding,<br>GO:0005524~ATP binding,<br>GO:0016887~ATPase activity,<br>GO:0030911~TPR domain binding,<br>GO:0051082~unfolded protein binding,<br>GO:0097718~disordered domain specific binding, | dme04141:Protein processing in endoplasmic reticulum, |
| **P04359** | FBgn0002626 | CG7939 | RpL32 | 60S ribosomal protein L32<br>(Ribosomal protein L32) | GO:0003735~structural constituent of ribosome, | dme03010:Ribosome, |
| **P06605** | FBgn0003885 | CG2512 | alphaTub84D | alpha-Tubulin at 84D<br>(Tubulin alpha-3 chain) | GO:0005200~structural constituent of cytoskeleton,<br>GO:0005525~GTP binding,<br>GO:0016787~hydrolase activity,<br>GO:0017022~myosin binding,<br>GO:0046872~metal ion binding, | dme04145:Phagosome,<br>dme04814:Motor proteins, |
| **P06607** | FBgn0004047 | CG11129 | Yp3 | Yolk protein 3<br>(Vitellogenin-3) | GO:0016298~lipase activity,<br>GO:0017171~serine hydrolase activity, | |
| **P06754** | FBgn0003721 | CG4898 | Tm1 | Tropomyosin-1, isoforms 9A/A/B | GO:0003779~actin binding,<br>GO:0019894~kinesin binding,<br>GO:0019901~protein kinase binding,<br>GO:0051015~actin filament binding, | dme04814:Motor proteins, |
| **P07486** | FBgn0001091 | CG12055 | Gapdh1 | Glyceraldehyde-3-phosphate dehydrogenase 1 | GO:0004365~glyceraldehyde-3-phosphate dehydrogenase (NAD+) (phosphorylating) activity,<br>GO:0016620~oxidoreductase activity, acting on the aldehyde or oxo group of donors, NAD or NADP as acceptor,<br>GO:0035605~peptidyl-cysteine S-nitrosylase activity,<br>GO:0050661~NADP binding,<br>GO:0051287~NAD binding, | dme00010:Glycolysis / Gluconeogenesis,<br>dme01100:Metabolic pathways,<br>dme01200:Carbon metabolism,<br>dme01230:Biosynthesis of amino acids, |
| **P07487** | FBgn0001092 | CG8893 | Gapdh2 | Glyceraldehyde-3-phosphate dehydrogenase 2 | GO:0004365~glyceraldehyde-3-phosphate dehydrogenase (NAD+) (phosphorylating) activity,<br>GO:0016620~oxidoreductase activity, acting on the aldehyde or oxo group of donors, NAD or NADP as acceptor,<br>GO:0016740~transferase activity,<br>GO:0035605~peptidyl-cysteine S-nitrosylase activity,<br>GO:0050661~NADP binding,<br>GO:0051287~NAD binding, | dme00010:Glycolysis / Gluconeogenesis,<br>dme01100:Metabolic pathways,<br>dme01200:Carbon metabolism,<br>dme01230:Biosynthesis of amino acids, |
| **P07764** | FBgn0000064 | CG6058 | Ald1 | Aldolase 1<br>(Fructose-bisphosphate aldolase) | GO:0004332~fructose-bisphosphate aldolase activity, | dme00010:Glycolysis / Gluconeogenesis,<br>dme00030:Pentose phosphate pathway,<br>dme00051:Fructose and mannose metabolism,<br>dme01100:Metabolic pathways,<br>dme01200:Carbon metabolism,<br>dme01230:Biosynthesis of amino acids, |
| **P08736** | FBgn0284245 | CG8280 | eEF1alpha1 | Elongation factor 1-alpha 1<br>(eukaryotic translation elongation factor 1 alpha 1) | GO:0003746~translation elongation factor activity,<br>GO:0003924~GTPase activity,<br>GO:0005525~GTP binding, | dme03013:Nucleocytoplasmic transport, |
| **P08879** | FBgn0000150 | CG2210 | awd | abnormal wing discs<br>(Nucleoside diphosphate kinase) | GO:0000287~magnesium ion binding,<br>GO:0004550~nucleoside diphosphate kinase activity,<br>GO:0005524~ATP binding,<br>GO:0005525~GTP binding,<br>GO:0008017~microtubule binding,<br>GO:0016301~kinase activity, | dme00230:Purine metabolism,<br>dme00240:Pyrimidine metabolism,<br>dme00983:Drug metabolism - other enzymes,<br>dme01100:Metabolic pathways,<br>dme01232:Nucleotide metabolism,<br>dme01240:Biosynthesis of cofactors, |
| **P08928** | FBgn0002525 | CG6944 | Lam | Lamin Dm0 | GO:0003682~chromatin binding,<br>GO:0005102~receptor binding,<br>GO:0005200~structural constituent of cytoskeleton,<br>GO:0005515~protein binding, | dme04214:Apoptosis - fly, |
| **P09180** | FBgn0003279 | CG5502 | RpL4 | 60S ribosomal protein L4<br>(Ribosomal protein L4) | GO:0003723~RNA binding,<br>GO:0003735~structural constituent of ribosome, | dme03010:Ribosome, |
| **P09491** | FBgn0004117 | CG4843 | Tm2 | Tropomyosin-2 | GO:0051015~actin filament binding, | dme04814:Motor proteins, |
| **P11046** | FBgn0261800 | CG7123 | LanB1 | Laminin subunit beta-1 | | dme04512:ECM-receptor interaction, |
| **P11147** | FBgn0266599 | CG4264 | Hsc70-4 | Heat shock 70 kDa protein cognate 4 | GO:0005524~ATP binding,<br>GO:0016787~hydrolase activity,<br>GO:0016887~ATPase activity,<br>GO:0031072~heat shock protein binding,<br>GO:0044183~protein binding involved in protein folding,<br>GO:0051082~unfolded protein binding,<br>GO:0051087~chaperone binding,<br>GO:0051787~misfolded protein binding, | dme03040:Spliceosome,<br>dme04141:Protein processing in endoplasmic reticulum,<br>dme04144:Endocytosis,<br>dme04213:Longevity regulating pathway - multiple species, |
| **P12024** | FBgn0267435 | CG1744 | chp | Chaoptin | | |
| **P12982** | FBgn0004103 | CG5650 | Pp1-87B | Serine/threonine-protein phosphatase alpha-2 isoform<br>(Protein phosphatase 1 at 87B) | GO:0004722~protein serine/threonine phosphatase activity,<br>GO:0005515~protein binding,<br>GO:0016787~hydrolase activity,<br>GO:0017018~myosin phosphatase activity,<br>GO:0046872~metal ion binding, | dme03015:mRNA surveillance pathway, |
| **P13060** | FBgn0000559 | CG2238 | eEF2 | Elongation factor 2<br>(Eukaryotic translation elongation factor 2) | GO:0003746~translation elongation factor activity,<br>GO:0003924~GTPase activity,<br>GO:0005525~GTP binding,<br>GO:0043022~ribosome binding, | |
| **P13217** | FBgn0262738 | CG3620 | norpA | no receptor potential A<br>(1-phosphatidylinositol 4,5-bisphosphate phosphodiesterase) | GO:0004435~phosphatidylinositol phospholipase C activity,<br>GO:0004629~phospholipase C activity,<br>GO:0005509~calcium ion binding,<br>GO:0008081~phosphoric diester hydrolase activity, | dme00562:Inositol phosphate metabolism,<br>dme01100:Metabolic pathways,<br>dme04070:Phosphatidylinositol signaling system,<br>dme04310:Wnt signaling pathway,<br>dme04745:Phototransduction - fly, |
| **P13395** | FBgn0250789 | CG1977 | alpha-Spec | Spectrin alpha chain | GO:0003779~actin binding,<br>GO:0005509~calcium ion binding,<br>GO:0005516~calmodulin binding,<br>GO:0008017~microtubule binding,<br>GO:0051015~actin filament binding, | |
| **P15372** | FBgn0000120 | CG5711 | Arr1 | Arrestin 1<br>(Phosrestin-2) | GO:0001664~G-protein coupled receptor binding,<br>GO:0002046~opsin binding, | |
| **P17336** | FBgn0000261 | CG6871 | Cat | Catalase | GO:0004096~catalase activity,<br>GO:0020037~heme binding,<br>GO:0046872~metal ion binding, | dme00380:Tryptophan metabolism,<br>dme00630:Glyoxylate and dicarboxylate metabolism,<br>dme01100:Metabolic pathways,<br>dme01200:Carbon metabolism,<br>dme04068:FoxO signaling pathway,<br>dme04146:Peroxisome,<br>dme04213:Longevity regulating pathway - multiple species, |
| **P17704** | FBgn0005533 | CG3922 | RpS17 | 40S ribosomal protein S17<br>(Ribosomal protein S17) | GO:0003735~structural constituent of ribosome, | dme03010:Ribosome, |
| **P18930** | FBgn0013681 | CG34076 | mt:ND3 | Mitochondrial NADH-ubiquinone oxidoreductase chain 3 | GO:0008137~NADH dehydrogenase (ubiquinone) activity, | dme00190:Oxidative phosphorylation,<br>dme01100:Metabolic pathways, |
| **P19967** | FBgn0000406 | CG13279 | Cyt-b5-r | Cytochrome b5-related protein | GO:0016491~oxidoreductase activity,<br>GO:0020037~heme binding,<br>GO:0046872~metal ion binding, | |
| **P20228** | FBgn0004516 | CG14994 | Gad1 | Glutamate decarboxylase,<br>(Glutamic acid decarboxylase 1) | GO:0004351~glutamate decarboxylase activity,<br>GO:0016830~carbon-carbon lyase activity,<br>GO:0016831~carboxy-lyase activity,<br>GO:0030170~pyridoxal phosphate binding, | dme00250:Alanine, aspartate and glutamate metabolism,<br>dme00410:beta-Alanine metabolism,<br>dme00430:Taurine and hypotaurine metabolism,<br>dme00650:Butanoate metabolism,<br>dme01100:Metabolic pathways, |
| **P20432** | FBgn0001149 | CG10045 | GstD1 | Glutathione S-transferase D1 | GO:0004364~glutathione transferase activity,<br>GO:0004602~glutathione peroxidase activity,<br>GO:0018833~DDT-dehydrochlorinase activity, | dme00480:Glutathione metabolism,<br>dme00980:Metabolism of xenobiotics by cytochrome P450,<br>dme00982:Drug metabolism - cytochrome P450,<br>dme00983:Drug metabolism - other enzymes,<br>dme01100:Metabolic pathways, |
| **P20478** | FBgn0001145 | CG1743 | Gs2 | Glutamine synthetase 2 cytoplasmic | GO:0003824~catalytic activity,<br>GO:0004356~glutamate-ammonia ligase activity,<br>GO:0005524~ATP binding, | dme00220:Arginine biosynthesis,<br>dme00250:Alanine, aspartate and glutamate metabolism,<br>dme00630:Glyoxylate and dicarboxylate metabolism,<br>dme00910:Nitrogen metabolism,<br>dme01100:Metabolic pathways,<br>dme01230:Biosynthesis of amino acids, |

| | | | | | | |
|---|---|---|---|---|---|---|
| **P21914** | FBgn0014028 | CG3283 | SdhB | Succinate dehydrogenase [ubiquinone] iron-sulfur subunit; mitochondrial<br>(Succinate dehydrogenase, subunit B (iron-sulfur)) | GO:0008177~succinate dehydrogenase (ubiquinone) activity,<br>GO:0009055~electron carrier activity,<br>GO:0016491~oxidoreductase activity,<br>GO:0046872~metal ion binding,<br>GO:0048039~ubiquinone binding,<br>GO:0051536~iron-sulfur cluster binding,<br>GO:0051537~2 iron, 2 sulfur cluster binding,<br>GO:0051538~3 iron, 4 sulfur cluster binding,<br>GO:0051539~4 iron, 4 sulfur cluster binding, | dme00020:Citrate cycle (TCA cycle),<br>dme00190:Oxidative phosphorylation,<br>dme01100:Metabolic pathways,<br>dme01200:Carbon metabolism, |
| **P22464** | FBgn0000083 | CG5730 | AnxB9 | Annexin B9 | GO:0001786~phosphatidylserine binding,<br>GO:0005509~calcium ion binding,<br>GO:0005515~protein binding,<br>GO:0005544~calcium-dependent phospholipid binding,<br>GO:0030507~spectrin binding, | |
| **P22700** | FBgn0263006 | CG3725 | SERCA | Sarco/endoplasmic reticulum Ca(2+)-ATPase<br>(Calcium-transporting ATPase sarcoplasmic/endoplasmic reticulum type) | GO:0000166~nucleotide binding,<br>GO:0005388~calcium-transporting ATPase activity,<br>GO:0005515~protein binding,<br>GO:0005524~ATP binding,<br>GO:0008553~hydrogen-exporting ATPase activity, phosphorylative mechanism,<br>GO:0015662~ATPase activity, coupled to transmembrane movement of ions, phosphorylative mechanism,<br>GO:0016887~ATPase activity,<br>GO:0046872~metal ion binding, | |
| **P25007** | FBgn0004432 | CG9916 | Cyp1 | Cyclophilin 1<br>(Peptidyl-prolyl cis-trans isomerase) | GO:0003755~peptidyl-prolyl cis-trans isomerase activity,<br>GO:0016018~cyclosporin A binding, | |
| **P29310** | FBgn0004907 | CG17870 | 14-3-3zeta | 14-3-3 protein zeta | GO:0001223~transcription coactivator binding,<br>GO:0005515~protein binding,<br>GO:0008134~transcription factor binding,<br>GO:0042803~protein homodimerization activity,<br>GO:0046982~protein heterodimerization activity, | dme04013:MAPK signaling pathway - fly,<br>dme04391:Hippo signaling pathway - fly, |
| **P29413** | FBgn0005585 | CG9429 | Calr | Calreticulin | GO:0005509~calcium ion binding,<br>GO:0030246~carbohydrate binding,<br>GO:0051082~unfolded protein binding, | dme04141:Protein processing in endoplasmic reticulum,<br>dme04145:Phagosome,<br>dme04148:Efferocytosis, |
| **P29613** | FBgn0086355 | CG2171 | Tpi | Triose phosphate isomerase | GO:0004807~triose-phosphate isomerase activity,<br>GO:0042803~protein homodimerization activity, | dme00010:Glycolysis / Gluconeogenesis,<br>dme00051:Fructose and mannose metabolism,<br>dme00562:Inositol phosphate metabolism,<br>dme01100:Metabolic pathways,<br>dme01200:Carbon metabolism,<br>dme01230:Biosynthesis of amino acids, |
| **P29742** | FBgn0000319 | CG9012 | Chc | Clathrin heavy chain | GO:0005198~structural molecule activity,<br>GO:0005515~protein binding,<br>GO:0032051~clathrin light chain binding, | dme04142:Lysosome,<br>dme04144:Endocytosis, |
| **P29829** | FBgn0004623 | CG8770 | Gbeta76C | Guanine nucleotide-binding protein subunit beta-2<br>(G protein beta-subunit 76C) | GO:0016004~phospholipase activator activity,<br>GO:0030159~receptor signaling complex scaffold activity,<br>GO:0031682~G-protein gamma-subunit binding,<br>GO:0046982~protein heterodimerization activity, | dme04745:Phototransduction - fly, |
| **P31009** | FBgn0004867 | CG5920 | RpS2 | 40S ribosomal protein S2<br>(Ribosomal protein S2) | GO:0003723~RNA binding,<br>GO:0003735~structural constituent of ribosome, | dme03010:Ribosome, |
| **P31409** | FBgn0005671 | CG17369 | Vha55 | V-type proton ATPase subunit B | GO:0005524~ATP binding,<br>GO:0016787~hydrolase activity,<br>GO:0046961~proton-transporting ATPase activity, rotational mechanism, | dme00190:Oxidative phosphorylation,<br>dme01100:Metabolic pathways,<br>dme04145:Phagosome,<br>dme04150:mTOR signaling pathway, |
| **P35381** | FBgn0011211 | CG3612 | blw | ATP synthase subunit alpha; mitochondrial<br>(bellwether) | GO:0005524~ATP binding,<br>GO:0032559~adenyl ribonucleotide binding,<br>GO:0043531~ADP binding,<br>GO:0046933~proton-transporting ATP synthase activity, rotational mechanism, | dme00190:Oxidative phosphorylation,<br>dme01100:Metabolic pathways, |
| **P35415** | FBgn0003149 | CG5939 | Prm | Paramyosin; long form | GO:0016740~transferase activity, | |
| **P38979** | FBgn0003517 | CG14792 | sta<br>(RpSA) | 40S ribosomal protein SA<br>(Ribosomal protein SA)<br>(stubarista) | GO:0003735~structural constituent of ribosome,<br>GO:0043022~ribosome binding, | dme03010:Ribosome, |
| **P39018** | FBgn0010412 | CG4464 | RpS19a | 40S ribosomal protein S19a<br>(Ribosomal protein S19a) | GO:0003723~RNA binding,<br>GO:0003735~structural constituent of ribosome, | dme03010:Ribosome, |
| **P46461** | FBgn0000346 | CG1618 | comt | Vesicle-fusing ATPase 1<br>(comatose) | GO:0005524~ATP binding,<br>GO:0016887~ATPase activity,<br>GO:0046872~metal ion binding, | |
| **P48148** | FBgn0014020 | CG8416 | Rho1 | Ras-like GTP-binding protein Rho1 | GO:0003779~actin binding,<br>GO:0003924~GTPase activity,<br>GO:0005515~protein binding,<br>GO:0005525~GTP binding,<br>GO:0019900~kinase binding,<br>GO:0019901~protein kinase binding, | dme04144:Endocytosis,<br>dme04150:mTOR signaling pathway,<br>dme04310:Wnt signaling pathway,<br>dme04350:TGF-beta signaling pathway, |
| **P48375** | FBgn0013954 | CG11001 | Fkbp12 | FK506-binding protein 12 kDa<br>(Peptidyl-prolyl cis-trans isomerase Fkbp12) | GO:0003755~peptidyl-prolyl cis-trans isomerase activity,<br>GO:0034713~type I transforming growth factor beta receptor binding, | |
| **P48554** | FBgn0014011 | CG8556 | Rac2 | Ras-related protein Rac2 | GO:0003924~GTPase activity,<br>GO:0005515~protein binding,<br>GO:0005525~GTP binding,<br>GO:0019901~protein kinase binding, | dme04013:MAPK signaling pathway - fly,<br>dme04145:Phagosome,<br>dme04148:Efferocytosis,<br>dme04310:Wnt signaling pathway, |
| **P48602** | FBgn0265262 | CG12403 | Vha68-1 | V-type proton ATPase catalytic subunit A isoform 1<br>(Vacuolar H+ ATPase 68kD subunit 1) | GO:0003824~catalytic activity,<br>GO:0005524~ATP binding,<br>GO:0016887~ATPase activity,<br>GO:0046933~proton-transporting ATP synthase activity, rotational mechanism,<br>GO:0046961~proton-transporting ATPase activity, rotational mechanism, | dme00190:Oxidative phosphorylation,<br>dme01100:Metabolic pathways,<br>dme04145:Phagosome,<br>dme04150:mTOR signaling pathway, |
| **P48610** | FBgn0000116 | CG32031 | Argk1 | Arginine kinase | GO:0004054~arginine kinase activity,<br>GO:0004111~creatine kinase activity,<br>GO:0005524~ATP binding,<br>GO:0016301~kinase activity,<br>GO:0016772~transferase activity, transferring phosphorus-containing groups, | dme00330:Arginine and proline metabolism, |
| **P49455** | FBgn0003721 | CG4898 | Tm1 | Tropomyosin-1, isoforms 33/34 | GO:0003779~actin binding,<br>GO:0019894~kinesin binding,<br>GO:0019901~protein kinase binding,<br>GO:0051015~actin filament binding, | dme04814:Motor proteins, |
| **P50882** | FBgn0015756 | CG6141 | RpL9 | 60S ribosomal protein L9<br>(Ribosomal protein L9) | GO:0003735~structural constituent of ribosome,<br>GO:0019843~rRNA binding, | dme03010:Ribosome, |
| **P52029** | FBgn0003074 | CG8251 | Pgi | Phosphoglucose isomerase<br>(Glucose-6-phosphate isomerase) | GO:0004347~glucose-6-phosphate isomerase activity,<br>GO:0048029~monosaccharide binding,<br>GO:0097367~carbohydrate derivative binding, | dme00010:Glycolysis / Gluconeogenesis,<br>dme00030:Pentose phosphate pathway,<br>dme00500:Starch and sucrose metabolism,<br>dme00520:Amino sugar and nucleotide sugar metabolism,<br>dme01100:Metabolic pathways,<br>dme01200:Carbon metabolism,<br>dme01250:Biosynthesis of nucleotide sugars, |
| **P52034** | FBgn0003071 | CG4001 | Pfk | 6-phosphofructokinase<br>(ATP-dependent 6-phosphofructokinase) | GO:0003677~DNA binding,<br>GO:0003872~6-phosphofructokinase activity,<br>GO:0005524~ATP binding,<br>GO:0016208~AMP binding,<br>GO:0042802~identical protein binding,<br>GO:0046872~metal ion binding,<br>GO:0048029~monosaccharide binding,<br>GO:0070095~fructose-6-phosphate binding, | dme00010:Glycolysis / Gluconeogenesis,<br>dme00030:Pentose phosphate pathway,<br>dme00051:Fructose and mannose metabolism,<br>dme00052:Galactose metabolism,<br>dme01100:Metabolic pathways,<br>dme01200:Carbon metabolism,<br>dme01230:Biosynthesis of amino acids,<br>dme03018:RNA degradation, |
| **P53777** | FBgn0259209 | CG42309 | Mlp60A | Muscle LIM protein 1<br>(Muscle LIM protein at 60A) | GO:0046872~metal ion binding, | |
| **P54385** | FBgn0001098 | CG5320 | Gdh | Glutamate dehydrogenase, mitochondrial | GO:0000166~nucleotide binding,<br>GO:0004352~glutamate dehydrogenase (NAD+) activity,<br>GO:0004353~glutamate dehydrogenase [NAD(P)+] activity,<br>GO:0004354~glutamate dehydrogenase (NADP+) activity,<br>GO:0005524~ATP binding,<br>GO:0005525~GTP binding,<br>GO:0016639~oxidoreductase activity, acting on the CH-NH2 group of donors, NAD or NADP as acceptor,<br>GO:0042802~identical protein binding, | dme00220:Arginine biosynthesis,<br>dme00250:Alanine, aspartate and glutamate metabolism,<br>dme00910:Nitrogen metabolism,<br>dme01100:Metabolic pathways,<br>dme01200:Carbon metabolism, |
| **P54611** | FBgn0283535 | CG1088 | Vha26 | V-type proton ATPase subunit E<br>(Vacuolar $H^+$-ATPase 26kD subunit) | GO:0008553~hydrogen-exporting ATPase activity, phosphorylative mechanism,<br>GO:0046961~proton-transporting ATPase activity, rotational mechanism, | dme00190:Oxidative phosphorylation,<br>dme01100:Metabolic pathways,<br>dme04145:Phagosome,<br>dme04150:mTOR signaling pathway, |
| **P55828** | FBgn0019936 | CG15693 | RpS20 | 40S ribosomal protein S20<br>(Ribosomal protein S20) | GO:0003723~RNA binding,<br>GO:0003735~structural constituent of ribosome, | dme03010:Ribosome, |
| **P55830** | FBgn0017545 | CG2168 | RpS3A | 40S ribosomal protein S3a<br>(Ribosomal protein S3A) | GO:0003735~structural constituent of ribosome, | dme03010:Ribosome, |

| | | | | | | |
|---|---|---|---|---|---|---|
| **P61851** | FBgn0003462 | CG11793 | Sod1 | Superoxide dismutase 1 [Cu-Zn] | GO:0004784~superoxide dismutase activity,<br>GO:0005507~copper ion binding,<br>GO:0042803~protein homodimerization activity,<br>GO:0046872~metal ion binding, | dme04146:Peroxisome,<br>dme04213:Longevity regulating pathway - multiple species, |
| **P80455** | FBgn0286213 | CG11271 | RpS12 | 40S ribosomal protein S12<br>(Ribosomal protein S12) | GO:0003735~structural constituent of ribosome, | dme03010:Ribosome, |
| **P83967** | FBgn0000047 | CG5178 | Act88F | Actin; indirect flight muscle<br>(Actin 88F) | GO:0005524~ATP binding,<br>GO:0016787~hydrolase activity, | |
| **P91929** | FBgn0019957 | CG6343 | ND-42 | NADH dehydrogenase (ubiquinone) 42 kDa subunit<br>(NADH dehydrogenase [ubiquinone]1 alpha subcomplex subunit 10; mitochondrial) | GO:0005515~protein binding,<br>GO:0016491~oxidoreductase activity, | dme00190:Oxidative phosphorylation,<br>dme01100:Metabolic pathways, |
| **P91938** | FBgn0020653 | CG2151 | Trxr1 | Thioredoxin reductase 1, mitochondrial | GO:0004791~thioredoxin-disulfide reductase activity,<br>GO:0016209~antioxidant activity,<br>GO:0016491~oxidoreductase activity,<br>GO:0016668~oxidoreductase activity, acting on a sulfur group of donors, NAD(P) as acceptor<br>GO:0042803~protein homodimerization activity,<br>GO:0050660~flavin adenine dinucleotide binding, | dme00450:Selenocompound metabolism, |
| **P92177** | FBgn0020238 | CG31196 | 14-3-3epsilon | 14-3-3 protein epsilon | GO:0001223~transcription coactivator binding,<br>GO:0005515~protein binding,<br>GO:0008134~transcription factor binding,<br>GO:0046982~protein heterodimerization activity,<br>GO:0050815~phosphoserine binding, | dme04391:Hippo signaling pathway - fly, |
| **Q00637** | FBgn0010213 | CG8905 | Sod2 | Superoxide dismutase 2 [Mn]; mitochondrial | GO:0004784~superoxide dismutase activity,<br>GO:0016209~antioxidant activity,<br>GO:0030145~manganese ion binding,<br>GO:0046872~metal ion binding, | dme04013:MAPK signaling pathway - fly,<br>dme04068:FoxO signaling pathway,<br>dme04146:Peroxisome,<br>dme04213:Longevity regulating pathway - multiple species, |
| **Q02645** | FBgn0263391 | CG9325 | hts | Protein hu-li tai shao | GO:0051015~actin filament binding, | |
| **Q02748** | FBgn0001942 | CG9075 | eIF4A | Eukaryotic translation initiation factor 4A | GO:0003676~nucleic acid binding,<br>GO:0003723~RNA binding,<br>GO:0003724~RNA helicase activity,<br>GO:0003743~translation initiation factor activity,<br>GO:0005524~ATP binding,<br>GO:0016787~hydrolase activity,<br>GO:0016887~ATP hydrolysis activity,<br>GO:0046332~SMAD binding, | |
| **Q07327** | FBgn0004574 | CG15811 | Rop | Ras opposite (Protein ROP) | GO:0000149~SNARE binding,<br>GO:0019905~syntaxin binding, | |
| **Q0E8V7** | FBgn0083968 | CG34132 | CG34132 | uncharacterized protein CG34132<br>(Mitochondrial import inner membrane translocase subunit)<br>(IP17740p) | GO:0046872~metal ion binding, | |
| **Q0KIA8** | FBgn0037440 | CG1041 | CRAT | Carnitine O-Acetyl-Transferase<br>(CG1041; isoform B) | GO:0004092~carnitine O-acetyltransferase activity,<br>GO:0016746~transferase activity, transferring acyl groups, | dme04146:Peroxisome, |
| **Q23997** | FBgn0013763 | CG5210 | Idgf6 | Imaginal disc growth factor 6<br>(Chitinase-like protein CG5210) | GO:0008061~chitin binding,<br>GO:0008084~imaginal disc growth factor receptor binding, | |
| **Q24048** | FBgn0015777 | CG9261 | nrv2 | Nervana 2<br>(Sodium/potassium-transporting ATPase subunit beta-2) | GO:0001671~ATPase activator activity, | |
| **Q24186** | FBgn0002590 | CG8922 | RpS5a | 40S ribosomal protein S5a<br>(Ribosomal protein S5a) | GO:0003723~RNA binding,<br>GO:0003729~mRNA binding,<br>GO:0003735~structural constituent of ribosome,<br>GO:0019843~rRNA binding, | dme03010:Ribosome, |
| **Q24251** | FBgn0016120 | CG6030 | ATPsynD | ATP synthase subunit d, mitochondrial | GO:0015078~hydrogen ion transmembrane transporter activity,<br>GO:0046933~proton-transporting ATP synthase activity, rotational | dme00190:Oxidative phosphorylation,<br>dme01100:Metabolic pathways, |
| **Q24253** | FBgn0010380 | CG12532 | AP-1-2beta | AP complex subunit beta<br>(Adaptor Protein complex 1/2, beta subunit)<br>(LP17054p) | GO:0030276~clathrin binding,<br>GO:0035615~clathrin adaptor activity, | dme04142:Lysosome, |
| **Q24400** | FBgn0014863 | CG1019 | Mlp84B | Muscle LIM protein at 84B | GO:0008307~structural constituent of muscle,<br>GO:0042805~actinin binding,<br>GO:0046872~metal ion binding, | |
| **Q24498** | FBgn0011286 | CG10844 | RyR | Ryanodine receptor 44F | GO:0005216~monoatomic ion channel activity,<br>GO:0005219~ryanodine-sensitive calcium-release channel activity,<br>GO:0005509~calcium ion binding,<br>GO:0048763~calcium-induced calcium release activity, | |
| **Q24560** | FBgn0284243 | CG9277 | betaTub56D | Tubulin beta-1 chain<br>(beta-tubulin at 56D) | GO:0003924~GTPase activity,<br>GO:0005200~structural constituent of cytoskeleton,<br>GO:0005525~GTP binding,<br>GO:0046872~metal ion binding, | dme04145:Phagosome,<br>dme04814:Motor proteins, |
| **Q26365** | FBgn0003360 | CG16944 | sesB | stress-sensitive B<br>(ADP,ATP carrier protein) | GO:0005471~ATP:ADP antiporter activity, | |
| **Q27268** | FBgn0014189 | CG7269 | Hel25E | ATP-dependent RNA helicase WM6<br>(Helicase at 25E) | GO:0003676~nucleic acid binding,<br>GO:0003723~RNA binding,<br>GO:0003724~RNA helicase activity,<br>GO:0003729~mRNA binding,<br>GO:0005524~ATP binding,<br>GO:0016787~hydrolase activity, | dme03013:Nucleocytoplasmic transport,<br>dme03015:mRNA surveillance pathway,<br>dme03040:Spliceosome, |
| **Q59DP8** | FBgn0259214 | CG42314 | PMCA | Plasma membrane calcium ATPase; isoform N<br>(Calcium-transporting ATPase) | GO:0000166~nucleotide binding,<br>GO:0005388~calcium-transporting ATPase activity,<br>GO:0005524~ATP binding,<br>GO:0016887~ATPase activity,<br>GO:0019829~cation-transporting ATPase activity, | |
| **Q59E65<br>(A8DYP0)** | FBgn0053519 | CG33519 | Unc-89 | Obscurin<br>(Unc-89; isoform B) | GO:0005085~guanyl-nucleotide exchange factor activity,<br>GO:0005515~protein binding,<br>GO:0005523~tropomyosin binding,<br>GO:0005524~ATP binding,<br>GO:0019901~protein kinase binding, | |
| **Q7JS69** | FBgn0032946 | CG8663 | nrv3 | Nervana 3; isoform D<br>(Na/K-ATPase beta subunit isoform 3)<br>(FI04632p) | GO:0001671~ATPase activator activity, | |
| **Q7JUS9** | FBgn0034497 | CG9090 | Mpcp1 | Mitochondrial phosphate carrier protein 1<br>(RH64567p) | GO:0005315~inorganic phosphate transmembrane transporter activity, | |
| **Q7JVH6** | FBgn0033101 | CG9436 | CG9436 | uncharacterized protein CG9436 (LD24696p) | GO:0004032~alditol:NADP+ 1-oxidoreductase activity,<br>GO:0016491~oxidoreductase activity,<br>GO:0047834~D-threo-aldose 1-dehydrogenase activity, | dme00040:Pentose and glucuronate interconversions,<br>dme00051:Fructose and mannose metabolism,<br>dme00052:Galactose metabolism,<br>dme00561:Glycerolipid metabolism,<br>dme00790:Folate biosynthesis,<br>dme01100:Metabolic pathways, |
| **Q7JXB5** | FBgn0034356 | CG10924 | Pepck2 | Phosphoenolpyruvate carboxykinase 2<br>(RE12569p) | GO:0004611~phosphoenolpyruvate carboxykinase activity,<br>GO:0004613~phosphoenolpyruvate carboxykinase (GTP) activity,<br>GO:0005525~GTP binding,<br>GO:0016301~kinase activity,<br>GO:0017076~purine nucleotide binding,<br>GO:0030145~manganese ion binding, | dme00010:Glycolysis / Gluconeogenesis,<br>dme00020:Citrate cycle (TCA cycle),<br>dme00620:Pyruvate metabolism,<br>dme01100:Metabolic pathways,<br>dme04068:FoxO signaling pathway, |
| **Q7K1C0** | FBgn0031228 | CG11455 | ND-15 | NADH dehydrogenase (ubiquinone) 15 kDa subunit<br>(MIP04232p) | | dme00190:Oxidative phosphorylation,<br>dme01100:Metabolic pathways, |
| **Q7K1Q6** | FBgn0259682 | CG42351 | Jabba | Jabba; isoform B<br>(LD47410p) | | |
| **Q7K4T8** | FBgn0033734 | CG8520 | CG8520 | uncharacterized protein CG8520 (LD23856p) | GO:0005524~ATP binding,<br>GO:0016787~hydrolase activity,<br>GO:0016887~ATPase activity, | |
| **Q7K569** | FBgn0022160 | CG8256 | Gpo1 | Glycerophosphate oxidase-1; isoform D<br>(Glycerol-3-phosphate dehydrogenase) | GO:0004368~glycerol-3-phosphate dehydrogenase activity,<br>GO:0005509~calcium ion binding,<br>GO:0016491~oxidoreductase activity,<br>GO:0052591~sn-glycerol-3-phosphate:ubiquinone-8 oxidoreductase activity, | dme00564:Glycerophospholipid metabolism, |
| **Q7K5K3** | FBgn0039635 | CG11876 | Pdhb | Pyruvate dehydrogenase E1 subunit beta, mitochondrial<br>(GH08474p) | GO:0003824~catalytic activity,<br>GO:0004739~pyruvate dehydrogenase (acetyl-transferring) activity, | dme00010:Glycolysis / Gluconeogenesis,<br>dme00020:Citrate cycle (TCA cycle),<br>dme00620:Pyruvate metabolism,<br>dme00785:Lipoic acid metabolism,<br>dme01100:Metabolic pathways,<br>dme01200:Carbon metabolism,<br>dme01210:2-Oxocarboxylic acid metabolism, |
| **Q7K860** | FBgn0033027 | CG12408 | TpnC4 | Troponin C isoform 4; isoform B | GO:0005509~calcium ion binding,<br>GO:0030234~enzyme regulator activity, | |
| **Q7KN62** | FBgn0286784 | CG2331 | TER94 | Transitional endoplasmic reticulum ATPase TER94 | GO:0005515~protein binding,<br>GO:0005524~ATP binding,<br>GO:0016887~ATPase activity, | dme04137:Mitophagy - animal,<br>dme04141:Protein processing in endoplasmic reticulum, |

| | | | | | | |
|---|---|---|---|---|---|---|
| **Q7KN75** | FBgn0027835 | CG5170 | Dp1 | Dodeca-satellite-binding protein 1; isoform F | GO:0003676~nucleic acid binding,<br>GO:0003696~satellite DNA binding,<br>GO:0003697~single-stranded DNA binding,<br>GO:0003723~RNA binding,<br>GO:0003729~mRNA binding,<br>GO:0003730~mRNA 3'-UTR binding, | |
| **Q7KN97** | FBgn0027580 | CG1516 | PCB | Pyruvate carboxylase | GO:0003824~catalytic activity,<br>GO:0004736~pyruvate carboxylase activity,<br>GO:0005524~ATP binding,<br>GO:0009374~biotin binding,<br>GO:0046872~metal ion binding, | dme00020:Citrate cycle (TCA cycle),<br>dme00620:Pyruvate metabolism,<br>dme01100:Metabolic pathways,<br>dme01200:Carbon metabolism,<br>dme01230:Biosynthesis of amino acids, |
| **Q7KR04** | FBgn0033555 | CG12324 | RpS15Ab | 40S ribosomal protein S15Ab<br>(Ribosomal protein S15Ab) | GO:0003735~structural constituent of ribosome, | dme03010:Ribosome, |
| **Q7KSQ0** | FBgn0037912 | CG6782 | sea | Citrate transport protein<br>(scheggia)<br>(RH27308p) | GO:0071913~citrate secondary active transmembrane transporter activity, | |
| **Q7KSU6** | FBgn0037607 | CG8036 | CG8036 | Transketolase<br>(CG8036; isoform D) | GO:0003824~catalytic activity,<br>GO:0004802~transketolase activity,<br>GO:0030976~thiamine pyrophosphate binding,<br>GO:0046872~metal ion binding, | dme00030:Pentose phosphate pathway,<br>dme01100:Metabolic pathways,<br>dme01200:Carbon metabolism,<br>dme01230:Biosynthesis of amino acids, |
| **Q7KTK9** | FBgn0283658 | CG5261 | Dlat<br>(muc) | Dihydrolipoamide S-acetyltransferase<br>(midline uncoordinated)<br>(CG5261; isoform A) | GO:0004742~dihydrolipoyllysine-residue acetyltransferase activity,<br>GO:0016746~transferase activity, transferring acyl groups, | dme00010:Glycolysis / Gluconeogenesis,<br>dme00020:Citrate cycle (TCA cycle),<br>dme00620:Pyruvate metabolism,<br>dme00785:Lipoic acid metabolism,<br>dme01100:Metabolic pathways,<br>dme01200:Carbon metabolism,<br>dme01210:2-Oxocarboxylic acid metabolism, |
| **Q7KTW5** | FBgn0037063 | CG9391 | CG9391 | Inositol-1-monophosphatase<br>(RE10407p) | GO:0008934~inositol monophosphate 1-phosphatase activity,<br>GO:0046872~metal ion binding,<br>GO:0052832~inositol monophosphate 3-phosphatase activity,<br>GO:0052833~inositol monophosphate 4-phosphatase activity, | dme00562:Inositol phosphate metabolism,<br>dme01100:Metabolic pathways,<br>dme04070:Phosphatidylinositol signaling system, |
| **Q7KUB0** | FBgn0001248 | CG7176 | Idh | Isocitrate dehydrogenase [NADP] | GO:0000287~magnesium ion binding,<br>GO:0004448~isocitrate dehydrogenase activity,<br>GO:0004450~isocitrate dehydrogenase (NADP+) activity,<br>GO:0016616~oxidoreductase activity, acting on the CH-OH group of donors, NAD or NADP as acceptor,<br>GO:0051287~NAD binding, | dme00020:Citrate cycle (TCA cycle),<br>dme00480:Glutathione metabolism,<br>dme01100:Metabolic pathways,<br>dme01200:Carbon metabolism,<br>dme01210:2-Oxocarboxylic acid metabolism,<br>dme01230:Biosynthesis of amino acids,<br>dme04146:Peroxisome, |
| **Q7KUQ5** | FBgn0261565 | CG42679 | Lmpt | Limpet; isoform F | GO:0008270~zinc ion binding,<br>GO:0046872~metal ion binding, | |
| **Q7KV27** | FBgn0030478 | CG1640 | Alat<br>(ALT) | Alanine transaminase<br>(CG1640; isoform F) | GO:0003824~catalytic activity,<br>GO:0004021~L-alanine:2-oxoglutarate aminotransferase activity,<br>GO:0008483~transaminase activity,<br>GO:0030170~pyridoxal phosphate binding, | dme00220:Arginine biosynthesis,<br>dme00250:Alanine, aspartate and glutamate metabolism,<br>dme01100:Metabolic pathways,<br>dme01200:Carbon metabolism,<br>dme01210:2-Oxocarboxylic acid metabolism,<br>dme01230:Biosynthesis of amino acids, |
| **Q7KVP4** | FBgn0034618 | CG9485 | Agl | Glycogen debranching enzyme<br>(CG9485; isoform E) | GO:0004134~4-alpha-glucanotransferase activity,<br>GO:0004135~amylo-alpha-1,6-glucosidase activity, | dme00500:Starch and sucrose metabolism,<br>dme01100:Metabolic pathways, |
| **Q7YU05** | FBgn0028325 | CG7010 | Pdha | Pyruvate dehydrogenase E1 alpha subunit<br>(Lethal (1) G0334; isoform B) | GO:0004739~pyruvate dehydrogenase (acetyl-transferring) activity,<br>GO:0016624~oxidoreductase activity, acting on the aldehyde or oxo group of donors, disulfide as acceptor, | dme00010:Glycolysis / Gluconeogenesis,<br>dme00020:Citrate cycle (TCA cycle),<br>dme00620:Pyruvate metabolism,<br>dme00785:Lipoic acid metabolism,<br>dme01100:Metabolic pathways,<br>dme01200:Carbon metabolism,<br>dme01210:2-Oxocarboxylic acid metabolism, |
| **Q868Z9** | FBgn0003137 | CG33103 | Ppn | Papilin | GO:0004222~metalloendopeptidase activity,<br>GO:0004867~serine-type endopeptidase inhibitor activity,<br>GO:0005201~extracellular matrix structural constituent,<br>GO:0030414~peptidase inhibitor activity, | |
| **Q86BQ4** | FBgn0031459 | CG2862 | HINT1 | Histidine triad nucleotide binding protein 1<br>(CG2862; isoform B) | GO:0003824~catalytic activity,<br>GO:0016787~hydrolase activity,<br>GO:0043530~adenosine 5'-monophosphoramidase activity, | |
| **Q8I0E9** | FBgn0001216 | CG8937 | Hsc70-1 | Heat shock protein 70 cognate 1; isoform D | GO:0005524~ATP binding,<br>GO:0016787~hydrolase activity,<br>GO:0016887~ATPase activity,<br>GO:0031072~heat shock protein binding,<br>GO:0044183~protein binding involved in protein folding,<br>GO:0051082~unfolded protein binding,<br>GO:0051787~misfolded protein binding, | dme03040:Spliceosome,<br>dme04141:Protein processing in endoplasmic reticulum,<br>dme04144:Endocytosis,<br>dme04213:Longevity regulating pathway - multiple species, |
| **Q8INE6** | FBgn0016691 | CG4307 | Oscp<br>(ATPsynO) | Oligomycin sensitivity-conferring protein; isoform B<br>(ATP synthase subunit O, mitochondrial) | GO:0046933~proton-transporting ATP synthase activity, rotational mechanism, | dme00190:Oxidative phosphorylation,<br>dme01100:Metabolic pathways, |
| **Q8IPE8** | FBgn0028479 | CG4389 | Mtpalpha | Mitochondrial trifunctional protein alpha subunit<br>(enoyl-CoA hydratase)<br>(CG4389; isoform C) | GO:0003824~catalytic activity,<br>GO:0003857~3-hydroxyacyl-CoA dehydrogenase activity,<br>GO:0004300~enoyl-CoA hydratase activity,<br>GO:0016491~oxidoreductase activity,<br>GO:0016509~long-chain-3-hydroxyacyl-CoA dehydrogenase activity,<br>GO:0016616~oxidoreductase activity, acting on the CH-OH group of donors, NAD or NADP as acceptor,<br>GO:0070403~NAD+ binding, | dme00062:Fatty acid elongation,<br>dme00071:Fatty acid degradation,<br>dme00280:Valine, leucine and isoleucine degradation,<br>dme00310:Lysine degradation,<br>dme00380:Tryptophan metabolism,<br>dme00410:beta-Alanine metabolism,<br>dme00640:Propanoate metabolism,<br>dme00650:Butanoate metabolism,<br>dme01100:Metabolic pathways,<br>dme01212:Fatty acid metabolism, |
| **Q8IPG9** | FBgn0261822 | CG31605 | Bsg | Basigin<br>(LD19437p) | GO:0098632~protein binding involved in cell-cell adhesion, | |
| **Q8IPP8** | FBgn0051548 | CG31548 | CG31548 | uncharacterized protein CG31548 | GO:0004303~estradiol 17-beta-dehydrogenase activity,<br>GO:0016491~oxidoreductase activity,<br>GO:0047045~testosterone 17-beta-dehydrogenase (NADP+) activity, | |
| **Q8IQ53** | FBgn0035601 | CG10640 | Uev1A | Ubiquitin-conjugating enzyme variant 1A; isoform B | GO:0031624~ubiquitin conjugating enzyme binding, | dme04624:Toll and Imd signaling pathway, |
| **Q8IQQ0** | FBgn0010352 | CG11661 | Ogdh<br>(Nc73EF) | Oxoglutarate dehydrogenase, mitochondrial<br>(Neural conserved at 73EF)<br>(RE42354p) | GO:0004591~oxoglutarate dehydrogenase (succinyl-transferring) activity,<br>GO:0016624~oxidoreductase activity, acting on the aldehyde or oxo group of donors, disulfide as acceptor,<br>GO:0030976~thiamine pyrophosphate binding,<br>GO:0034602~oxoglutarate dehydrogenase (NAD+) activity, | dme00020:Citrate cycle (TCA cycle),<br>dme00785:Lipoic acid metabolism,<br>dme01100:Metabolic pathways,<br>dme01200:Carbon metabolism,<br>dme01210:2-Oxocarboxylic acid metabolism, |
| **Q8IQW2** | FBgn0031066 | CG14235 | COX6B | Cytochrome c oxidase subunit 6B<br>(RE29690p) | GO:0016491~oxidoreductase activity, | dme00190:Oxidative phosphorylation,<br>dme01100:Metabolic pathways, |
| **Q8IR13** | FBgn0262734 | CG4429 | eIF4H1 | Eukaryotic translation initiation factor 4H1<br>(RNA-binding protein 2; isoform B) | GO:0003676~nucleic acid binding,<br>GO:0003723~RNA binding,<br>GO:0003729~mRNA binding,<br>GO:0003743~translation initiation factor activity,<br>GO:0033592~RNA strand annealing activity,<br>GO:0034057~RNA strand-exchange activity,<br>GO:0043024~ribosomal small subunit binding, | |
| **Q8IRQ5** | FBgn0286222 | CG4094 | Fum1 | Fumarate hydratase 1<br>(Fumarase 1)<br>(Lethal (1) G0255; isoform B) | GO:0003824~catalytic activity,<br>GO:0004333~fumarate hydratase activity,<br>GO:0016829~lyase activity, | dme00020:Citrate cycle (TCA cycle),<br>dme00620:Pyruvate metabolism,<br>dme01100:Metabolic pathways,<br>dme01200:Carbon metabolism, |
| **Q8MRK8** | FBgn0085479 | CG12136 | CG12136 | uncharacterized protein CG12136 (GH26280p) | | |
| **Q8MSI2** | FBgn0020908 | CG15848 | Scp1 | Sarcoplasmic calcium-binding protein 1 | GO:0005509~calcium ion binding, | |
| **Q8MST5** | FBgn0003890 | CG4869 | betaTub97EF | beta-Tubulin at 97EF<br>(LP10436p) | GO:0003924~GTPase activity,<br>GO:0005200~structural constituent of cytoskeleton,<br>GO:0005525~GTP binding, | dme04145:Phagosome,<br>dme04814:Motor proteins, |
| **Q8SY61** | FBgn0034470 | CG11218 | Obp56d | General odorant-binding protein 56d | GO:0005549~odorant binding, | |
| **Q8SY69** | FBgn0069923 | CG41128 | Mic10b | MICOS complex subunit MIC10<br>(RH61753p) | | |
| **Q8T8R1** | FBgn0034802 | CG3800 | CNBP | CCHC-type zinc finger nucleic acid binding protein | GO:0003676~nucleic acid binding,<br>GO:0003727~single-stranded RNA binding,<br>GO:0003729~mRNA binding,<br>GO:0008270~zinc ion binding,<br>GO:0045182~translation regulator activity,<br>GO:1905538~polysome binding, | |
| **Q94511** | FBgn0017566 | CG2286 | ND-75 | NADH dehydrogenase (ubiquinone) 75 kDa subunit | GO:0008137~NADH dehydrogenase (ubiquinone) activity,<br>GO:0016491~oxidoreductase activity,<br>GO:0016651~oxidoreductase activity, acting on NAD(P)H,<br>GO:0046872~metal ion binding,<br>GO:0051536~iron-sulfur cluster binding,<br>GO:0051537~2 iron, 2 sulfur cluster binding,<br>GO:0051539~4 iron, 4 sulfur cluster binding, | dme00190:Oxidative phosphorylation,<br>dme01100:Metabolic pathways, |

| | | | | | | |
|---|---|---|---|---|---|---|
| **Q94514** | FBgn0019624 | CG14724 | COX5A | Cytochrome c oxidase subunit 5A; mitochondrial | GO:0004129~cytochrome-c oxidase activity,<br>GO:0016491~oxidoreductase activity,<br>GO:0046872~metal ion binding, | dme00190:Oxidative phosphorylation,<br>dme01100:Metabolic pathways, |
| **Q94516** | FBgn0019644 | CG8189 | ATPsynB | ATP synthase subunit b, mitochondrial | GO:0015078~hydrogen ion transmembrane transporter activity,<br>GO:0046933~proton-transporting ATP synthase activity, rotational | dme00190:Oxidative phosphorylation,<br>dme01100:Metabolic pathways, |
| **Q94522** | FBgn0004888 | CG1065 | Scsalpha1 | Succinyl-CoA ligase [ADP/GDP-forming] subunit alpha; mitochondrial<br>(Succinyl-coenzyme A synthetase alpha subunit 1) | GO:0000166~nucleotide binding,<br>GO:0003824~catalytic activity,<br>GO:0004774~succinate-CoA ligase activity,<br>GO:0004775~succinate-CoA ligase (ADP-forming) activity,<br>GO:0004776~succinate-CoA ligase (GDP-forming) activity,<br>GO:0004778~succinyl-CoA hydrolase activity, | dme00020:Citrate cycle (TCA cycle),<br>dme00640:Propanoate metabolism,<br>dme01100:Metabolic pathways,<br>dme01200:Carbon metabolism, |
| **Q960M4** | FBgn0038570 | CG7217 | Prx5 | Peroxiredoxin-5<br>(RE23139p) | GO:0008379~thioredoxin peroxidase activity,<br>GO:0016491~oxidoreductase activity, | dme04146:Peroxisome, |
| **Q9GNH8** | FBgn0001186 | CG3001 | Hex-A | Hexokinase A<br>(Phosphotransferase) | GO:0004340~glucokinase activity,<br>GO:0004396~hexokinase activity,<br>GO:0005524~ATP binding,<br>GO:0005536~glucose binding,<br>GO:0008865~fructokinase activity,<br>GO:0016773~phosphotransferase activity, alcohol group as acceptor,<br>GO:0019158~mannokinase activity, | dme00010:Glycolysis / Gluconeogenesis,<br>dme00051:Fructose and mannose metabolism,<br>dme00052:Galactose metabolism,<br>dme00500:Starch and sucrose metabolism,<br>dme00520:Amino sugar and nucleotide sugar metabolism,<br>dme01100:Metabolic pathways,<br>dme01200:Carbon metabolism,<br>dme01250:Biosynthesis of nucleotide sugars, |
| **Q9GU68** | FBgn0285952 | CG3186 | eEF5 | Eukaryotic translation initiation factor 5A | GO:0003723~RNA binding,<br>GO:0003746~translation elongation factor activity,<br>GO:0043022~ribosome binding, | |
| **Q9TVP3** | FBgn0027654 | CG2239 | jdp | J domain-containing protein | GO:0051082~unfolded protein binding,<br>GO:0051087~chaperone binding, | |
| **Q9V3P0** | FBgn0040309 | CG1633 | Prx2<br>(Jafrac1) | Peroxiredoxin 2<br>(Jafrac 1) | GO:0004601~peroxidase activity,<br>GO:0008379~thioredoxin peroxidase activity,<br>GO:0016209~antioxidant activity,<br>GO:0016491~oxidoreductase activity,<br>GO:0051920~peroxiredoxin activity, | dme04146:Peroxisome, |
| **Q9V3W0** | FBgn0086691 | CG15261 | UK114 | RutC family protein UK114 | GO:0019239~deaminase activity, | |
| **Q9V3W2** | FBgn0001989 | CG13240 | ND-B17 | NADH dehydrogenase (ubiquinone) B17 subunit<br>(Lethal (2) 35Di; isoform A) | GO:0003954~NADH dehydrogenase activity,<br>GO:0016491~oxidoreductase activity, | dme00190:Oxidative phosphorylation,<br>dme01100:Metabolic pathways, |
| **Q9V427** | FBgn0027108 | CG4590 | Inx2 | Innexin 2 | GO:0005243~gap junction channel activity, | |
| **Q9V496** | FBgn0087002 | CG11064 | apolpp | Apolipophorins | GO:0005102~receptor binding,<br>GO:0005319~lipid transporter activity,<br>GO:0005504~fatty acid binding,<br>GO:0008017~microtubule binding,<br>GO:0015293~symporter activity,<br>GO:0019841~retinol binding,<br>GO:0020037~heme binding,<br>GO:0046872~metal ion binding, | |
| **Q9V4E0** | FBgn0039909 | CG1970 | ND-49 | NADH dehydrogenase (ubiquinone) 49 kDa subunit<br>(LD47962p) | GO:0003954~NADH dehydrogenase activity,<br>GO:0008137~NADH dehydrogenase (ubiquinone) activity,<br>GO:0016651~oxidoreductase activity, acting on NAD(P)H,<br>GO:0048038~quinone binding,<br>GO:0051287~NAD binding, | dme00190:Oxidative phosphorylation,<br>dme01100:Metabolic pathways, |
| **Q9V6U9** | FBgn0033883 | CG16935 | Mecr | Mitochondrial trans-2-enoyl-CoA reductase | GO:0016491~oxidoreductase activity,<br>GO:0016631~enoyl-[acyl-carrier-protein] reductase activity (NAD(P)H),<br>GO:0019166~trans-2-enoyl-CoA reductase (NADPH) activity, | dme00061:Fatty acid biosynthesis,<br>dme00062:Fatty acid elongation,<br>dme01100:Metabolic pathways,<br>dme01212:Fatty acid metabolism, |
| **Q9V8M5** | FBgn0034390 | CG15093 | CG15093 | Probable 3-hydroxyisobutyrate dehydrogenase, mitochondrial | GO:0008442~3-hydroxyisobutyrate dehydrogenase activity,<br>GO:0016491~oxidoreductase activity,<br>GO:0050661~NADP binding,<br>GO:0051287~NAD binding, | dme00280:Valine, leucine and isoleucine degradation,<br>dme00410:beta-Alanine metabolism,<br>dme00640:Propanoate metabolism,<br>dme01100:Metabolic pathways,<br>dme01200:Carbon metabolism, |
| **Q9VAC1** | FBgn0039737 | CG7920 | CG7920 | uncharacterized protein CG7920<br>(GM14349p) | GO:0003986~acetyl-CoA hydrolase activity,<br>GO:0008410~CoA-transferase activity,<br>GO:0008775~acetate CoA-transferase activity, | |
| **Q9VAJ4** | FBgn0039678 | CG18111 | Obp99a | General odorant-binding protein 99a | GO:0005549~odorant binding, | |
| **Q9VAM6** | FBgn0062442 | CG1458 | Cisd2 | CDGSH iron-sulfur domain | GO:0046872~metal ion binding,<br>GO:0051537~2 iron, 2 sulfur cluster binding, | |
| **Q9VAN7** | FBgn0014869 | CG1721 | Pglym78 | Phosphoglyceromutase 78; isoform C | GO:0003824~catalytic activity,<br>GO:0004082~bisphosphoglycerate mutase activity,<br>GO:0004619~phosphoglycerate mutase activity,<br>GO:0016868~intramolecular phosphotransferase activity,<br>GO:0046538~2,3-bisphosphoglycerate-dependent phosphoglycerate mutase activity, | dme00010:Glycolysis / Gluconeogenesis,<br>dme00260:Glycine, serine and threonine metabolism,<br>dme01100:Metabolic pathways,<br>dme01200:Carbon metabolism,<br>dme01230:Biosynthesis of amino acids, |
| **Q9VB69** | FBgn0029155 | CG5889 | Men-b | Malic enzyme b | GO:0004470~malic enzyme activity,<br>GO:0004471~malate dehydrogenase (decarboxylating) (NAD+) activity,<br>GO:0004473~malate dehydrogenase (decarboxylating) (NADP+) activity,<br>GO:0046872~metal ion binding,<br>GO:0051287~NAD binding, | dme00620:Pyruvate metabolism,<br>dme01100:Metabolic pathways,<br>dme01200:Carbon metabolism, |
| **Q9VBP6** | FBgn0039349 | CG4685 | Ssadh | Succinic semialdehyde dehydrogenase; isoform E | GO:0004777~succinate-semialdehyde dehydrogenase (NAD+) activity,<br>GO:0009013~succinate-semialdehyde dehydrogenase [NAD(P)+] activity,<br>GO:0016491~oxidoreductase activity,<br>GO:0016620~oxidoreductase activity, acting on the aldehyde or oxo group of donors, NAD or NADP as acceptor, | dme00250:Alanine, aspartate and glutamate metabolism,<br>dme00650:Butanoate metabolism,<br>dme01100:Metabolic pathways, |
| **Q9VBU9** | FBgn0039300 | CG10423 | RpS27 | 40S ribosomal protein S27<br>(Ribosomal protein S27) | GO:0003723~RNA binding,<br>GO:0003735~structural constituent of ribosome,<br>GO:0046872~metal ion binding, | dme03010:Ribosome, |
| **Q9VCD9** | FBgn0039145 | CG6000 | CG6000 | uncharacterized protein CG6000; isoform A | | |
| **Q9VCI3** | FBgn0039114 | CG10374 | Lsd-1 | Lipid storage droplets surface-binding protein 1 | | |
| **Q9VD58** | FBgn0038922 | CG6439 | Idh3b | Isocitrate dehydrogenase 3b<br>(GH26270p) | GO:0004449~isocitrate dehydrogenase (NAD+) activity, | dme00020:Citrate cycle (TCA cycle),<br>dme01100:Metabolic pathways,<br>dme01200:Carbon metabolism,<br>dme01210:2-Oxocarboxylic acid metabolism,<br>dme01230:Biosynthesis of amino acids, |
| **Q9VEB1** | FBgn0262559 | CG7998 | Mdh2 | Malate dehydrogenase 2, mitochondrial<br>(RE60471p) | GO:0003824~catalytic activity,<br>GO:0016491~oxidoreductase activity,<br>GO:0016615~malate dehydrogenase activity,<br>GO:0016616~oxidoreductase activity, acting on the CH-OH group of donors, NAD or NADP as acceptor,<br>GO:0030060~L-malate dehydrogenase activity, | dme00020:Citrate cycle (TCA cycle),<br>dme00270:Cysteine and methionine metabolism,<br>dme00620:Pyruvate metabolism,<br>dme00630:Glyoxylate and dicarboxylate metabolism,<br>dme01100:Metabolic pathways,<br>dme01200:Carbon metabolism, |
| **Q9VEJ0** | FBgn0038519 | CG5826 | Prx3 | Peroxiredoxin 3<br>(Thioredoxin peroxidase 3) | GO:0008379~thioredoxin peroxidase activity,<br>GO:0016209~antioxidant activity,<br>GO:0016491~oxidoreductase activity,<br>GO:0051920~peroxiredoxin activity, | |
| **Q9VEN1** | FBgn0014141 | CG3937 | cher | Filamin A<br>(cheerio) | GO:0003779~actin binding,<br>GO:0051015~actin filament binding, | |
| **Q9VEX6** | FBgn0287225 | CG6815 | bor | Belphegor<br>(ATPase family AAA domain-containing protein 3A homolog) | GO:0005524~ATP binding,<br>GO:0016887~ATPase activity, | |
| **Q9VF24** | FBgn0265048 | CG31150 | cv-d | Crossveinless d<br>(GH05619p) | GO:0005319~lipid transporter activity,<br>GO:0036122~BMP binding,<br>GO:0043395~heparan sulfate proteoglycan binding, | |
| **Q9VF27** | FBgn0017567 | CG3944 | ND-23 | NADH dehydrogenase (ubiquinone) 23 kDa subunit<br>(NADH:ubiquinone reductase 23kD subunit) | GO:0003954~NADH dehydrogenase activity,<br>GO:0008137~NADH dehydrogenase (ubiquinone) activity,<br>GO:0016651~oxidoreductase activity, acting on NAD(P)H,<br>GO:0046872~metal ion binding,<br>GO:0051539~4 iron, 4 sulfur cluster binding, | dme00190:Oxidative phosphorylation,<br>dme01100:Metabolic pathways, |
| **Q9VFC8** | FBgn0266064 | CG6904 | Glys | Glycogen synthase | GO:0004373~glycogen (starch) synthase activity, | dme00500:Starch and sucrose metabolism,<br>dme01100:Metabolic pathways, |
| **Q9VFF0** | FBgn0038271 | CG3731 | UQCR-C1 | Ubiquinol-cytochrome c reductase core protein 1<br>(GH01077p) | GO:0004222~metalloendopeptidase activity,<br>GO:0046872~metal ion binding, | |
| **Q9VFR2** | FBgn0038181 | CG9297 | CG9297 | uncharacterized protein CG9297<br>(FI18807p1) | | |
| **Q9VG32** | FBgn0002719 | CG10120 | Men | Malic enzyme | GO:0004470~malic enzyme activity,<br>GO:0004471~malate dehydrogenase (decarboxylating) (NAD+) activity,<br>GO:0004473~malate dehydrogenase (decarboxylating) (NADP+) activity,<br>GO:0046872~metal ion binding,<br>GO:0051287~NAD binding, | dme00620:Pyruvate metabolism,<br>dme01100:Metabolic pathways,<br>dme01200:Carbon metabolism, |
| **Q9VG82** | FBgn0038037 | CG11466 | Cyp9f2 | Probable cytochrome P450 9f2 | GO:0004497~monooxygenase activity,<br>GO:0005506~iron ion binding,<br>GO:0016705~oxidoreductase activity, acting on paired donors, with incorporation or reduction of molecular oxygen,<br>GO:0020037~heme binding, | |

| | | | | | | |
|---|---|---|---|---|---|---|
| **Q9VGS2** | FBgn0037874 | CG4800 | Tctp | Translationally-controlled tumor protein homolog | GO:0005085~guanyl-nucleotide exchange factor activity,<br>GO:0005509~calcium ion binding,<br>GO:0030295~protein kinase activator activity, | |
| **Q9VHJ8** | FBgn0037643 | CG11963 | ScsbetaA | Succinyl-coenzyme A synthetase β subunit, ADP-forming<br>(Succinate-CoA ligase [ADP-forming] subunit beta; mitochondria)<br>(SkpA associated protein; isoform B) | GO:0000287~magnesium ion binding,<br>GO:0004775~succinate-CoA ligase (ADP-forming) activity,<br>GO:0004776~succinate-CoA ligase (GDP-forming) activity,<br>GO:0005524~ATP binding,GO:0046872~metal ion binding, | dme00020:Citrate cycle (TCA cycle),<br>dme00640:Propanoate metabolism,<br>dme01100:Metabolic pathways,<br>dme01200:Carbon metabolism, |
| **Q9VHX4** | FBgn0037537 | CG2767 | CG2767 | uncharacterized protein CG2767<br>(LD24679p) | GO:0004032~alditol:NADP+ 1-oxidoreductase activity,<br>GO:0047834~D-threo-aldose 1-dehydrogenase activity,<br>GO:0016491~oxidoreductase activity, | dme00010:Glycolysis / Gluconeogenesis,<br>dme00040:Pentose and glucuronate interconversions,<br>dme00053:Ascorbate and aldarate metabolism,<br>dme00561:Glycerolipid metabolism,<br>dme00620:Pyruvate metabolism,<br>dme01100:Metabolic pathways,<br>dme01240:Biosynthesis of cofactors, |
| **Q9VIE8** | FBgn0010100 | CG9244 | mAcon1 | Mitochondrial aconitase 1 | GO:0003994~aconitate hydratase activity,<br>GO:0046872~metal ion binding,<br>GO:0047780~citrate dehydratase activity,<br>GO:0051539~4 iron, 4 sulfur cluster binding, | dme00020:Citrate cycle (TCA cycle),<br>dme00630:Glyoxylate and dicarboxylate metabolism,<br>dme01100:Metabolic pathways,<br>dme01200:Carbon metabolism,<br>dme01210:2-Oxocarboxylic acid metabolism,<br>dme01230:Biosynthesis of amino acids, |
| **Q9VIQ8** | FBgn0032833 | CG10664 | COX4 | Cytochrome c oxidase subunit 4<br>(MIP33903p1) | GO:0004129~cytochrome-c oxidase activity, | dme00190:Oxidative phosphorylation,<br>dme01100:Metabolic pathways, |
| **Q9VJZ4** | FBgn0032511 | CG9306 | ND-B22 | NADH dehydrogenase (ubiquinone) B22 subunit | GO:0016491~oxidoreductase activity, | dme00190:Oxidative phosphorylation,<br>dme01100:Metabolic pathways, |
| **Q9VK60** | FBgn0032453 | CG6180 | CG6180 | uncharacterized protein CG6180<br>(GH25425p) | | |
| **Q9VK99** | FBgn0032422 | CG6579 | atilla | Atilla, isoform B | GO:0034235~GPI anchor bingding, | |
| **Q9VL70** | FBgn0040064 | CG4600 | yip2 | Yippee interacting protein 2<br>(Acetyl-CoA acyltransferase) | GO:0003985~acetyl-CoA C-acetyltransferase activity,<br>GO:0003988~acetyl-CoA C-acyltransferase activity,<br>GO:0016746~transferase activity, transferring acyl groups,<br>GO:0016747~acyltransferase activity, transferring groups other than amino-acyl groups, | dme00062:Fatty acid elongation,<br>dme00071:Fatty acid degradation,<br>dme00280:Valine, leucine and isoleucine degradation,<br>dme01100:Metabolic pathways,<br>dme01212:Fatty acid metabolism, |
| **Q9VLB7** | FBgn0004868 | CG4422 | Gdi | GDP dissociation inhibitor<br>(LP03430p) | GO:0005092~GDP-dissociation inhibitor activity,<br>GO:0005093~Rab GDP-dissociation inhibitor activity,<br>GO:0005096~GTPase activator activity, | |
| **Q9VLP2** | FBgn0032021 | CG7781 | CG7781 | uncharacterized protein CG7781<br>(RE71014p) | GO:0034235~GPI anchor binding, | |
| **Q9VLZ3** | FBgn0040299 | CG6976 | Myo28B1 | Myosin 28B1; isoform A | GO:0000146~microfilament motor activity,<br>GO:0005524~ATP binding,<br>GO:0051015~actin filament binding, | dme04814:Motor proteins, |
| **Q9VMI3** | FBgn0031771 | CG9140 | ND-51 | NADH dehydrogenase (ubiquinone) 51 kDa subunit<br>(GM14163p) | GO:0008137~NADH dehydrogenase (ubiquinone) activity,<br>GO:0010181~FMN binding,<br>GO:0046872~metal ion binding,<br>GO:0051287~NAD binding,<br>GO:0051539~4 iron, 4 sulfur cluster binding, | dme00190:Oxidative phosphorylation,<br>dme01100:Metabolic pathways, |
| **Q9VMQ9** | FBgn0000318 | CG11024 | cl | clot<br>(Thioredoxin domain-containing protein 17)<br>(RE06767p) | GO:0004601~peroxidase activity,<br>GO:0047134~protein-disulfide reductase activity, | |
| **Q9VMS1** | FBgn0015031 | CG14028 | cype | Cytochrome c oxidase subunit Vic<br>(cyclope) | GO:0016491~oxidoreductase activity, | dme00190:Oxidative phosphorylation,<br>dme01100:Metabolic pathways, |
| **Q9VMT2** | FBgn0031692 | CG6514 | TpnC25D | Troponin C at 25D | GO:0005509~calcium ion binding,<br>GO:0030234~enzyme regulator activity, | |
| **Q9VMU0** | FBgn0031684 | CG8680 | ND-13A | NADH dehydrogenase (ubiquinone) 13 kDa A subunit<br>(LP20380p) | | dme00190:Oxidative phosphorylation,<br>dme01100:Metabolic pathways, |
| **Q9VN13** | FBgn0037239 | CG11739 | Sfxn1-3 | Sideroflexin 1/3<br>(RH48017p) | GO:0015075~monoatomic ion transmembrane transporter activity,<br>GO:0015194~L-serine transmembrane transporter activity,<br>GO:0022857~transmembrane transporter activity, | |
| **Q9VNB9** | FBgn0037328 | CG2099 | RpL35A | Ribosomal protein L35A | GO:0003735~structural constituent of ribosome, | dme03010:Ribosome, |
| **Q9VNE9** | FBgn0037351 | CG1475 | RpL13A | 60S ribosomal protein L13a,<br>(Ribosomal protein L13A) | GO:0003729~mRNA binding,<br>GO:0003735~structural constituent of ribosome, | dme03010:Ribosome, |
| **Q9VNW6** | FBgn0037146 | CG7470 | P5CS | Delta-1-pyrroline-5-carboxylate synthase<br>(GH12632p) | GO:0003824~catalytic activity,<br>GO:0004349~glutamate 5-kinase activity,<br>GO:0004350~glutamate-5-semialdehyde dehydrogenase activity,<br>GO:0005524~ATP binding,<br>GO:0016301~kinase activity,<br>GO:0016491~oxidoreductase activity,<br>GO:0016620~oxidoreductase activity, acting on the aldehyde or oxo group of donors, NAD or NADP as acceptor | dme00330:Arginine and proline metabolism,<br>dme01100:Metabolic pathways,<br>dme01230:Biosynthesis of amino acids, |
| **Q9VNX4** | FBgn0037138 | CG7145 | P5CDh1 | Delta-1-Pyrroline-5-carboxylate dehydrogenase 1<br>(Multifunctional fusion protein)<br>(FI08816p) | GO:0003842~1-pyrroline-5-carboxylate dehydrogenase activity,<br>GO:0016491~oxidoreductase activity,<br>GO:0016620~oxidoreductase activity, acting on the aldehyde or oxo group of donors, NAD or NADP as acceptor, | dme00250:Alanine, aspartate and glutamate metabolism,<br>dme00330:Arginine and proline metabolism,<br>dme01100:Metabolic pathways, |
| **Q9VP61** | FBgn0012034 | CG9390 | AcCoAS | Acetyl-coenzyme A synthetase | GO:0003987~acetate-CoA ligase activity,<br>GO:0005524~ATP binding,<br>GO:0016208~AMP binding,<br>GO:0031955~short-chain fatty acid-CoA ligase activity, | dme00010:Glycolysis / Gluconeogenesis,<br>dme00620:Pyruvate metabolism,<br>dme00630:Glyoxylate and dicarboxylate metabolism,<br>dme00640:Propanoate metabolism,<br>dme01100:Metabolic pathways,<br>dme01200:Carbon metabolism, |
| **Q9VPE2** | FBgn0037001 | CG6020 | ND-39 | NADH dehydrogenase (ubiquinone) 39 kDa subunit<br>(NADH dehydrogenase [ubiquinone] 1 alpha subcomplex subunit 9, mitochondrial) | GO:0016491~oxidoreductase activity,<br>GO:0044877~macromolecular complex binding, | dme00190:Oxidative phosphorylation,<br>dme01100:Metabolic pathways, |
| **Q9VQ61** | FBgn0001125 | CG4233 | Got2 | Aspartate aminotransferase<br>(Glutamate oxaloacetate transaminase 2) | GO:0003824~catalytic activity,<br>GO:0004069~L-aspartate:2-oxoglutarate aminotransferase activity,<br>GO:0008483~transaminase activity,<br>GO:0030170~pyridoxal phosphate binding, | dme00220:Arginine biosynthesis,<br>dme00250:Alanine, aspartate and glutamate metabolism,<br>dme00270:Cysteine and methionine metabolism,<br>dme00330:Arginine and proline metabolism,<br>dme00350:Tyrosine metabolism,<br>dme00360:Phenylalanine metabolism,<br>dme00400:Phenylalanine, tyrosine and tryptophan biosynthesis,<br>dme01100:Metabolic pathways,<br>dme01200:Carbon metabolism,<br>dme01210:2-Oxocarboxylic acid metabolism,<br>dme01230:Biosynthesis of amino acids, |
| **Q9VQG4** | FBgn0019830 | CG3057 | colt | Congested-like trachea protein | GO:0015227~acyl carnitine transmembrane transporter activity,<br>GO:0022857~transmembrane transporter activity, | |
| **Q9VQL7** | FBgn0283427 | CG3523 | FASN1 | Fatty acid synthase 1; isoform A | GO:0004312~fatty acid synthase activity,<br>GO:0004313~[acyl-carrier-protein] S-acetyltransferase activity,<br>GO:0004315~3-oxoacyl-[acyl-carrier-protein] synthase activity,<br>GO:0004316~3-oxoacyl-[acyl-carrier-protein] reductase (NADPH) activity,<br>GO:0004317~3-hydroxypalmitoyl-[acyl-carrier-protein] dehydratase activity,<br>GO:0004320~oleoyl-[acyl-carrier-protein] hydrolase activity,<br>GO:0008659~(3R)-hydroxymyristoyl-[acyl-carrier-protein] dehydratase activity,<br>GO:0008693~3-hydroxydecanoyl-[acyl-carrier-protein] dehydratase activity,<br>GO:0016295~myristoyl-[acyl-carrier-protein] hydrolase activity,<br>GO:0016296~palmitoyl-[acyl-carrier-protein] hydrolase activity,<br>GO:0016491~oxidoreductase activity,<br>GO:0017171~serine hydrolase activity,<br>GO:0031177~phosphopantetheine binding,<br>GO:0047117~enoyl-[acyl-carrier-protein] reductase (NADPH, A-specific) activity,<br>GO:0047451~3-hydroxyoctanoyl-[acyl-carrier-protein] dehydratase activity,<br>[illegible] | dme00061:Fatty acid biosynthesis,<br>dme01100:Metabolic pathways,<br>dme01212:Fatty acid metabolism, |
| **Q9VQR2** | FBgn0021967 | CG8844 | ND-PDSW | NADH dehydrogenase (ubiquinone) PDSW subunit<br>(NADH dehydrogenase [ubiquinone] 1 beta subcomplex subunit 10)<br>(RE65754p) | GO:0016491~oxidoreductase activity, | dme00190:Oxidative phosphorylation,<br>dme01100:Metabolic pathways, |
| **Q9VQT8** | FBgn0031561 | CG16712 | IM33 | Immune induced molecule 33<br>(RH38008p) | GO:0004867~serine-type endopeptidase inhibitor activity, | |
| **Q9VS34** | FBgn0035753 | CG8615 | RpL18 | 60S ribosomal protein L18<br>(Ribosomal protein L18) | GO:0003723~RNA binding,<br>GO:0003735~structural constituent of ribosome, | dme03010:Ribosome, |
| **Q9VSA3** | FBgn0035811 | CG12262 | Mcad | Medium-chain specific acyl-CoA dehydrogenase; mitochondrial | GO:0003995~acyl-CoA dehydrogenase activity,<br>GO:0016627~oxidoreductase activity, acting on the CH-CH group of donors,<br>GO:0050660~flavin adenine dinucleotide binding,<br>GO:0070991~medium-chain-acyl-CoA dehydrogenase activity, | dme00071:Fatty acid degradation,<br>dme00280:Valine, leucine and isoleucine degradation,<br>dme01100:Metabolic pathways,<br>dme01212:Fatty acid metabolism, |
| **Q9VSA9** | FBgn0035817 | CG7409 | CG7409 | uncharacterized protein CG7409 (RH03891p) | GO:0051082~unfolded protein binding, | |
| **Q9VSU6** | FBgn0035964 | CG4665 | Dhpr | Dihydropteridine reductase<br>(RE58329p) | GO:0004155~6,7-dihydropteridine reductase activity,<br>GO:0016491~oxidoreductase activity,<br>GO:0070402~NADPH binding,<br>GO:0070404~NADH binding, | dme00790:Folate biosynthesis,<br>dme01100:Metabolic pathways, |

| | | | | | | |
|---|---|---|---|---|---|---|
| **Q9VTB4** | FBgn0047038 | CG6463 | ND-13B | NADH dehydrogenase (ubiquinone) 13 kDa B subunit<br>(RH11203p) | GO:0008137~NADH dehydrogenase (ubiquinone) activity, | dme00190:Oxidative phosphorylation,<br>dme01100:Metabolic pathways, |
| **Q9VTK9** | FBgn0086254 | CG6084 | Akr1B | Aldo-keto reductase 1B<br>(LD06393p) | GO:0004032~alditol:NADP+ 1-oxidoreductase activity,<br>GO:0016491~oxidoreductase activity,<br>GO:0047718~indanol dehydrogenase activity,<br>GO:0047834~D-threo-aldose 1-dehydrogenase activity, | dme00040:Pentose and glucuronate interconversions,<br>dme00051:Fructose and mannose metabolism,<br>dme00052:Galactose metabolism,<br>dme00561:Glycerolipid metabolism,<br>dme00790:Folate biosynthesis,<br>dme01100:Metabolic pathways, |
| **Q9VTY2** | FBgn0036290 | CG10638 | CG10638 | uncharacterized protein CG10638; isoform A | GO:0004032~alditol:NADP+ 1-oxidoreductase activity,<br>GO:0047834~D-threo-aldose 1-dehydrogenase activity,<br>GO:0016491~oxidoreductase activity, | dme00040:Pentose and glucuronate interconversions,<br>dme00051:Fructose and mannose metabolism,<br>dme00052:Galactose metabolism,<br>dme00561:Glycerolipid metabolism,<br>dme00790:Folate biosynthesis,<br>dme01100:Metabolic pathways, |
| **Q9VU39** | FBgn0036337 | CG11255 | Adk2<br>(AdenoK) | Adenosine kinase<br>(GH14845p) | GO:0004001~adenosine kinase activity,<br>GO:0004017~adenylate kinase activity,<br>GO:0005524~ATP binding,<br>GO:0016301~kinase activity, | dme00230:Purine metabolism,<br>dme01100:Metabolic pathways,<br>dme01232:Nucleotide metabolism, |
| **Q9VU68** | FBgn0260049 | CG10724 | flr | flare<br>(Actin-interacting protein 1) | GO:0051015~actin filament binding, | |
| **Q9VUY9** | FBgn0003076 | CG5165 | Pgm1 | Phosphoglucomutase<br>(Phosphoglucose mutase 1) | GO:0000287~magnesium ion binding,<br>GO:0004614~phosphoglucomutase activity,<br>GO:0016868~intramolecular phosphotransferase activity, | dme00010:Glycolysis / Gluconeogenesis,<br>dme00030:Pentose phosphate pathway,<br>dme00052:Galactose metabolism,<br>dme00230:Purine metabolism,<br>dme00500:Starch and sucrose metabolism,<br>dme00520:Amino sugar and nucleotide sugar metabolism,<br>dme01100:Metabolic pathways,<br>dme01250:Biosynthesis of nucleotide sugars, |
| **Q9VV75** | FBgn0250814 | CG4169 | UQCR-C2 | Ubiquinol-cytochrome c reductase core protein 2 | GO:0046872~metal ion binding, | dme00190:Oxidative phosphorylation,<br>dme01100:Metabolic pathways, |
| **Q9VVH3** | FBgn0036726 | CG7603 | QIL1 | MICOS complex subunit MIC13 homolog QIL1<br>(RE63168p) | | |
| **Q9VVL7** | FBgn0289579 | CG7430 | Dld<br>(E3) | Dihydrolipoyl dehydrogenase | GO:0004148~dihydrolipoyl dehydrogenase activity,<br>GO:0050660~flavin adenine dinucleotide binding, | dme00010:Glycolysis / Gluconeogenesis,<br>dme00020:Citrate cycle (TCA cycle),<br>dme00260:Glycine, serine and threonine metabolism,<br>dme00280:Valine, leucine and isoleucine degradation,<br>dme00310:Lysine degradation,<br>dme00380:Tryptophan metabolism,<br>dme00620:Pyruvate metabolism,<br>dme00630:Glyoxylate and dicarboxylate metabolism,<br>dme00640:Propanoate metabolism,<br>dme00785:Lipoic acid metabolism,<br>dme01100:Metabolic pathways,<br>dme01200:Carbon metabolism,<br>dme01210:2-Oxocarboxylic acid metabolism,<br>dme01240:Biosynthesis of cofactors, |
| **Q9VVU1** | FBgn0036824 | CG3902 | GH07925p | uncharacterized protein CG3902<br>(Short/branched chain specific acyl-CoA dehydrogenase, mitochondrial)<br>(FI09602p) | GO:0003853~2-methylbutanoyl-CoA dehydrogenase activity,<br>GO:0003995~acyl-CoA dehydrogenase activity,<br>GO:0016627~oxidoreductase activity, acting on the CH-CH group of donors,<br>GO:0016937~short-chain fatty acyl-CoA dehydrogenase activity,<br>GO:0050660~flavin adenine dinucleotide binding, | dme00071:Fatty acid degradation,<br>dme00280:Valine, leucine and isoleucine degradation,<br>dme01100:Metabolic pathways,<br>dme01212:Fatty acid metabolism, |
| **Q9VW68** | FBgn0036927 | CG7433 | Gabat | Gamma-aminobutyric acid transaminase; isoform C<br>((S)-3-amino-2-methylpropionate transaminase) | GO:0003867~4-aminobutyrate transaminase activity,<br>GO:0008483~transaminase activity,<br>GO:0030170~pyridoxal phosphate binding,<br>GO:0034386~4-aminobutyrate:2-oxoglutarate transaminase activity,<br>GO:0047298~(S)-3-amino-2-methylpropionate transaminase activity, | dme00250:Alanine, aspartate and glutamate metabolism,<br>dme00280:Valine, leucine and isoleucine degradation,<br>dme00410:beta-Alanine metabolism,<br>dme00640:Propanoate metabolism,<br>dme00650:Butanoate metabolism,<br>dme01100:Metabolic pathways, |
| **Q9VWH4** | FBgn0027291 | CG12233 | Idh3a | Probable isocitrate dehydrogenase [NAD] subunit alpha, mitochondria<br>(Isocitrate dehydrogenase (NAD+) 3 catalytic subunit alpha) | GO:0000287~magnesium ion binding,<br>GO:0004449~isocitrate dehydrogenase (NAD+) activity,<br>GO:0016616~oxidoreductase activity, acting on the CH-OH group of donors, NAD or NADP as acceptor,<br>GO:0051287~NAD binding, | dme00020:Citrate cycle (TCA cycle),<br>dme01100:Metabolic pathways,<br>dme01200:Carbon metabolism,<br>dme01210:2-Oxocarboxylic acid metabolism,<br>dme01230:Biosynthesis of amino acids, |
| **Q9VWV6** | FBgn0022355 | CG6186 | Tsf1 | Transferrin 1 | GO:0005506~iron ion binding,<br>GO:0046872~metal ion binding, | |
| **Q9VX36** | FBgn0030853 | CG5703 | ND-24 | NADH dehydrogenase (ubiquinone) 24 kDa subunit | GO:0003954~NADH dehydrogenase activity,<br>GO:0008137~NADH dehydrogenase (ubiquinone) activity,<br>GO:0016491~oxidoreductase activity,<br>GO:0046872~metal ion binding,<br>GO:0051537~2 iron, 2 sulfur cluster binding, | dme00190:Oxidative phosphorylation,<br>dme01100:Metabolic pathways, |
| **Q9VXI6** | FBgn0030733 | CG3560 | UQCR-14 | Cytochrome b-c1 complex subunit 7<br>(Ubiquinol-cytochrome c reductase 14 kDa subunit)<br>(RH44664p) | | dme00190:Oxidative phosphorylation,<br>dme01100:Metabolic pathways, |
| **Q9VY93** | FBgn0030518 | CG11134 | CG11134 | Probable methylthioribulose-1-phosphate dehydratase | GO:0008270~zinc ion binding,<br>GO:0046570~methylthioribulose 1-phosphate dehydratase activity,<br>GO:0046872~metal ion binding, | dme00270:Cysteine and methionine metabolism,<br>dme01100:Metabolic pathways, |
| **Q9VYR1** | FBgn0030362 | CG1803 | regucalcin | Regucalcin<br>(SD03837p) | GO:0004341~gluconolactonase activity,<br>GO:0005509~calcium ion binding,<br>GO:0017171~serine hydrolase activity, | |
| **Q9VYU9** | FBgn0030332 | CG9360 | CG9360 | uncharacterized protein CG9360<br>(RH17287p) | GO:0000253~3-keto sterol reductase activity,<br>GO:0004303~estradiol 17-beta-dehydrogenase activity,<br>GO:0072555~17-beta-ketosteroid reductase activity,<br>GO:0072582~17-beta-hydroxysteroid dehydrogenase (NADP+) activity, | dme00900:Terpenoid backbone biosynthesis,<br>dme00981:Insect hormone biosynthesis, |
| **Q9VZ01** | FBgn0030292 | CG11752 | NdufV3 | NADH:ubiquinone oxidoreductase subunit V3<br>(RH44935p) | | |
| **Q9VZF6** | FBgn0035515 | CG14997 | Sqor | Sulfide quinone oxidoreductase<br>(GH04863p) | GO:0016491~oxidoreductase activity,<br>GO:0070224~sulfide:quinone oxidoreductase activity,<br>GO:0071949~FAD binding, | dme00920:Sulfur metabolism,<br>dme01100:Metabolic pathways, |
| **Q9VZR2** | FBgn0035434 | CG10812 | Drsl5 | Drosomycin-like 5 | | dme04624:Toll and Imd signaling pathway, |
| **Q9VZU4** | FBgn0266582 | CG12079 | ND-30<br>(NDUFS3) | NADH dehydrogenase (ubiquinone) 30 kDa subunit<br>(NADH dehydrogenase [ubiquinone] iron-sulfur protein 3, mitochondrial)<br>(RH59487p) | GO:0005515~protein binding,<br>GO:0008137~NADH dehydrogenase (ubiquinone) activity,<br>GO:0016651~oxidoreductase activity, acting on NAD(P)H, | dme00190:Oxidative phosphorylation,<br>dme01100:Metabolic pathways, |
| **Q9W0P5** | FBgn0035147 | CG12030 | Gale | UDP-glucose 4-epimerase<br>(UDP-galactose 4'-epimerase) | GO:0003974~UDP-N-acetylglucosamine 4-epimerase activity,<br>GO:0003978~UDP-glucose 4-epimerase activity, | dme00052:Galactose metabolism,<br>dme00520:Amino sugar and nucleotide sugar metabolism,<br>dme01100:Metabolic pathways,<br>dme01250:Biosynthesis of nucleotide sugars, |
| **Q9W125** | FBgn0035046 | CG3683 | ND-19<br>(NDUFA8) | NADH dehydrogenase (ubiquinone) 19 kDa subunit<br>(NADH dehydrogenase [ubiquinone] 1 alpha subcomplex subunit 8)<br>(RE35078p) | GO:0016491~oxidoreductase activity, | dme00190:Oxidative phosphorylation,<br>dme01100:Metabolic pathways, |
| **Q9W1C9** | FBgn0011695 | CG11390 | EbpIII | Ejaculatory bulb-specific protein 3 | | |
| **Q9W303** | FBgn0026415 | CG1780 | Idgf4 | Imaginal disc growth factor 4<br>(Chitinase-like protein Idgf4) | GO:0004568~chitinase activity,<br>GO:0008061~chitin binding,<br>GO:0008084~imaginal disc growth factor receptor binding, | |
| **Q9W334** | FBgn0030136 | CG2998 | RpS28b | Ribosomal protein S28b<br>(40S ribosomal protein S28) | GO:0003735~structural constituent of ribosome, | dme03010:Ribosome, |
| **Q9W369** | FBgn0086367 | CG12120 | t | tan<br>(Beta-alanyl-dopamine/carcinine hydrolase) | GO:0003677~DNA binding,<br>GO:0003693~P-element binding,<br>GO:0003832~beta-alanyl-dopamine hydrolase activity,<br>GO:0004803~transposase activity,<br>GO:0016740~transferase activity,<br>GO:0016787~hydrolase activity,<br>GO:0031964~beta-alanyl-histamine hydrolase activity,<br>GO:0046872~metal ion binding,<br>GO:0046983~protein dimerization activity, | |
| **Q9W3B3** | FBgn0030066 | CG1885 | Uros1 | Uroporphyrinogen-III synthase 1<br>(GH17465p) | GO:0004852~uroporphyrinogen-III synthase activity, | dme00860:Porphyrin metabolism,<br>dme01100:Metabolic pathways,<br>dme01240:Biosynthesis of cofactors, |
| **Q9W3L4** | FBgn0029990 | CG2233 | CG2233 | uncharacterized protein CG2233<br>(GH20802p) | | |
| **Q9W3R8** | FBgn0029942 | CG2059 | CG2059 | uncharacterized protein CG2059 | GO:0017171~serine hydrolase activity,<br>GO:0052689~carboxylic ester hydrolase activity,<br>GO:0080030~methyl indole-3-acetate esterase activity, | |

| | | | | | | |
|---|---|---|---|---|---|---|
| **Q9W401** | FBgn0261955 | CG3861 | Cs1<br>(kdn) | Citrate synthase 1<br>(knockdown)<br>(Probable citrate synthase, mitochondrial) | GO:0004108~citrate (Si)-synthase activity,<br>GO:0046912~transferase activity, transferring acyl groups, acyl groups converted into alkyl on transfer, | dme00020:Citrate cycle (TCA cycle),<br>dme00630:Glyoxylate and dicarboxylate metabolism,<br>dme01100:Metabolic pathways,<br>dme01200:Carbon metabolism,<br>dme01210:2-Oxocarboxylic acid metabolism,<br>dme01230:Biosynthesis of amino acids, |
| **Q9W4P5** | FBgn0285910 | CG2934 | VhaAC39-1 | V-type proton ATPase subunit d 1<br>(Vacuolar H[+] ATPase AC39 subunit 1) | GO:0046961~proton-transporting ATPase activity, rotational mechanism, | dme00190:Oxidative phosphorylation,<br>dme01100:Metabolic pathways,<br>dme04142:Lysosome,<br>dme04145:Phagosome, |
| **Q9W4Y3** | FBgn0284408 | CG33950 | trol | Terribly reduced optic lobes; isoform F | | |
| **Q9XTL9** | FBgn0004507 | CG7254 | Glyp | Glycogen phosphorylase | GO:0004645~1,4-alpha-oligoglucan phosphorylase activity,<br>GO:0008184~glycogen phosphorylase activity,<br>GO:0030170~pyridoxal phosphate binding,<br>GO:0042803~protein homodimerization activity, | dme00500:Starch and sucrose metabolism,<br>dme01100:Metabolic pathways, |
| **Q9XYN7** | FBgn0026380 | CG11981 | Prosbeta3 | Proteasome subunit beta type-3 | GO:0004175~endopeptidase activity,<br>GO:0004298~threonine-type endopeptidase activity, | dme03050:Proteasome, |
| **Q9Y112** | FBgn0027552 | CG10863 | CG10863 | uncharacterized protein CG10863 | GO:0004032~aldose reductase (NADPH) activity,<br>GO:0016491~oxidoreductase activity,<br>GO:0016616~oxidoreductase activity, acting on the CH-OH group of donors, NAD or NADP as acceptor,<br>GO:0047718~indanol dehydrogenase activity,<br>GO:0047834~D-threo-aldose 1-dehydrogenase activity, | dme00040:Pentose and glucuronate interconversions,<br>dme00051:Fructose and mannose metabolism,<br>dme00052:Galactose metabolism,<br>dme00561:Glycerolipid metabolism,<br>dme00790:Folate biosynthesis,<br>dme01100:Metabolic pathways, |

**Supplementary Table 4 | Oligonucleotide primers used for cloning and genotyping of the Atg4a-WT and Atg4a-C102S flies**
Lennicke *et al.*

| Primer | Sequence | Purpose |
|---|---|---|
| SOL897 | TATATAGGAAAGATATCCGGGTGAACTTCGATTATCGTATAGACGAGCAGGTTTTAGAGCTAGAAATAGCAAG | 5´ primer gRNA |
| SOL898 | ATTTTAACTTGCTATTTCTAGCTCTAAAACAATGTAAGTCGATCAGGGCCGACGTTAAATTGAAAATAGGTC | 3´ primer gRNA |
| SOL926 | ATGCGGCCGCAGAGCGAGCAGTGAGGGATAAG | 5´ primer donor construct + *NotI* site |
| SOL927 | TAGGTACCGTCCCTGCCGCTCTCTTCTGG | 3´ primer donor construct + *KpnI* site |
| SOL928 | CGCCCGTCACTGCCAACAAC | sequencing primer donor construct |
| SOL929 | CCACTCGCCGACCGCTTTG | sequencing primer donor construct + 3´ primer for FLAG genotyping |
| SOL930 | CCAGAACCCTTGTCATCGTCATC | sequencing primer donor construct |
| SOL931 | CCTGCAGAAGAAGCGATTGAAGAA | sequencing primer donor construct |
| SOL932 | CAACACCCTATCCCTTTTCAGAGC | sequencing primer donor construct |
| SOL933 | GGTATTGGATGACGTATGTAAGTGTGG | sequencing primer donor construct |
| SOL934 | CCAAGTTGAAGGAGGAGGTGCTC | sequencing primer donor construct |
| SOL935 | CGGCATCCTCGACCATTTCC | sequencing primer donor construct |
| SOL953 | GTCCACCCCCTTCTCCTTCTGC | 5´ primer to identify homozygous FLAG-ATG4a flies (179 bp on FLAG, 137 bp on WT) |
| SOL954 | GGCCAACTGCACTAGCGAGGAC | 3´ primer to identify homozygous FLAG-ATG4a flies (179 bp on FLAG, 137 bp on WT) |
| SOL955 | GGGTTCTGGAGGTTCTGGACTGG | 5´ primer FLAG genotyping |

**Supplementary Table 5 | QPCR primers**
Lennicke *et al.*

| Gene | GenBank Accession | Forward (5' - 3') | Reverse (5' - 3') | Reference |
|---|---|---|---|---|
| actin (Act5c) | NM_167053.2 | CACACCAAATCTTACAAAATGTGTGA | AATCCGGCCTTGCACATG | Bjedov *et al.* 2010 |
| Atg4a | NM_134719.4 | TTAACCCGCTTTACGTGCCT | CCTGCCGCTCTCTTCAACTA | This study |
| Atg5 | NM_132162.4 | GACATCCAACCGCTCTGCGCA | CAGACGATGACTTCACGTACACC | Bjedov *et al.* 2010 |
| Cat | NM_080483.3 | GAACTACTTTGCTGAGGTGGA | CATCTTGTCCGGAGAGGGCT | This study |
| tubulin (alphaTub84B) | NM_057424.4 | TGTCGCGTGTGAAACACTTC | AGCAGGCGTTTCCAATCTG | This study |
